# The arousal-motor hypothesis of dopamine function: evidence that dopamine facilitates reward seeking in part by maintaining arousal

**DOI:** 10.1101/2021.12.08.471650

**Authors:** Marcin Kaźmierczak, Saleem M. Nicola

**Affiliations:** Departments of Neuroscience and Psychiatry Albert Einstein College of Medicine 1300 Morris Park Ave Forchheimer 111 Bronx, NY 10461 USA

**Keywords:** Nucleus accumbens, arousal, reward-seeking behavior, caffeine, conditioned approach, sleep

## Abstract

Dopamine facilitates approach to reward via its actions on dopamine receptors in the nucleus accumbens. For example, blocking either D1 or D2 dopamine receptors in the accumbens reduces the proportion of reward-predictive cues to which rats respond with cued approach. Recent evidence indicates that accumbens dopamine also promotes wakefulness and arousal, but the relationship between dopamine’s roles in arousal and reward seeking remains unexplored. Here, we show that the ability of systemic or intra-accumbens injections of the D1 antagonist SCH23390 to reduce cued approach to reward depends on the animal’s state of arousal. Handling the animal, a manipulation known to increase arousal, was sufficient to reverse the behavioral effects of the antagonist. In addition, SCH23390 reduced spontaneous locomotion and increased time spent in sleep postures, both consistent with reduced arousal, but also increased time spent immobile in postures inconsistent with sleep. In contrast, the ability of the D2 antagonist haloperidol to reduce cued approach was not reversible by handling. Haloperidol reduced spontaneous locomotion but did not increase sleep postures, instead increasing immobility in non-sleep postures. We place these results in the context of the extensive literature on dopamine’s contributions to behavior, and propose the arousal-motor hypothesis. This novel synthesis, which proposes that two main functions of dopamine are to promote arousal and facilitate motor behavior, accounts both for our findings and many previous behavioral observations that have led to disparate and conflicting conclusions.

## Introduction

Dopamine is widely believed to be critical for learning about rewards and for enhancing performance aimed at obtaining them. Accordingly, the most prevalent hypotheses implicate dopamine in reward prediction error encoding and reinforcement on the one hand (Glimcher, 2011; Schultz, 2016), and motivation to pursue rewards and exert effort on the other (Salamone and Correa, 2012; Salamone et al., 2018). However, dopamine is also critical for motor function and arousal. Recent evidence shows that even the structures considered central for reinforcement and motivation, such as the ventral tegmental area (VTA) (Chowdhury et al., 2019; Eban-Rothschild et al., 2016; Oishi et al., 2017a; Takata et al., 2018; Taylor et al., 2016; Yu et al., 2019) and the nucleus accumbens (NAc) (Luo et al., 2018; Oishi et al., 2017b), are directly and critically involved in the regulation of wakefulness. These findings reintroduce conundrums that were thought to be settled decades ago: to what extent are the behavioral effects of dopamine manipulations that ostensibly alter reinforcement or motivation the result of more generalized performance deficits, such as diminished arousal? To address this question, we first review the early catecholamine behavioral literature and place recent arousal findings in this historical context. We also review influential historical arguments supporting the view that dopaminergic disruptions, under certain conditions, impair performance of appetitive operant tasks due to reward-specific effects, rather than generalized performance problems. We conclude that those arguments were erroneous because the effects of diminished arousal, and the interaction between arousal and motor deficits, were not considered. Next, we review more recent literature that establishes roles for mesolimbic dopamine, dopamine D1 receptors, and dopamine D2 receptors in arousal, and contrast these observations with studies in which the disruptive effects of manipulations of mesolimbic dopamine on operant performance were interpreted as evidence for impaired motivation or reinforcement. We then experimentally test the hypothesis that dopamine antagonist-induced deficits in operant performance are due to arousal and motor deficits rather than deficits in motivation or reinforcement. Finally, we integrate these results with the broader literature to propose a novel hypothesis of dopamine function, the arousal-motor hypothesis.

The views of the neurophysiological and behavioral functions of dopamine have been radically changing over the last sixty years. The importance of monoamine neurotransmitters for reward was first recognized in 1961 (Stein and Seifter, 1961); intracranial self-stimulation (ICSS) in rats was enhanced or inhibited by drugs that correspondingly increased or decreased brain levels of monoamines. Interestingly, it was not dopamine but norepinephrine whose importance for reward was initially emphasized; Wise and Stein (1969) showed that disulfiram, an inhibitor of norepinephrine synthesis, disrupts ICSS. Moreover, rats’ performance was fully restored by intraventricular administration of norepinephrine, but not dopamine or serotonin. The authors firmly concluded that the “rewarding effect of medial forebrain bundle stimulation may depend on the availability of norepinephrine as a transmitter, but not on dopamine or serotonin”. This interpretation was then questioned by Roll (1970), who noticed that while animals administered the norepinephrine synthesis inhibitor disulfiram indeed stopped pressing a bar for ICSS, they also appeared sleepy. When handled and put back on the bar, the animals resumed pressing at normal rates before succumbing to sleep again. Roll concluded that decreased performance resulted from the effects of norepinephrine depletion on wakefulness, rather than reward. Similar conclusions drawn by other groups ultimately led to the rejection of the “noradrenergic theory of reward” (Rolls et al., 1974a).

This early controversy unearthed a fundamental problem pertaining to all experiments involving interventions (e.g., pharmacological, lesion, optogenetic) that inhibit the pursuit of rewards: how can one know whether an intervention interfered with a reward-related function, such as reinforcement or motivation, or instead caused “performance problems” by making experimental animals sleepy, cataleptic, or unable to satisfy specific operant requirements? These concerns are particularly relevant to the study of dopaminergic systems because dopamine deficiency has long been implicated in the motor deficits observed in Parkinson’s patients (Hornykiewicz, 1972). Moreover, when selective lesions of dopaminergic pathways became possible in animals, they resulted in profound inhibition of almost all behaviors, including spontaneous movement (Ungerstedt, 1971). Some early studies argued that the inhibition of ICSS by neuroleptics, which mainly antagonize D2 receptors, is due to motor deficits, which can be subtle and not always manifested as a gross inhibition of movement (Fibiger et al., 1976; Rolls et al., 1974b). The hypothesis that dopamine mediates reinforcement began to gain momentum after an influential study in which Fouriezos and Wise (1976) noticed that rats administered the neuroleptic drug pimozide exhibited gradual within-session declines in ICSS responding. The authors argued that their pimozinde-administered rats did not have performance problems because they were capable of vigorous lever pressing at the beginning of the operant session. The rate of lever pressing declined only after the animals had experienced some rewards under the influence of the drug. They hypothesized that the rewards were no longer reinforcing when the drug was on board, and therefore the decline reflected extinction.

The interpretation of neuroleptic-induced within-session declines as a consequence of impaired reinforcement led to the formulation of the “dopamine theory of reward” (Wise, 1978), which provided a framework for interpretation of an astounding variety of behavioral experiments, including food-reinforced performance, conditioned place preference, drug self-administration, and animal models of neuropsychiatric illnesses. The argument against performance problems became so influential that many neuroscientists ceased to seriously consider the possibility that dopamine manipulations affect arousal or motor function and focused exclusively on “reward-specific” concepts such as liking, learning, and wanting. Considering the tremendous influence that the argument based on within-session declines of performance has had on the field, it is worthwhile to reassess its credibility. The idea that neuroleptic-induced gradual declines result from impaired reinforcement has been heavily criticized. First, neuroleptics markedly decrease lever pressing under extinction conditions when no rewards are consumed (Mason et al., 1980; Phillips and Fibiger, 1979). Second, while reward omission and administration of neuroleptics produce superficially similar patterns of responding on a fixed ratio schedule (Fouriezos and Wise, 1976), these conditions greatly differ in their sensitivity to the operant requirement. For example, neuroleptic doses that abolished lever pressing for ICSS were much less effective when rats were required to nose-poke for the same reward (Ettenberg et al., 1981). This result demonstrated that extinction was not at play and reinforcement was not impaired, but rather the drug compromised the ability to press a lever. Third, Fibiger (1976) reported that pimozide and haloperidol (HAL), another neurolpetic, produced a uniform decrease in responding throughout the session, rather than a gradual decline, when rats performed on a variable interval 60 s schedule (VI60). Moreover, when the interval was increased to 4 min, it was evident that the rate of lever pressing declined before any rewards were consumed (Phillips and Fibiger, 1979). These results are clearly incompatible with the extinction-like interpretation.

Further incompatible with a reinforcement deficit, neuroleptics block avoidance of aversive stimuli, such as electric shock (Sanger, 1986). Moreover, animals with disrupted dopaminergic function show signs of distress during the presentation of shock-predictive stimuli, such as urination and defecation (Fibiger et al., 1975; Lenard and Beer, 1975), yet they do not escape. These and other experiments on the effects of neuroleptics on conditioned avoidance (Beninger et al., 1980) strongly suggest that the drugs simply disrupt the initiation of a motor response. It is unlikely that the same drugs had completely different, non-motor effects on animals pursuing rewards. A more parsimonious explanation is that certain motor deficits, including difficulties with movement initiation, underlie both disrupted pursuit of rewards and shock avoidance.

To account for the published results described so far, the specific neuroleptic-induced performance deficit must explain within-session decline patterns of responding. In fact, there is clear evidence for progressive, reward-independent motor problems after neuroleptic administration. Liao and Fowler (1990) showed that the length of individual lever presses in HAL-treated rats performing on an FR20 schedule for sweetened milk gradually increases throughout the operant session. This phenomenon does not occur during extinction when no drugs are administered (Faustman and Fowler, 1981) and indicates a progressive slowing of movement in HAL-treated rats (Liao and Fowler, 1990). These results suggest that neuroleptics induce a progressive “fatigue-like” motor impairment, which could increasingly impair operant performance as the session progresses in time. Proponents of the reinforcement interpretation argued against this alternative by emphasizing that a mid-session disturbance, such as sensory stimulation or temporary restriction of the access to the operandum, can cause the animal to resume responding while still under the influence of the neuroleptic (Fouriezos and Wise, 1976; Franklin and McCoy, 1979; Wise, 2004). Indeed, if *physical* fatigue is responsible for within-session declines in performance, then it should not simply go away in response to such stimulation.

However, this argument does not mean that the reinforcement hypothesis must be correct. We propose another possibility that is compatible with the evidence discussed so far. Specifically, arousal declines naturally during behavior sessions in which the animal is asked to perform repetitive operant tasks. This decline in arousal can interfere with task performance depending on the specific task structure. We propose that dopamine disruption amplifies these performance deficits either by potentiating the natural reduction in arousal or enhancing the ability of a decline in arousal to reduce operant performance. In support of this hypothesis, it has long been known that motor deficits in Parkinson’s patients can be significantly influenced by arousal. While minor stressors and brief unexpected stimuli can occasionally aggravate the symptoms, potently arousing stimuli or those that convey motivational value can alleviate or even completely abolish motor deficits. For example, severely akinetic patients can transiently resume normal functioning in response to a fire alarm (Schwab and Zieper, 1965). This phenomenon, called “paradoxical kinesia”, has been replicated in dopamine-depleted rats. After treatment with the catecholamine-specific neurotoxin 6-hydroxydopamine, severely incapacitated animals that are unable to eat, drink, or move on their own can nevertheless rear or even jump out of the sink in response to cold water, run away from a cat, or actively interact and even mount with other rats. Such powerfully arousing stimuli transiently alleviate almost all investigated motor symptoms (Marshall et al., 1976). Paradoxical kinesia is highly relevant to the study of behavioral effects of neuroleptics because researchers studying animal behavior usually let the subjects initiate task performance soon after they are transferred to operant chambers. Being transferred from the home cage to an operant chamber is a highly arousing stimulus, which could alleviate neuroleptic-induced motor deficits (paradoxical kinesia). This means that the gradual decline in operant performance after neuroleptic administration is neither due to a reinforcement deficit nor a motivational deficit, nor is it the result of physical fatigue. Rather, it is simply due to declining arousal. That neuroleptic effects can be alleviated mid-session by delivering arousing stimuli supports this hypothesis.

Disruptions of dopamine could contribute to performance deficits in two ways. In states of low (but not high) arousal, dopamine may be required for certain operant behaviors. Within this interpretation, dopaminergic disruptions do not directly affect arousal but cause performance problems when arousal declines naturally. Alternatively, dopamine could be required to maintain a threshold level of arousal, such that animals with deficient dopamine transmission become too sleepy to perform unless an arousing stimulus triggers a compensatory increase in dopamine release or engages dopamine-independent arousal mechanisms. Supporting the latter hypothesis, it is well documented that neuroleptics promote sleep (Monti and Monti, 2007; Ongini et al., 1993; Ongini and Trampus, 1992; Trampus and Ongini, 1990), and genetic deletion of D2 receptors decreases wakefulness (Qu et al., 2010). Conversely, psychostimulant drugs such as amphetamines or modafinil are extremely effective in promoting wakefulness (Boutrel and Koob, 2004) due to their dopamine-enhancing activity (Qu et al., 2008; Wisor et al., 2001), and selective D1 and D2 receptor agonists promote electrocortical wakefulness and inhibit sleep (Isaac and Berridge, 2003). Activation of D1 receptors was even sufficient to elicit neurophysiological signs of arousal in anesthetized animals and precipitate reanimation from general anesthesia (Taylor et al., 2013). If animals administered dopamine antagonists are sleepy, then both a gradual within-session decline of performance and reactivation of performance by wake-promoting stimuli are precisely what one would expect.

Further supporting the arousal hypothesis, recent studies established that the ventral tegmental area plays a critical role in the regulation of arousal (Chowdhury et al., 2019; Eban-Rothschild et al., 2016; Oishi et al., 2017a; Takata et al., 2018; Taylor et al., 2016; Yu et al., 2019). Stimulation of VTA dopamine neurons was sufficient to induce emergence from general anesthesia (Taylor et al., 2016); optogenetic activation induced an almost instantaneous transition from non-REM sleep to wakefulness (Eban-Rothschild et al., 2016); chemogenetic *inhibition* induced episodes of sleep preceded by diligent nesting (Eban-Rothschild et al., 2016); and semi-chronic optogenetic *stimulation* potently inhibited both sleep and nest-building behavior (Eban-Rothschild et al., 2016). Conversely, activation and inhibition of VTA GABA neurons was found to promote sleep and wakefulness, respectively (Chowdhury et al., 2019; Yu et al., 2019), in part due to the impact of these manipulations on the activity of VTA dopamine neurons. Because the VTA is the major source of dopaminergic innervation of the NAc, these findings are consistent with multiple additional lines of evidence implicating the NAc in the regulation of arousal and sleep (Barik and de Beaurepaire, 2005; Lazarus et al., 2011; Lena et al., 2005; Luo et al., 2018; Oishi et al., 2017b; Qiu et al., 2012; Wang et al., 2019; Zhou et al., 2019). For example, the wake-promoting actions of caffeine (Lazarus et al., 2011) and modafinil (Qiu et al., 2012) crucially depend on their effects in the NAc. Furthermore, it has been recently reported that both D1 receptor-expressing and D2 receptor / adenosine 2A receptor-expressing subpopulations of NAc medium spiny neurons are critically involved in the control of sleep: chemogenetic or optogenetic activation/inhibition of D1-expressing neurons respectively counteracted/ promoted sleep (Luo et al., 2018), while bidirectional modulation of the activity of D2-expressing neurons had the opposite effects (Oishi et al., 2017b). Finally, both glutamatergic (Yu et al., 2019) and dopaminergic (Eban-Rothschild et al., 2016) projection from the VTA specifically to the NAc promotes wakefulness.

The evidence that mesolimbic dopamine contributes to arousal calls into question the interpretation of many previous findings concerning the sensitivity of certain reward-oriented tasks to NAc dopamine disruptions (reviewed in Nicola, 2007, 2016). The ability of these disruptions to impair performance is strongly correlated with the length of the inter-trial-interval, with greater effects when the intervals are longer (Nicola, 2007). Similarly, increasing the length of the intertrial interval (ITI) in a cued FR1 task while keeping all other parameters constant markedly increases task sensitivity to intra-NAc infusions of dopamine antagonists (Nicola, 2010). Long intervals promote movement away from the operandum, forcing the animals to respond to the cues with a flexible or “taxic” approach strategy. Therefore, we previously hypothesized that NAc dopamine is particularly important for taxic approach to the operandum that requires guidance by sensory cues (du Hoffmann and Nicola, 2016; Nicola, 2010). However, another aspect of long-interval tasks that may contribute to their vulnerability to NAc dopamine disruptions is that the rate of arousing events (such as cues, rewards and reward-oriented behaviors) is lower. Logically, the subject’s overall arousal level is likely to be lower during a task that includes long periods in which nothing happens than a task in which cues are presented rapidly and operant performance is consistently rewarded. Hence, long-interval tasks should be particularly susceptible to manipulations that decrease arousal. Importantly, in humans, the effects of sleepiness on task performance are usually negligible during the initial 10 min of the task and then gradually increase in severity (Donnell, 1969; Wilkinson, 1960, 1961) although instantaneous effects can also be observed under certain conditions (Gillberg and Akerstedt, 1998). Similarly, we have noticed that intra-NAc administration of the D1 antagonist SCH23390 (SCH) causes a delayed effect on responding to sucrose-predictive cues with a maximal effect after 20-30 min (Yun et al., 2004), in line with the possible involvement of drowsiness. However, some studies have implicated D1 receptors in reinforcement (Ikemoto et al., 1997; White et al., 1991; Wolterink et al., 1993), and therefore a gradual decline could instead reflect an extinction-like phenomenon. Finally, the delayed effect of the drug could be caused by slow pharmacodynamics: the drug may act at a distance, requiring time to diffuse from the infusion site (Caine et al., 1995); or a gradual attenuation of the intracellular effects of D1 receptor activity may occur (Undieh, 2010); or there may be some other delayed biological effect of the drug. Interestingly, while the gradual within-session declines caused by drugs antagonizing predominantly D2 receptors have attracted much attention, those induced by D1 antagonists have not been explored. Therefore, whereas the declines caused by typical antipsychotics are unlikely to represent impaired reinforcement, as discussed above, the possibility that D1 antagonists inhibit the pursuit of rewards by blocking reinforcement has not been ruled out.

To address the question of whether D1 receptors contribute to arousal or reinforcement in a cued long interval task, we explored the within-session temporal pattern of responding to reward-predictive cues in rats administered the D1 antagonist SCH. We observed pronounced gradual performance declines after either intra-NAc or systemic administrations of the drug. By establishing specific conditions under which these declines occur, we were able to conclude that they reflect declining arousal. For comparison, we explored the effects of the neuroleptic dug HAL on cue responding and discovered that similar gradual declines do not occur. We found that this difference is a consequence of a difference in the efficacy with which natural arousing stimuli alleviate the effects of SCH and HAL on cued pursuit of sucrose. We further compared the effects of natural arousing stimuli to those of caffeine (CAF) and discovered that CAF is effective in alleviating the effects of either dopamine antagonist. Finally, an analysis of video recordings provided support for the involvement of sleepiness and motor problems in the effects of the drugs on performance and no indication of reward-specific deficits. We conclude that decreased responsiveness to reward-predictive cues in rats administered SCH and HAL is likely due to a combination of drowsiness and motor deficits. Our study provides clear evidence that high rates of operant performance at the beginning of the session cannot be used as an argument to rule out performance deficits, and indicates that care must be taken to rule out performance and arousal effects when evaluating the putative effects of mesolimbic dopamine manipulations on reinforcement and motivation.

## Methods

### Animals

Male Long-Evans rats weighing 300-330 g were purchased from Charles River Laboratories. One week after arrival, animals were put on a restricted diet of 18 g of Bio-Serv formula F-173 rodent pellets per day, which was gradually increased to 24 g per day while the animals were gaining weight. The average weight of the animals at the time when pharmacology experiments were performed was about 600 g for cohort 1 and 450 g for cohort 2. This difference was due to longer training for the animals from cohort 1 (∼4 weeks), followed by surgeries and an additional two weeks of retraining. The animals from cohort 2 underwent shorter training (∼2 weeks) after which the experiments commenced immediately. Altogether, 32 animals were used, 16 for each cohort. Three animals from cohort 1 were lost during the surgeries and one additional animal became sick during the course of experiments. Therefore, the n for cohort 1 experiments, performed using a within-subject design, was either 13 or 12. Cohort 2 was separated into two subgroups of n = 8 animals each and the experiments were performed using a between-subject design to minimize the impact of multiple testing on our conclusions. Multiple separate experiments were conducted with the subjects in cohort 1, but only one experiment was conducted with cohort 2. Table 1 provides a list of all experiments performed with cohorts 1 and 2, as well as an index of figures that utilize the data from each experiment. The procedures were compliant with the guidelines from the National Institute of Health and were approved by the Institutional Animal Care and Use Committee of the Albert Einstein College of Medicine.

**Table 1.**
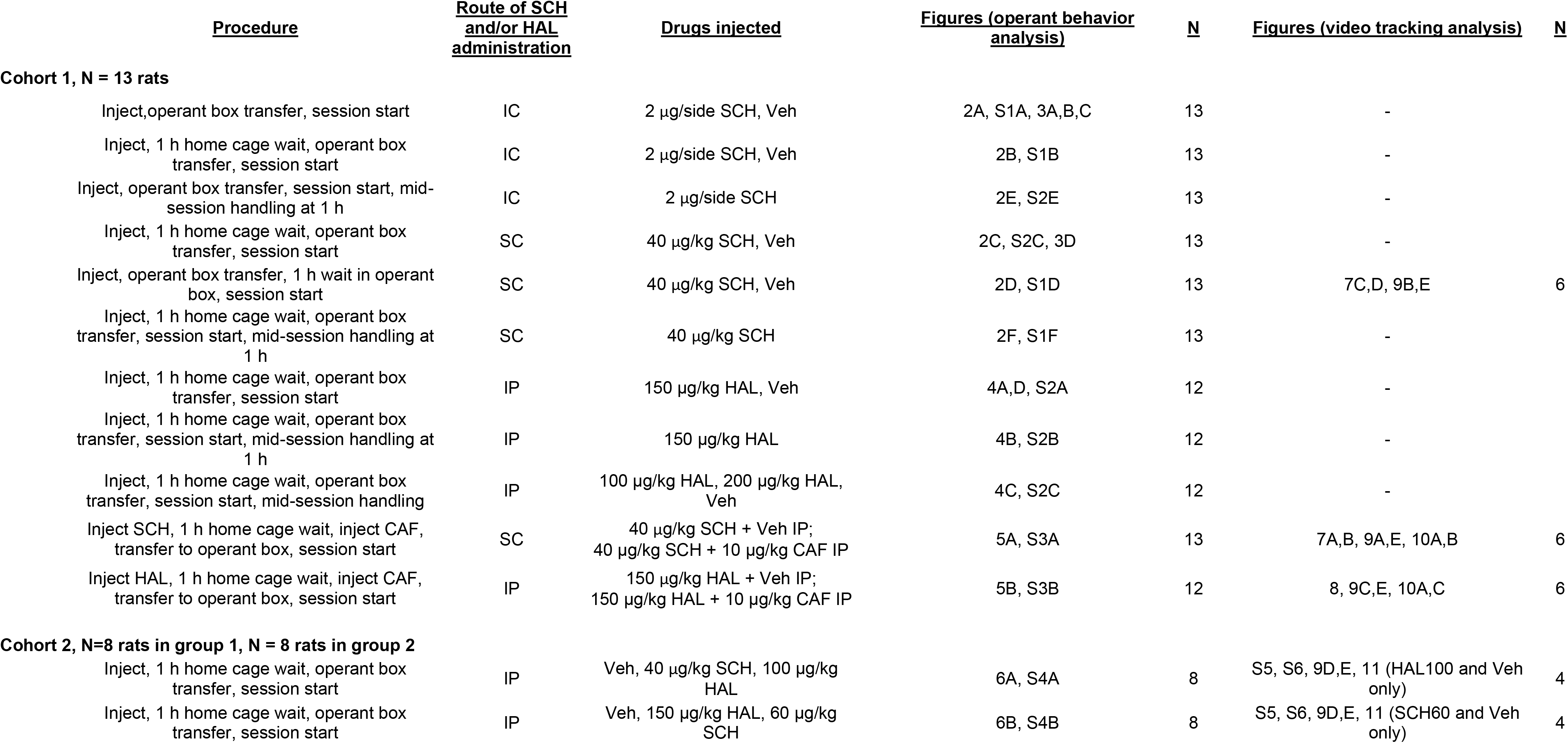
List of experiments performed and index of figures. Two cohorts of rats were used for these experiments, cohort 1 (N=13) and cohort 2 (divided into group 1 and group 2, N = 8 each). The experiments performed with each cohort or group are listed in individual rows; multiple experiments were performed with cohort 1. The “Procedures” column describes the events in chronological order, with no imposed intervals between events unless explicitly stated. Analysis of data that did not involve video tracking utilized all subjects in the cohort or group. Additional analyses of video tracking data were performed for subsets of animals in some experiments. The figures utilizing data from each experiment, and the N of rats used for the analyses, are listed in separate columns for operant behavior (not involving tracking) and video tracking analyses.

| <u>Procedure</u> | <u>Route of SCH</u><br><u>and/or HAL</u><br><u>administration</u> | <u>Drugs injected</u> | <u>Figures (operant behavior</u><br><u>analysis)</u> | <u>N</u> | <u>Figures (video tracking analysis)</u> | <u>N</u> |
| --- | --- | --- | --- | --- | --- | --- |
| **Cohort 1, N = 13 rats** |  |  |  |  |  |  |
| Inject, operant box transfer, session start | IC | 2 µg/side SCH, Veh | 2A, S1A, 3A,B,C | 13 | - |  |
| Inject, 1 h home cage wait, operant box transfer, session start | IC | 2 µg/side SCH, Veh | 2B, S1B | 13 | - |  |
| Inject, operant box transfer, session start, mid-session handling at 1 h | IC | 2 µg/side SCH | 2E, S2E | 13 | - |  |
| Inject, 1 h home cage wait, operant box transfer, session start | SC | 40 µg/kg SCH, Veh | 2C, S2C, 3D | 13 | - |  |
| Inject, operant box transfer, 1 h wait in operant box, session start | SC | 40 µg/kg SCH, Veh | 2D, S1D | 13 | 7C,D, 9B,E | 6 |
| Inject, 1 h home cage wait, operant box transfer, session start, mid-session handling at 1 h | SC | 40 µg/kg SCH | 2F, S1F | 13 | - |  |
| Inject, 1 h home cage wait, operant box transfer, session start | IP | 150 µg/kg HAL, Veh | 4A,D, S2A | 12 | - |  |
| Inject, 1 h home cage wait, operant box transfer, session start, mid-session handling at 1 h | IP | 150 µg/kg HAL | 4B, S2B | 12 | - |  |
| Inject, 1 h home cage wait, operant box transfer, session start, mid-session handling | IP | 100 µg/kg HAL, 200 µg/kg HAL, Veh | 4C, S2C | 12 | - |  |
| Inject SCH, 1 h home cage wait, inject CAF, transfer to operant box, session start | SC | 40 µg/kg SCH + Veh IP;<br>40 µg/kg SCH + 10 µg/kg CAF IP | 5A, S3A | 13 | 7A,B, 9A,E, 10A,B | 6 |
| Inject HAL, 1 h home cage wait, inject CAF, transfer to operant box, session start | IP | 150 µg/kg HAL + Veh IP;<br>150 µg/kg HAL + 10 µg/kg CAF IP | 5B, S3B | 12 | 8, 9C,E, 10A,C | 6 |
| **Cohort 2, N=8 rats in group 1, N = 8 rats in group 2** |  |  |  |  |  |  |
| Inject, 1 h home cage wait, operant box transfer, session start | IP | Veh, 40 µg/kg SCH, 100 µg/kg HAL | 6A, S4A | 8 | S5, S6, 9D,E, 11 (HAL100 and Veh only) | 4 |
| Inject, 1 h home cage wait, operant box transfer, session start | IP | Veh, 150 µg/kg HAL, 60 µg/kg SCH | 6B, S4B | 8 | S5, S6, 9D,E, 11 (SCH60 and Veh only) | 4 |

### Behavioral training

Behavioral experiments were performed in 30 x 25 cm Med Associates operant chambers enclosed in melamine cabinets. Two 28 V white house lights and white noise (65 dB) were on at all times when the rat was in the chamber. The chambers were equipped with a liquid reward receptacle through which 70 µl of 10% sucrose reward was delivered. A photobeam across the receptacle was used to detect both receptacle entries and exits. After one day of magazine training during which the reward was delivered upon each receptacle entry, animals were trained on FI10s, FI20s and FI30s on three consecutive days. The task was then switched to VI20-40s and a reward-predictive auditory cue was introduced (discriminative stimulus, DS) that informed animals about reward availability. The initial length of the cue was 20 s and receptacle entry during cue presentation resulted in reward delivery. After animals met the criterion of responding to more than 80% of the cues, a second distinct auditory cue was introduced that did not predict reward availability (neutral stimulus, NS). The two cues were a siren cue (which cycled in frequency from 4 to 8 kHz over 400 ms) and an intermittent cue (6 kHz that was on for 40 s, off for 50 ms). The assignment of these cues to DS and NS was counterbalanced across animals. DS and NS presentations were randomly interspersed throughout operant sessions. The lengths of cues were subsequently shortened to 10 s and the ITI was gradually increased to VI20-80s. The initial session length of 1 h was increased to 2 h after the introduction of the second stimulus. The training proceeded until the response ratio to the NS decreased to less than 20% (cohort 1, longer training) or 40% (cohort 2, shorter training), while the response ratio to the DS remained above 80%. Two blue cue lights were placed on each side of the receptacle. These lights were not used during initial training, but they were introduced later to signal the duration of auditory stimuli (left cue light for DS, right for NS), to facilitate video analysis.

### Video tracking and mobility analysis

Four out of eight operant chambers were equipped with a video tracking system (Cineplex, Plexon Inc.), allowing us to perform video tracking analysis for half of the rats from each cohort. Video recordings for the intra-NAc infusion experiments (cohort 1) were incomplete due to technical problems and, therefore, no tracking analysis for these experiments is presented. However, visual assessment of the available incomplete video recordings allowed us to conclude that the effects of intra-NAc SCH on mobility and sleep were not substantially different from those of systemic SCH. The tracking analysis for the sessions for which complete recordings were obtained was performed using the whole-body tracking mode of the Cineplex software to determine the coordinates of the centroid of the animal’s two-dimensional projection on the horizontal plane. Periods of immobility were determined in R using custom code. To do so, the data for *x* and *y* centroid positions were smoothed (moving average in 1 s time bins) and standard deviations *sd(x)* and *sd(y)* of these coordinates within a 1 s time window centered around each time point were computed. A rat was considered immobile at time *t* if *sd(x) < thr* and *sd(y) < thr*, where the threshold *thr* was set so that the automated detection of immobility periods agreed with our visual assessment. The threshold was set once for all the experimental sessions and it was the same for all animals. We verified that although changing the threshold slightly impacted the assessment of the overall time spent by the animals in immobility, it did not substantially change our results or conclusions. The procedure described above determines if the animal was immobile at each time point. Immobility periods were defined as continuous sequences of immobile time points and their lengths were computed. A weighted distribution of lengths of immobility periods was obtained by determining how much time was spent in immobile periods from a designated class of lengths. Visual quantification of video recordings was performed by the observers blind to drug treatment as described in the Results.

### Surgeries

Stainless steel bilateral guide cannulae (27 ga) purchased from Plastics One were implanted under isoflurane anesthesia so that the tips of guide cannulae were 2 mm above the target brain region. The animals were placed in a stereotactic apparatus, the scalp was retracted, and the skull was leveled so that the dorsoventral coordinate of bregma and lambda are equal. Holes were drilled at target coordinates at AP 1.2, ML 2.0 (in mm relative to bregma) and the cannulae were lowered so that their tips were at DV 6.0. The implant was secured with bone screws and dental cement. Animals were treated with enrofloxacin, ketoprofen, and a topical anesthetic/antibiotic powder containing neomycin, isoflupredone and tetracaine (Neopredef). All animals were allowed to recover with ad libitum access to food for at least 7 days before food restriction and operant training resumed.

### Intracranial microinjections

Internal cannulae (injectors) purchased from Plastics One were connected to mineral oil-filled tubing that was connected to Hamilton syringes placed in an electronic syringe pump. The injectors were backfilled with SCH solution (4 µg/µl) or vehicle (saline). Before the operant session, injectors were inserted into guide cannulae. Injector lengths were set so that they extended 2 mm from the tip of guide cannulae thus targeting DV 8.0 mm relative to bregma. 0.5 µl of drug solution was infused into each hemisphere over 2 min and after an additional 1 min diffusion period the injectors were withdrawn. During the infusion period, the animals were allowed to explore the lap of the experimenter as freely as possible, although rotations were partially restricted to prevent tangling of the tubing. The animals were habituated in advance to this partial restraint to minimize their level of stress. To clear the guide cannulae, the day before the experiment the animals had the injectors inserted into the brain but no infusions were performed. After infusions, the animals were placed in operant chambers either immediately or after a 60 min wait period and the session began.

### Systemic injections

SCH, HAL and CAF were purchased from Sigma. SCH and caffeine were dissolved in sterile 0.9% saline so that the final administered volume was 1 ml/kg. HAL was initially dissolved in glacial acetic acid and then diluted in warm 0.9% saline in such a way that the final administered volume for each experiment was 1 ml/kg of HAL in 0.4% acetic acid. Vehicle injections consisted of 0.9% saline. For experiments with cohort 1 of rats, SCH was administered SC while HAL and CAF were administered IP. For experiments with cohort 2, both SCH and HAL were administered IP.

### Histology

Animals anesthetized with pentobarbital were perfused intracardially with saline and 4% paraformaldehyde. 40 µm brain sections were cut on a vibrating microtome and Nissl staining was performed. The sections were mounted on microscope slides and examined under the microscope to locate cannula tracts and injection sites.

### Statistical analysis

The majority of statistical analyses were performed on response ratios, which are defined as the fraction of cues presented in a given period of time to which the animal made a response. To display the time course of the response ratio, sessions were divided into 10 or 20 min bins, and the mean and SEM of the response ratios for all cues occurring in each bin were graphed (e.g., Fig. 2A1). For statistical analysis, we focused on epochs within the session that were sometimes larger than the bin width, in which case we averaged the response ratio for a given subject across bins (e.g., Fig. 2A2). Mean and individual response ratio values ranged from 0% to 100%, often with many points near 100 or 0. Such data is not normally distributed and cannot be normalized by transformation. Therefore, we used a nonparametric test, the permutation test, for paired statistical comparisons on fractional data. To run this test, we first computed *X_actual_*, the cross-subject mean of the differences between the measured variable (e.g., response ratio) under the two conditions, Condition 1 and Condition 2 (e.g., vehicle and drug). We then produced a distribution of all possible mean values by systematically assigning the two values for each subject to Condition 1 and Condition 2 and recomputing *X*, and repeating this for all possible permutations (combinations of assignments across subjects). In most analyses, the *N* was 13, 12 or 8 subjects, and therefore the distributions consisted of a total of 2^13^, 2^12^ or 2^8^ (8192, 4096 or 256) permuted *X* values. Next, we determined the fraction of the distribution that was less than or equal to *X_actual_*, and the fraction that was greater than or equal to *X_actual_*, which constituted exact probabilities (P values) that *X_actual_* was greater and lower, respectively, than the *X* that could be expected under the null hypothesis of no systematic difference in the measured variable under Condition 1 and Condition 2.

Next, to assess statistical significance at the critical value of 0.05 (two-tailed), we considered whether either of the P values obtained as described above indicated that *X_actual_* fell outside the central 95% of the distribution of permuted *X* values. Specifically, if either P value was less than 0.025, we considered the result potentially significant. Because in most cases we made more than one comparison for a given data set, we adjusted this critical significance level (0.025) for multiple comparisons using the Holm-Sidak method (Glantz, 2002). We report the unadjusted P values in the Results, and in the figures we indicate significant P values that survive multiple comparison adjustment with asterisks (*). P values less than 0.025 that did not survive adjustment for multiple comparisons were considered trends and are indicated in the figures with a hash (#).

For some comparisons (Figs. 7, 8), the *N* was 6, and therefore the distribution of permuted *X* values consisted of only 2^6^ = 64 values. The smallest possible P value (e.g., the cases in which *X_actual_*was either greater or less than all 63 other permuted *X* values) was 1/64 = 0.016, and the next smallest was 2/64 = 0.031. Because adjusting the critical P value of 0.025 for multiple comparisons would yield a value lower than the smallest possible value, it was not possible to adjust for multiple comparisons. Therefore, in these figures we indicate comparisons that yielded P values of 0.016 or 0.031 with single (†) or double (‡) daggers, respectively. Finally, in some further cases, the *N* was 4, such that a distribution of permuted *X* values would contain only 16 values and the smallest possible P value would be 1/16 = 0.063. We did not compute P values in these cases, but we report the results in supplemental figures (Figs. S5, S6B3, B4).

In cases in which absolute counts or summed times were analyzed (i.e., data that was not fractional), the data was generally found to be normally distributed (Kolmogorov-Smirnov test). Therefore, the results were compared using two-tailed paired or unpaired t-tests, with the critical P value (0.05) adjusted for multiple comparisons with the Holm-Sidak method (Figs. 10A,B1,B2,C1,C2, 11B, S6A,B1,B2). Significant P values that survived adjustment for multiple comparisons are indicated in the figures by an asterisk (*), and those that were < 0.05 but did not survive multiple comparison adjustment are indicated by a hash (#). All statistical analyses were conducted using Excel functions (t-tests, Pearson correlations) or custom-written macros (permutation tests).

**Figure 1.**
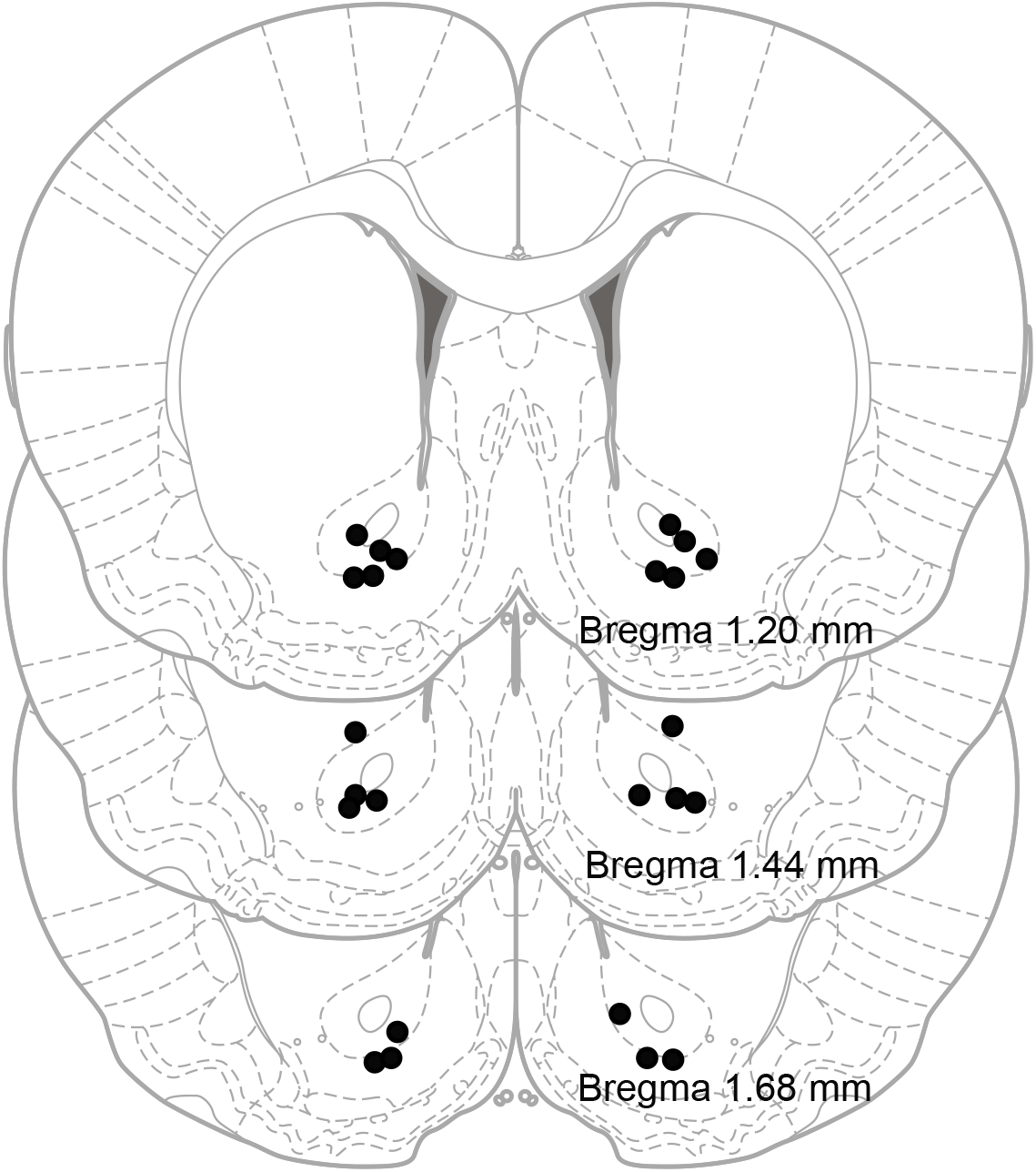
Histological reconstruction of the SCH23390 microinfusion sites in cohort 1. Black dots represent the estimated locations of the tips of injectors, located 2 mm beneath the tips of the guide cannulae. The schematic picture of the coronal brain slice was redrawn from “The Rat Brain in Stereotaxic Coordinates” by G. Paxinos and C. Watson.

**Figure 2.**
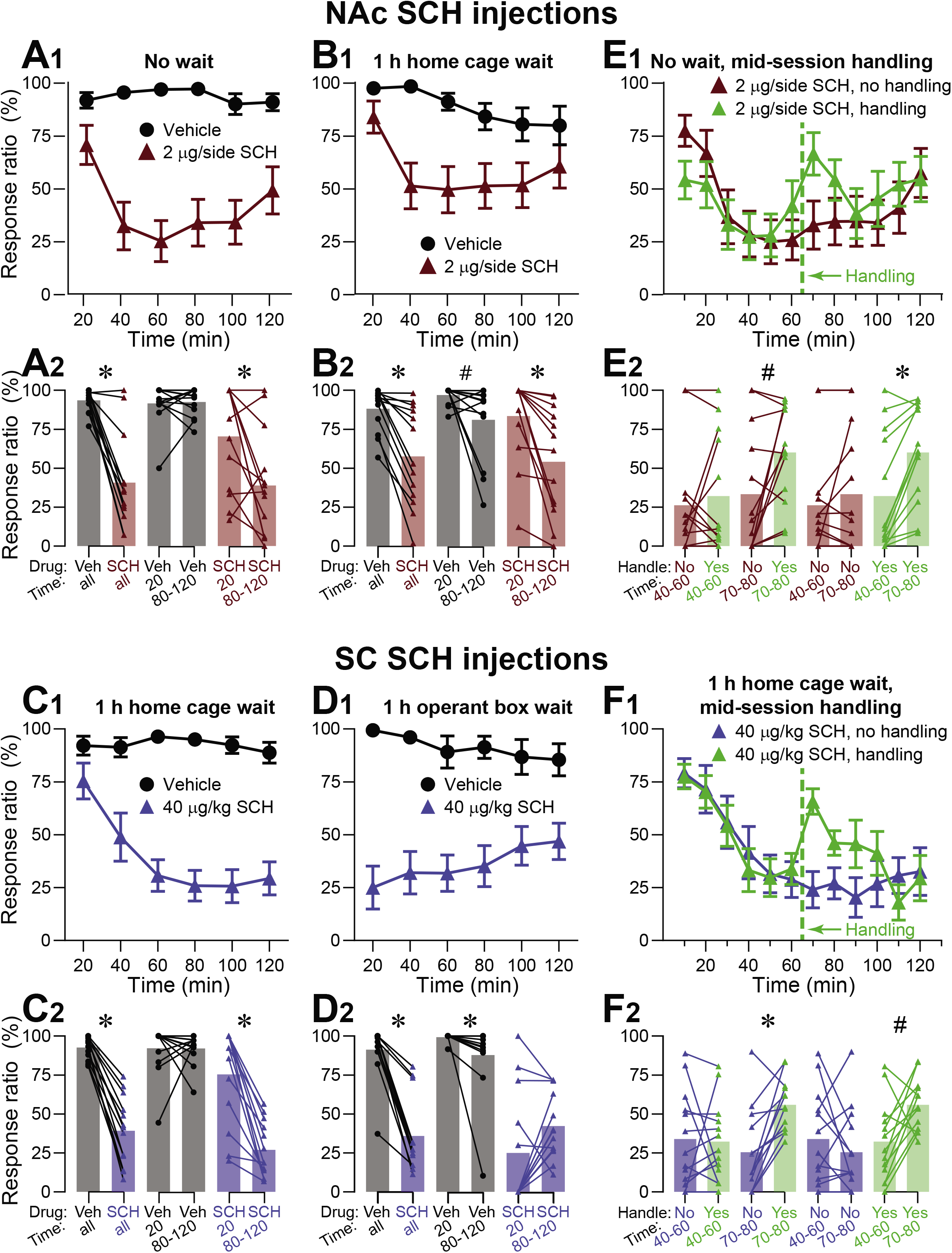
Elevated arousal abolishes the inhibitory effects of intra-NAc and systemic SCH23390 on cued sucrose seeking. Time course graphs show the cross-animal mean ± SEM of the DS response ratio, defined as the percentage of cues presented within sequential 20 min (A-D) or 10 min (E, F) time bins during which the animal made a receptacle entry response. Session length was 2 h. In the bar graphs, the superimposed dots show each subject’s mean DS response ratio across the time bins in the indicated epochs, and the bars show the mean of these values across subjects. SCH or vehicle was administered into the NAc core (2 µg/hemisphere, A, B, E) or subcutaneously (40 µg/kg, C, D, F). Intra-NAc administration of SCH immediately before (A) or 1 h before (B) the operant session resulted in a gradual decline in cue responding. Similarly, systemic administration of SCH 1 h before the session resulted in a gradual decline in responding if animals spent the time between drug administration and session onset in their home cage (C), but not if they waited in the operant box (D). High DS responding was transiently restored by mid-session handling of the animals that received SCH intracranially (E) or systemically (F). This procedure, indicated by vertical dashed green lines in (E, F), consisted of opening the chamber door and 10 – 20 s of gentle handling. Statistical tests were performed only on the data displayed in the bar graphs. Symbols above the connecting lines show the results of paired comparisons (two-tailed permutation test) between the points connected by the lines: *, P < 0.05, adjusted for multiple comparisons; #, P < 0.05, but the result did not reach significance after adjusting the critical P value for multiple comparisons; no symbol, P > 0.05.

The significance of Pearson correlation coefficients in Fig. 9 was established by calculating the t statistic *t* = 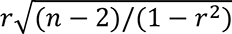, where *r* is the Pearson correlation coefficient and *n* is the number of data points used to compute it. Subsequently, P values were obtained using the Excel function as *p* = *T*. *DIST*. 2*T*(*ABS*(*t*), *n* − 2). For the visual quantification of video recordings by the observers blinded to drug treatment, the probability of the null hypothesis was calculated exactly. In this experiment, four animals were treated with SCH and the other four with HAL. Both independent observers concluded that four out of eight rats had clear sleep episodes and these animals happened to be treated with SCH. The null hypothesis posed that this association happened by chance. The number of 4-element subsets of an 8-element set is equal to:

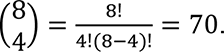

The subset of four rats that received SCH represents one of these 70 subsets and, therefore, the probability that it is selected by chance is 1/70. The probability that two independent observes perform such identification by chance is equal to the product of probabilities, and so p = (1/70)^2^ = 0.0002.

## Results

### Heightened arousal abolishes the effects of intra-NAc and systemic SCH on responding to sucrose-predictive cues

In order to determine whether the delay in the effects of intra-NAc D1 receptor antagonism on responding to sucrose-predictive cues (Yun et al., 2004) is due to slow pharmacodynamics, an extinction-like phenomenon, or a gradual decline in arousal, we trained rats on a cued sucrose seeking task in which an auditory DS predicts delivery of sucrose reward upon entry into a receptacle, whereas responses during a distinct NS have no programmed consequence. These stimuli were presented on a random interval schedule with 50 s mean ITI. Rats were then implanted with bilateral cannulae into the NAc core (Fig. 1). The response ratio (proportion of cues to which the animal responded with a receptacle entry) was analyzed separately for the DS (Fig. 2) and NS (Fig. S1). The results of this and all subsequent analyses of NS response ratios are presented in supplemental figures. Intra-NAc administration of the D1 antagonist SCH (2 µg/hemisphere) immediately before the beginning of the 2 h operant session caused a gradual decline in DS responding with a maximal effect at 1 h (Fig. 2A1). Permutation tests revealed that the mean DS response ratio across the entire session differed after SCH vs vehicle injection (P = 0.0002, N = 13, Fig. 2A2), and that the DS response ratio was greater in the last 1 h of the session than the first 20 min in SCH (P = 0.008, Fig. 2A2) but not vehicle (P = 0.59, Fig. 2A2). The response ratio in the last 1 h was significantly lower in SCH than in vehicle (P = 0.0002, compare the “Veh 80-120” to “SCH 80-120” bars in Fig. 2A2), whereas the response ratio in the first 20 min was not lower in SCH (P = 0.037).

We reasoned that if the delay in the effects of SCH were due to slow pharmacodynamics then no delay should be observed if the drug were infused 1 h before the beginning of the session. Therefore, in the next experiment, animals were placed in their home cages after drug administration and then transferred to the operant box at 1 h, at which point the session was started. (According to the previous experiment, 1 h is the time point at which the drug effect should be maximal.) Although the overall effect of the drug was weaker, a gradual decline in cue responding with maximal effect at 1 h after the beginning of the session was observed again (Fig. 2B1), confirmed by permutation tests showing that the mean response ratio across the session was lower in SCH than in vehicle (P = 0.0001, N=13, Fig. 2B2) and that in SCH, responding was lower in the last 1 h than the first 20 min (P = 0.0001, Dig. 2B2). Although the response ratio in vehicle also trended towards lower values in the last 1 h than the first 20 min (P = 0.021, Fig. 2B2), responding in the last 1 h in SCH was significantly lower than in vehicle (P = 0.0005, compare last 1 h bars for SCH vs vehicle in Fig. 2B2) whereas this was not the case for the first 20 min (P = 0.066, compare 20 min bars in Fig. 2B2).

These results rule out slow pharmacodynamics as a possible explanation of the delayed effects of intra-NAc SCH on cued sucrose seeking. They also demonstrate that high responding to sucrose-predictive cues at the beginning of the session is possible even when SCH, delivered into the NAc at a very high dose (Yun et al., 2004), acts at its full efficacy, suggesting that the activity of NAc D1 receptors is not essential for high initial performance. To determine whether the activity of D1 receptors in other brain regions is necessary for cued sucrose seeking at the beginning of the session, we evaluated the effects of systemic SCH (40 µg/kg SC) on performance in our task. After receiving SCH or vehicle, rats were returned to their home cage and the session commenced 1 h later, the time point when the efficacy of subcutaneously administered SCH is already past its peak and decreasing (Lappalainen et al., 1989). Similar to the intra-NAc experiments, we observed high initial responding that gradually declined (Fig. 2C1). The mean DS response ratio across the entire session was lower in SCH than in vehicle (P = 0.0001, N = 13, Fig. 2C2), and in SCH responding was lower in the last 1 h than in the first 20 min (P = 0.0001, Fig. 2C2) whereas this was not the case in vehicle (P = 0.5, Fig. 2C2). Responding in the last 1 h was lower in SCH than in vehicle (P = 0.0001, compare 80-120 min bars for SCH and Veh in Fig. 2C2), but not in the first 20 min (P = 0.05). Therefore, a very high rate of responding to sucrose-predictive cues at the beginning of the session is possible even if D1 receptors are antagonized systemically. Thus, the results so far suggest that D1 receptors in the NAc or elsewhere are not required for cued approach performance at the beginning of the session.

Having ruled out the slow pharmacodynamics hypothesis, we set out to differentiate between the most likely remaining possibilities: the gradual decline in responding caused by SCH is the result of an extinction-like process, or it is due to declining arousal. Extinction should occur only when the animal responds to cues and earns rewards, which then fail to reinforce further cue responding due to the presence of SCH. On the other hand, if a gradual decline in arousal occurs after the animal is placed in the operant chamber, it should happen regardless of whether cues and sucrose rewards are presented. Therefore, we reasoned that the extinction and arousal hypotheses could be differentiated by allowing the animal to wait in the operant chamber (rather than the home cage) for 1 h after systemic SCH injection. After the 1 h wait, the operant session was quietly started without handling the animal or alerting it in any way other than presenting the first cue. After these manipulations, the high initial performance was abolished in SCH-treated rats, whereas the 1 h wait in the operant box did not impact the DS responding of vehicle-treated animals (Fig. 2D1). The mean DS response ratio was lower in SCH than in vehicle (P = 0.0001, N = 13, Fig. 2D2), and while the response ratio in vehicle was slightly slower in the last hour than in the first 20 min (P = 0.004, Fig. 2D2), this was not the case in SCH (P = 0.97, Fig. 2D2). These results indicate that, in contrast to the results presented so far, SCH did not cause a gradual decline, a conclusion that is further confirmed by the observation that the response ratios in both the first 20 min (P = 0.0002) and last 1 h (P = 0.0004) were lower in SCH than in vehicle (compare 20 min SCH vs vehicle bars, and last 1 h SCH vs vehicle bars, in Fig. 2D2). These results are incompatible with the extinction interpretation because within this interpretation, the rats would have to have consumed some rewards under the influence of SCH in order for their performance to decline. This was clearly not the case, since most of the rats (8 out of 13) were so unresponsive to cues after the 1 h in-chamber wait that they did not obtain any rewards at all during the first 20 min time bin (Fig. 2D2). The most likely hypothesis is therefore that transferring the animal to the operant chamber elevates its arousal level, resulting in cue responding that is resistant to SCH until the arousal level declines.

Multiple factors could contribute to a heightened arousal level at the beginning of the session, including the introduction of the animal into the operant box context and handling animals during the transfer. To assess the importance of handling, we re-handled SCH-injected rats in the middle of the session by opening the operant box door and touching them for 5-10 seconds (Supplementary Video 1). When SCH was injected in the NAc, mid-session handling caused a brief increase in DS responding la5/12/2022sting approximately 20 min (Fig. 2E1). The response ratio was significantly increased in the 20 min after handling at 60 min (mean of 70 and 80 min bins; note that the times indicate the end of the 10 min bin) compared with the 30 min immediately prior (mean of the 40, 50 and 60 min bins) (P = 0.0006, N = 13, Fig. 2E2), whereas comparison of the same time bins in animals that were not handled yielded no significant effect (P = 0.20, Fig. 2E2). There was a trend towards a greater DS response ratio in the 20 min occurring after handling (70-80) than in the same epoch in animals that were not handled (P = 0.01, Fig. 2E2) whereas, as expected, there was no difference in response ratio in the 40-60 min (pre-handling) epoch in sessions in which animals were later handled vs sessions in which they were not (P = 0.25, Fig. 2E2). Animals injected systemically with SCH showed similar results (Fig. 2F1), with a trend towards a significant increase in DS responding in the 20 min post-handling vs the 30 min prior (P = 0.011, N= 13, Fig. 2F2) but not when the same epochs were compared in animals that were not handled (P = 0.8, Fig. 2F2). In addition, DS responding was significantly greater in the 20 min post-handling vs the same epoch when the animals were not handled (P = 0.001, Fig. 2F2), whereas this was not the case for the pre-handling 30 min epoch (P = 0.054, Fig. 2F2).

While the experiment in which animals waited for 1 h in the operant chamber after systemic administration of SCH (Fig. 2D) clearly indicates that impaired performance cannot be explained by an extinction-like phenomenon, a similar experiment could not be performed after intra-NAc administration because the drug would likely diffuse to other brain regions within the 1 h waiting period and the effects observed after this time could not be unequivocally attributed to the local effects of SCH. Although the ability to reactivate high rates of sucrose seeking with mid-session handling in rats administered SCH into the NAc (Fig. 2E) supports the involvement of decreased arousal in the gradual decline of responding, we sought an independent behavioral variable that would allow us to differentiate the extinction and arousal hypotheses. We reasoned that if SCH causes sucrose to be less reinforcing, animals would tend to leave the receptacle earlier, resulting in shorter entries. On the other hand, decreased arousal should slow animals’ performance and thus promote longer entries.

The time the animals spent in the receptacle strongly depended on whether sucrose was delivered as a result of an entry (rewarded entry) or not (unrewarded entry). The distribution of the lengths of entries in vehicle-treated animals (Fig. 3A) was bimodal, with the two peaks at 1-2 s and 4-5 s corresponding to unrewarded and rewarded entries, respectively. After intra-NAc administration of SCH, animals spent overall less time in the receptacle (Fig. 3B, note that the axes are scaled the same for Fig. 3A and 3B), which was expected, since the number of responses to both DS and NS was decreased (Figs. 2A, S1A). However, we also noticed a change in the distribution of entry lengths; the first peak at 1-2 s almost completely disappeared and another peak emerged for long entries (> 7 s). To precisely examine the impact of SCH on the shape of the distribution, we normalized the data so that the total time spent in a receptacle by each rat is 100%. Intra-NAc SCH injection decreased the percentage of time allocated to short entries (1-2 s) and increased the time allocated to long entries (> 7 s) compared to vehicle injection; however, although the P values for these comparisons were lower than the unadjusted critical value of 0.025 (1-2 s: P = 0.003; >7 s: P = 0.013; N = 13, Fig. 3C), these values exceeded the critical values adjusted for 8 comparisons. Importantly, SCH-treated animals typically took 3-6 s to consume the rewards, just as they did after vehicle treatment, suggesting that their appreciation of the reward was not diminished. However, after some entries (either rewarded or not, Fig. 3B), animals tended to stay in the receptacle much longer than necessary, resulting in the new peak of the distribution for entries longer than 7 s (Fig. 3C). Freezing in the receptacle for elongated periods of time is consistent with performance problems such as drowsiness or catalepsy and certainly cannot result from impaired reinforcement. We also compared the distributions of receptacle entry lengths after systemic administration of either vehicle or SCH and observed an even more pronounced shift towards longer times (0-1 s: P = 0.0002; 1-2 s: P = 0.003; 3-4 s: P = 0.003; >7 s: P = 0.0007; N=13, Fig. 3D), suggesting that antagonism of D1 receptors outside the NAc aggravates the performance problems.

**Figure 3.**
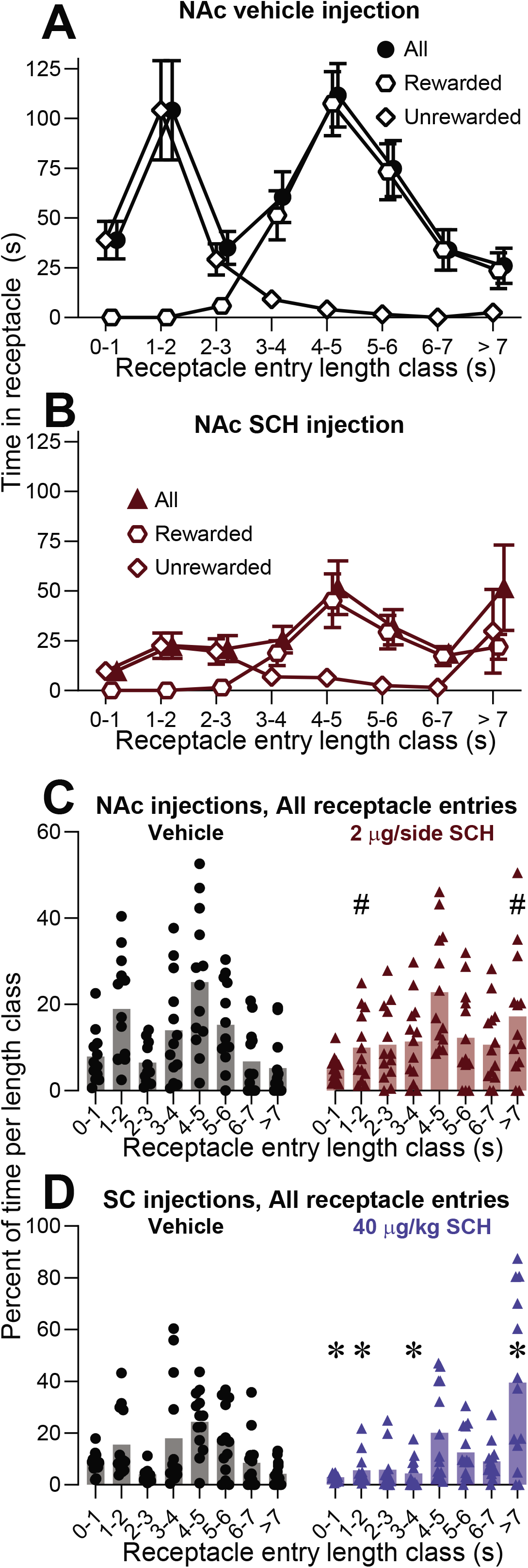
Intra-NAc and systemic SCH23390 injections promote longer receptacle entries. Analysis of the lengths of receptacle entries in rats administered vehicle or SCH into the NAc (A, B, C) or systemically (D). The data used for the analysis correspond to the experiments shown in Figs. 2A and C. A, B, Time spent in the receptacle during entries belonging to a class of length indicated on the abscissa after intra-NAc administration of vehicle (A) or SCH (B). The total time per class (filled symbols) is the sum of the time of rewarded entries (opem hexagons) and unrewarded entries (open diamonds). C, D, Percent of the total time spent in the receptacle that can be attributed to entries from a class of length indicated on the abscissa after intra-NAc (C) or subcutaneous (D) administration of either vehicle (black symbols, grey bars) or SCH (colored symbols and bars). The data were normalized so that the total time spent in the receptacle by each rat is equal to 100%. Symbols above the SCH bars show the results of paired comparisons (two-tailed permutation test) to the corresponding bar in vehicle: *, P < 0.05, adjusted for multiple comparisons; #, P < 0.05, but the result did not reach significance after adjusting the critical P value for multiple comparisons; no symbol, P > 0.05.

Together, these data show that the gradual decline in responding to reward-predictive cues after administration of the D1 blocker is caused by a decline in arousal, rather than slow pharmacodynamics or an extinction-like process. The similarity between these arousal effects in animals administered SCH systemically vs. intracranially into the NAc suggests that the effects observed after systemic administration are due at least in part to blockade of D1 receptors within the NAc.

### HAL-induced disruption of cued approach behavior is not alleviated by handling-induced arousal

Having established that arousal potently modulates the effects of the D1 antagonist on cued approach for sucrose, we sought to determine whether similar effects are observed when D2-like dopamine receptors are antagonized. It has been previously shown that intra-NAc administration of a selective D2 receptor antagonist raclopride caused an immediate inhibition of cued approach, rather than a gradual decline (Yun et al., 2004). However, the short duration of the effects of raclopride would not allow for mid-session handling experiments. We therefore investigated the impact of the D2/D3 receptor blocker HAL, characterized by its slow pharmacokinetics (Lappalainen et al., 1989), on performance in our task. We chose doses of HAL within the 100 – 200 µg/kg range, in which off-target activity on nondopaminergic receptors has been shown to be negligible (Anden et al., 1970). Rats were systemically administered HAL (150 µg/kg IP) or vehicle and allowed to wait for 1 h in their home cages before the beginning of the operant session. In stark contrast with the effects of SCH, we observed an immediate inhibitory effect on responding to sucrose-predictive cues (Fig. 4A; see Fig S2 for NS results). The mean DS response ratio across the session was significantly lower after HAL injection than after vehicle injection (P = 0.0002, N = 12, Fig. 4A), and there was no within-session decline in responding as indicated by the absence of significant differences in the response ratio in the last 1 h vs the first 20 min in either vehicle (P = 0.33, Fig. 4A) or HAL (P = 0.94, Fig. 4A). The response ratio in HAL was lower than in vehicle in both the first 20 min (P = 0.0002, compare Veh 20 min and HAL 20 min bars in Fig. 4A) and the last 1 h (P = 0.0005, compare Veh and HAL 80-120 min bars in Fig. 4A). These results are consistent with earlier studies (see the discussion of Experiment 2 in Fibiger et al., 1976) and show that, unlike for SCH, elevated arousal at the beginning of the session does not abolish the effect of HAL on cued approach.

**Figure 4.**
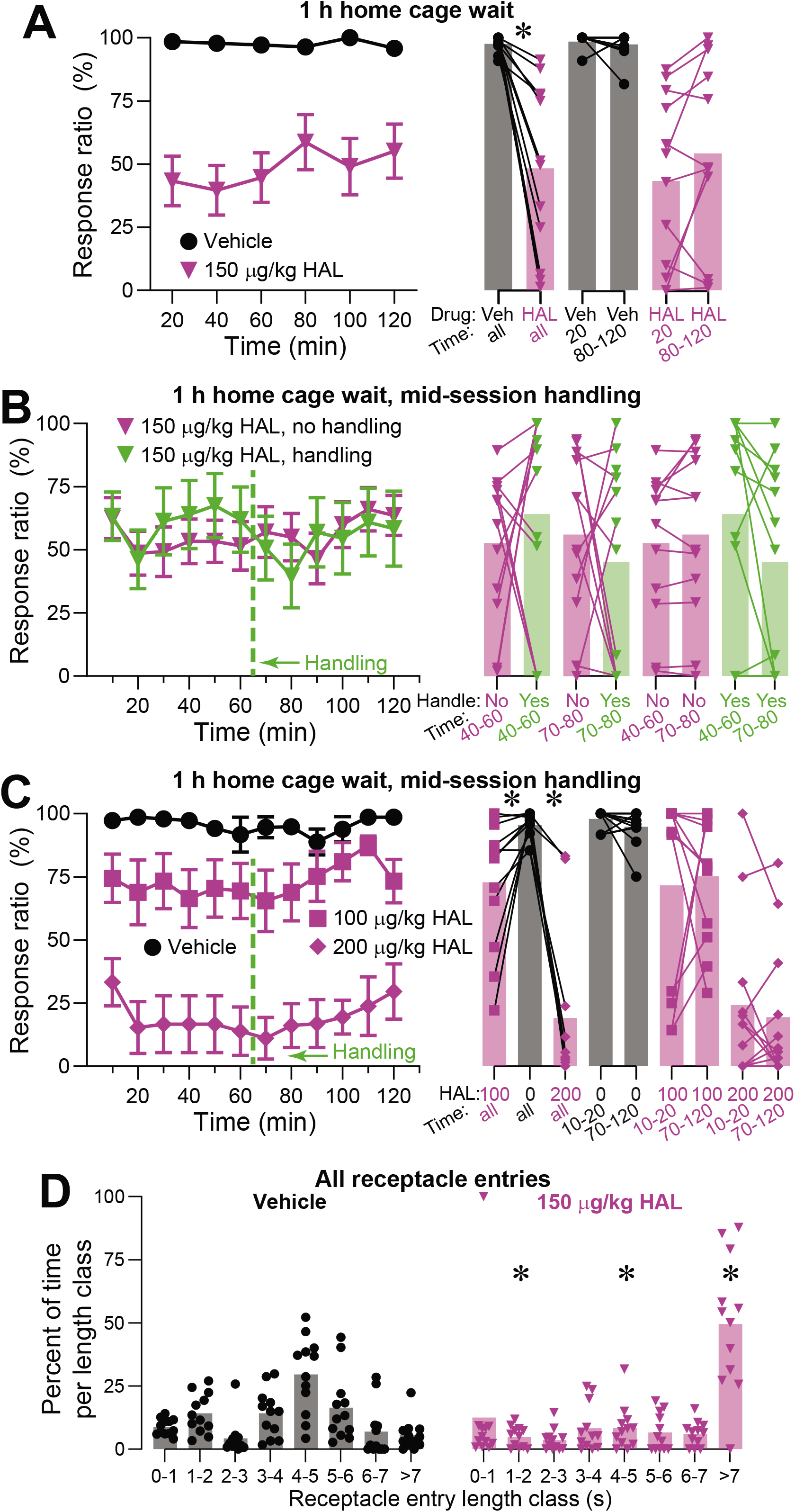
Elevated arousal does not alleviate the effects of haloperidol on cued sucrose seeking. A, B, C, The left graph in each panel shows the mean ± SEM DS response ratio in sequential 20 min (A) or 10 min (B, C) time bins. Rats received intraperitoneal injections of vehicle or the dose of HAL indicated in the legend. After the injection, animals waited 1 h in their home cage before the session began. Animals in the experiments presented in B and C received mid-session handling, indicated by a vertical dashed line. The right graphs show each subject’s mean DS response ratio across the time bins in the indicated epochs (symbols) and the mean of these values across animals (bars). Symbols above the connecting lines show the results of paired comparisons (two-tailed permutation test) between the points connected by the lines: *, P < 0.05, adjusted for multiple comparisons; no symbol, P > 0.05. D, Analysis of the normalized distribution of receptacle entry lengths for the experiment shown in A, following the same format as Fig. 3C,D. Symbols above the HAL bars show the results of paired comparisons (two-tailed permutation test) to the corresponding bar in vehicle: *, P < 0.05, adjusted for multiple comparisons; no symbol, P > 0.05.

We then directly investigated the impact of handling-induced arousal on the performance of HAL-treated rats with the mid-session handling procedure. In two separate experiments, animals that had been administered the same dose of HAL (150 µg/kg) were either handled in the middle of the session or not and their performance was compared (Fig. 4B). When examined with 10 min temporal resolution, the DS response ratio fluctuated, but did not follow any noticeable trend across time. The response ratio did not differ in HAL vs vehicle in either the 20 min after mid-session handling at 60 min or in the preceding 30 min; nor was there an increase in responding in the 20 min after handling vs the preceding 30 min in either vehicle or HAL (P > 0.14 for all comparisons, Fig. 4B, N = 12).

The ability of handling to enhance responding could depend on the administered dose of the dopamine antagonist. Although the doses of SCH and HAL used in our study (40 and 150 µg/kg, respectively) produced similar degrees of overall inhibition of cued approach, we investigated the possibility that other doses of HAL could produce an effect that does depend on arousal. We administered 0, 100, or 200 µg/kg of HAL, ran the animals on the DS task after a 1 hr wait in the home cage, and handled the animals mid-session (Fig. 4C). HAL dose-dependently inhibited cued approach (for entire session mean DS response ratio, P = 0.001 for 100 µg/kg HAL and P = 0.0002 for 200 µg/kg HAL, N = 12, Fig. 4C) but within-session declines were not observed (DS response ratio in the last 1 h vs first 20 min: vehicle, P = 0.13; 100 µg/kg HAL, P = 0.61; 200 µg/kg HAL, P = 0.20, Fig. 4C). There were no clear effects of mid-session handling (Fig. 4C). Finally, the effects of 150 µg/kg HAL on the time spent in the receptacle were similar but even more pronounced than those of SCH (compare Fig. 3D with 4D); 50% of the total time spent by HAL-treated animals in the receptacle was accounted for by entries longer than 7 s, which almost never occur after vehicle administration (comparisons of HAL to vehicle: 1-2 s, P = 0.002; 4-5 s, P = 0.004; >7 s, P = 0.0005; Fig. 4D).

We conclude that elevation of arousal, caused by either transfer of animals into the operant chamber or mid-session handling, has no impact on the probability of response to the reward-predictive cues in rats administered HAL, which distinguishes the effects of HAL from those of SCH. On the other hand, both dopamine antagonists induce freezing in the receptacle for extended periods of time, indicating that reward-independent performance problems contribute to impaired responding to the reward-predictive cues.

### Caffeine abolishes the effects of either SCH or HAL on responding to sucrose-predictive cues

Our finding that the inhibition of cued approach behavior caused by HAL cannot be reversed by manipulations that increase arousal was surprising because HAL-induced deficits in performance of a fixed-ratio 5 task have been shown to be reversed by an arousing drug, the adenosine receptor antagonist CAF (Salamone et al., 2009). To determine whether CAF similarly opposes the effects of dopamine receptor blockade on performance of our cued approach task, we administered SCH or HAL, allowed animals to wait for 1 h in the home cage, and then injected them with CAF (10 mg/kg IP) or vehicle immediately before the beginning of the operant session. We first asked whether CAF is as effective as natural arousal in alleviating the effects of SCH. The gradual decline in DS responding observed after administration of SCH was alleviated by CAF (whole-session mean DS response ratio in SCH+CAF vs in SCH+vehicle: P = 0.013, N= 13, Fig. 5A). Next, we examined the effect of CAF on performance of HAL-treated rats. In contrast to natural arousal, CAF almost completely prevented the HAL-induced inhibition of cued approach (whole-session mean DS response ratio in HAL+CAF vs in HAL+vehicle: P = 0.0002, N= 12, Fig. 5B). We conclude that in SCH-treated rats, CAF and arousal elicited by natural stimuli similarly affect cued approach behavior. However, whereas CAF counteracts the effects of HAL on performance, natural arousal does not.

**Figure 5.**
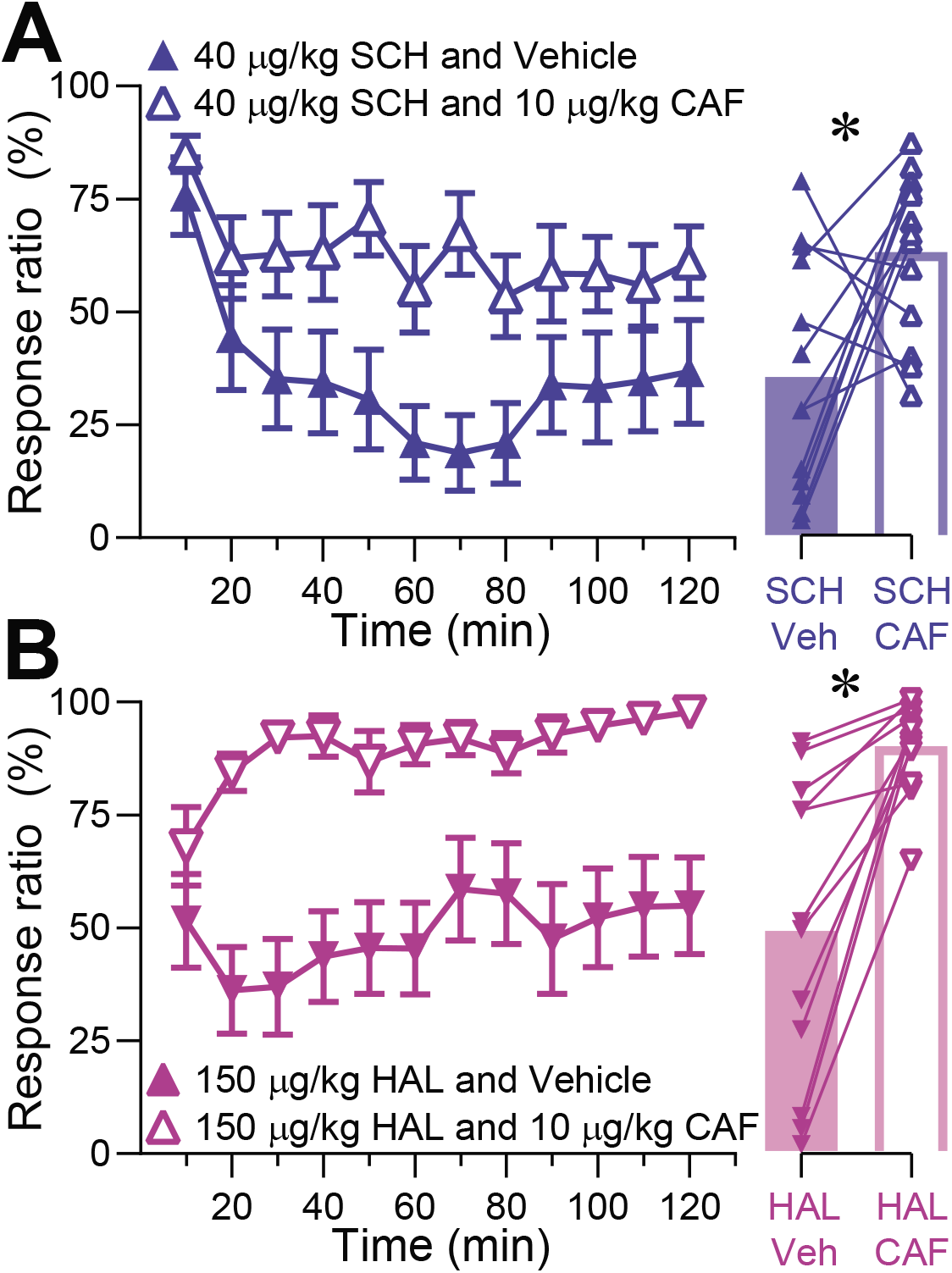
Effects of caffeine on sucrose seeking in rats administered haloperidol and SCH23390. The left graphs show the mean ± SEM DS response ratio in sequential 10 min time bins for rats that received the following drug administrations: A, 40 µg/kg SCH 1 h before the session, and either vehicle or 10 mg/kg CAF immediately before the session; B, 150 µg/kg HAL 1 h before the session, and either vehicle or 10 mg/kg CAF immediately before the session. The symbols in the right graphs show each subject’s mean DS response ratio across the entire session, and the bars indicate the mean of these values across subjects. *, P < 0.05, two-tailed permutation test; no symbol, P > 0.05.

### Confirmation of the differential impact of natural arousal on rats administered SCH and HAL

All of the preceding results were collected in longitudinal experiments performed on the same cohort of animals, and most of the experiments involving SCH were performed prior to those involving HAL. Consequently, even though experiments were always separated by multiple sessions during which no drugs were administered and normal baseline levels of reward seeking were observed, it is possible that experimental history or order effects influenced the results. Moreover, SCH was administered SC whereas HAL was administered IP, which, although unlikely, could also have contributed to the differences between SCH and HAL effects. To rule out these possibilities, we trained a new cohort of 16 drug-naïve animals on the same cued sucrose seeking task and then randomly assigned them to two groups. These groups exhibited similar baseline performance (compare vehicle conditions for groups 1 and 2 in Fig. 6A,B). In a parallel experiment, group 1 was administered 40 µg/kg SCH and group 2 was given 150 µg/kg HAL; all injections were IP. After drug administration, animals waited for 1 h in their home cages until the operant session commenced. Intriguingly, the overall magnitudes of the effects were different in the new cohort compared with the old. Specifically, these animals’ sensitivity to SCH was lower (session mean of the DS response ratio in 40 µg/kg SCH trended towards being lower than vehicle, P = 0.016, N = 8, Fig. 6A; compare the magnitude of the reduction to Fig. 2C) whereas their sensitivity to HAL was higher (session mean in 150 µg/kg HAL was significantly different from vehicle, P = 0.004, N = 8, Fig. 6B; compare the magnitude of the reduction to Fig. 4A). Despite these differences, consistent with our hypothesis, the impact of 150 µg/kg HAL on cued sucrose seeking was immediate (difference between the last 1 h and first 20 min: P = 0.75 in HAL and P = 0.84 in vehicle, Fig. 6B) while the effect of 40 µg/kg SCH developed gradually, although the difference between the last 1 h and first 20 min did not reach significance (P = 0.06 in SCH, P = 0.87 in vehicle, Fig. 6A). Because of the differences in sensitivity of cohort 2 vs cohort 1, we repeated the experiment, but the assignment of animals to the HAL and SCH groups was reversed and the dosage was adjusted. These experiments were performed after a six-day period of recovery, during which the animals’ return to baseline performance was confirmed in a control operant session. 60 ug/kg SCH and 100 ug/kg HAL produced effects of similar overall strength (Fig. 6A,B): the total DS response ratios throughout the entire session were 26 ± 1 % and 28 ± 2 % (mean ± SEM) for SCH and HAL, respectively, which differed significantly from vehicle (60 ug/kg SCH: P = 0.004, N = 8, Fig. 6B; 100 ug/kg HAL: P = 0.004, N = 8, Fig. 6A). However, we observed that the decline in performance after SCH was gradual and the effect of HAL was immediate, again confirming our hypothesis (trend towards significant reduction in the last 1 h vs the first 20 min in 60 ug/kg SCH: P = 0.011, Fig. 6B; no significant reduction in 100 ug/kg HAL: P = 0.56). These observations affirm that drug history did not contribute substantially to our main conclusion that arousal can abolish the effects of SCH on cued approach whereas it cannot alleviate the effects of HAL.

**Figure 6.**
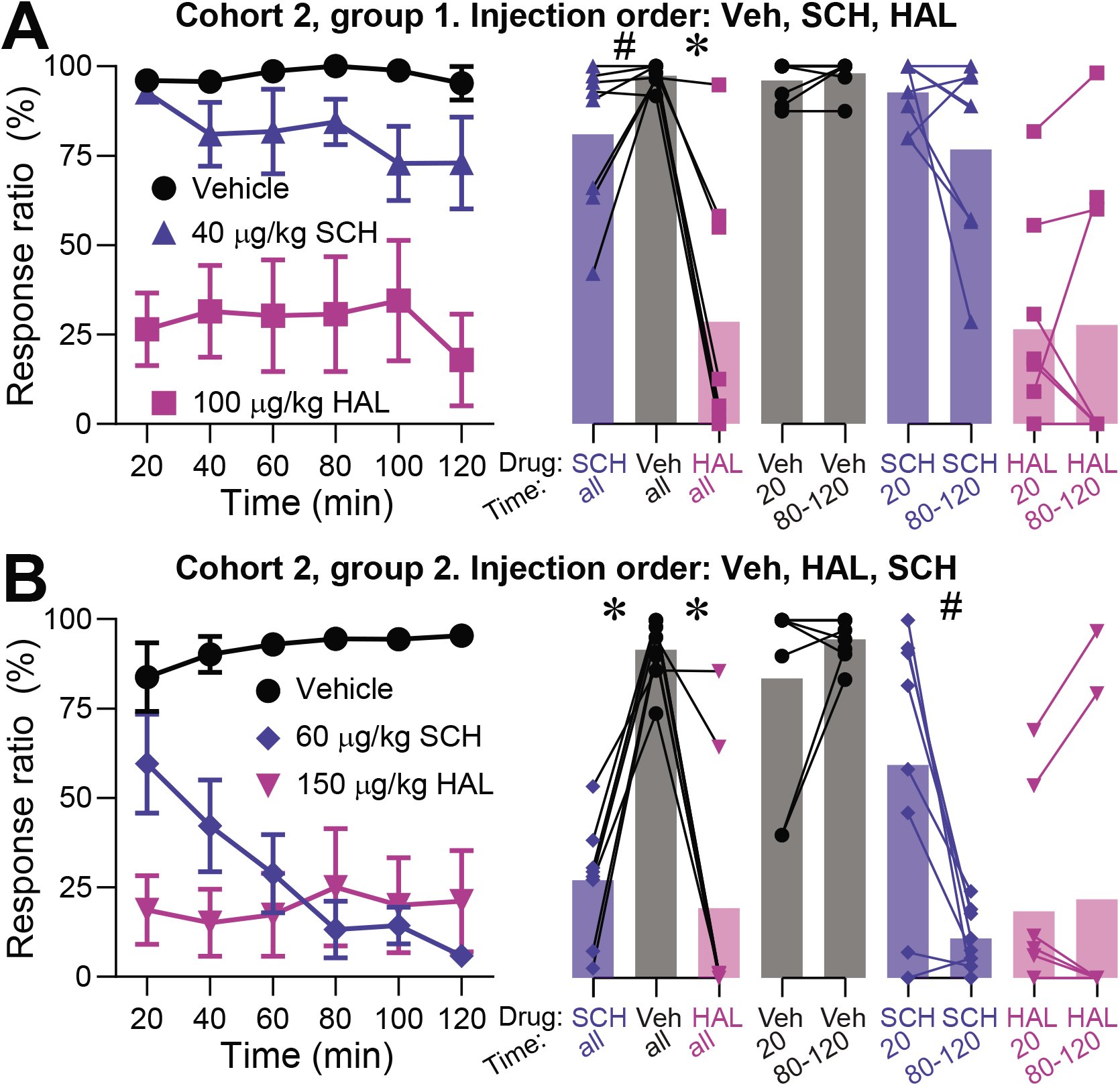
Experiments with cohort 2 confirm differential effects of arousal on reward seeking in animals administered SCH23390 and haloperidol. In cohort 2, 16 rats were divided into group 1 and group 2, which were administered HAL and SCH in different orders and at different doses as indicated in the supertitles and legends. Left and right graphs follow the same conventions as in previous figures. All graphs show the DS response ratio. A, Group 1 received (in order) vehicle, 40 µg/kg SCH, and 100 µg/kg HAL. B, Group 2 received vehicle, 150 µg/kg HAL, and 60 µg/kg SCH. *, P < 0.05, adjusted for multiple comparisons; #, P < 0.05, but the result did not reach significance after adjusting the critical P value for multiple comparisons; no symbol, P > 0.05; two-tailed permutation tests.

### Mobility gradually declines during operant sessions, and it is decreased by both SCH and HAL, but it correlates with DS responding exclusively in SCH-treated rats

In order to gain more insight into the variability of arousal occurring throughout operant sessions, we sought a behavioral variable that is highly correlated with the level of arousal but independent of responsivity to the reward-predictive cues. A behavioral definition of generalized arousal has recently been proposed based on responsiveness to sensory stimuli of all modalities, emotional lability, and the degree of motor activity (Pfaff et al., 2008); these behavioral variables were correlated across a broad range of experimental conditions (Garey et al., 2003; Weil et al., 2010). Multiple research groups have reported a high degree of correlation between animals’ mobility and other measures of arousal, and have described potential neural mechanisms for these relationships (French et al., 1952; Garcia-Rill et al., 2004; Garcia-Rill et al., 2016; Goetz et al., 2016; Grillner et al., 2008; Lee et al., 2014; Lindsley et al., 1950; Shikano et al., 2018; Skinner et al., 2004; Vinck et al., 2015). Motivated by the tight link between arousal and animals’ propensity to move, we analyzed whole-body video tracking data gathered for 6 of 12 subjects from cohort 1, as well as 8 of 16 subjects from cohort 2. We then determined periods of mobility (movement of the rat’s centroid) and immobility (no change in centroid’s position) for each animal (see Methods). Finally, in analogy to the analysis of DS response ratios, we divided operant sessions into 10 min time bins and calculated average mobility within bins as the percentage of time spent in motion.

Rats administered vehicle spent more than 70% of their time in motion at the beginning of the session, but mobility gradually declined to less than 50% after 30 min and remained restricted within the 30 – 50% range until the end of the session (Fig. 7A1, black circles; mobility in the last 1 h vs the first 10 min, P = 0.016, N = 6, Fig. 7A2). Despite this decline in mobility, animals responded to almost all sucrose-predictive cues throughout the vehicle injection session (Fig. 7B1, black circles; there was, however a slight trend towards a lower response ratio in the last 1 h vs the first 10 min, P = 0.031, Fig. 7B2). The initial decline in mobility likely reflects the decline in arousal that occurs independently of the presentation of reward-predictive cues and the associated operant performance. Consistent with this idea, when we transferred animals to the operant chamber and gave them a 1 h waiting period, mobility in vehicle-treated rats similarly declined from 80% to 40% within 30 min from the transfer (Fig. 7C1, black circles; mobility in the last 30 min of the waiting period vs the 1^st^ 10 min, P = 0.016, N = 6, Fig. 7C2), even though no cues were presented to the animals during this time. Mobility remained restricted within the 30 – 50% range in the following 2 h period when cues were presented (Fig. 7C1, black circles), during which vehicle-treated animals responded to nearly 100% of the reward-predictive cues (Fig. 7D1; response ratio in the last 1 h vs the first 10 min after cue presentation start, P = 0.73, Fig. 7D2). Thus, in the control condition, the initial decline in mobility, likely reflecting a decline in arousal, does not prevent the animals from responding to nearly 100% of DS cues.

**Figure 7.**
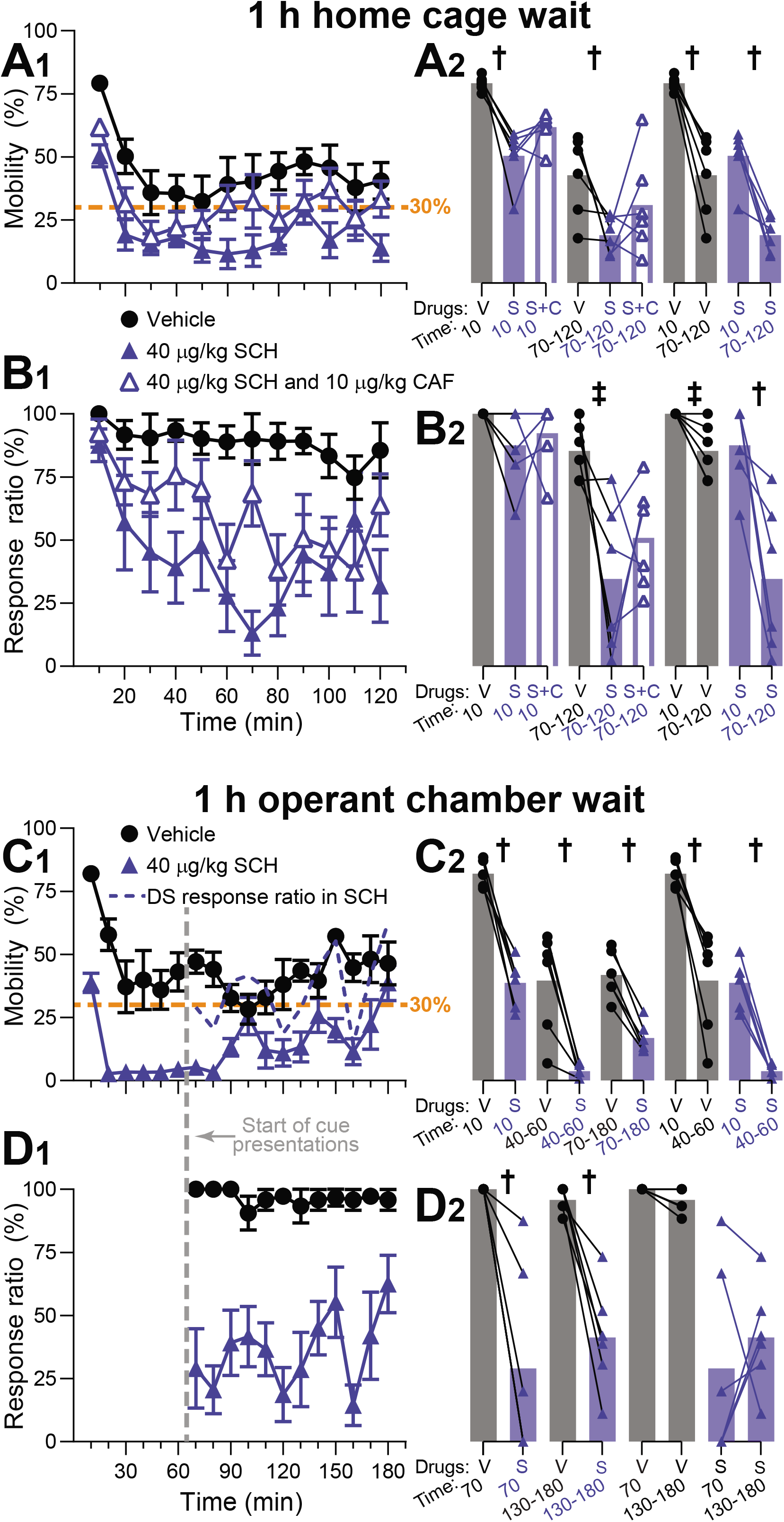
Impact of SCH23390 on the mobility of rats performing cued sucrose seeking task. The graphs show data from the subset of subjects for which video tracking data was available: A, B show 6 rats from the experiment shown in Fig. 5A, and C, D show 6 rats from the experiment shown in Fig. 2D. A1, C1 show mobility (percent of time spent in motion within the indicated time bin) whereas B1, D1 show DS response ratio in the same bins. A2, B2, C2 and D2 show individual subjects (symbols) and mean (bars) values as in previous figures. The dashed data line in C1 shows the DS response ratio after SCH injection re-plotted from D1. Dashed vertical line in C1, D1 indicates the beginning of cue presentation after the 1 h wait in the operant box. Dashed horizontal line in A1 and C1 indicate 30% mobility. Because the N was 6, the lowest possible P value for permutation tests was P = 0.016 and the next-lowest was P = 0.031 (see Methods, Statistical Analysis). †, P = 0.016 for the two-tailed permutation test comparing the two groups connected by lines; ‡, P = 0.031; no symbol, P > 0.045.

In agreement with previous research (Meyer et al., 1993), mobility was lower overall in SCH than in vehicle (Fig. 7A1, closed blue triangles; mobility in the first 10 min in SCH vs vehicle, P = 0.016). During the initial 10 min, SCH-treated animals spent 50% of their time in motion, substantially less than after vehicle injection (Fig. 7A1, first 10 min bin; P = 0.016, Fig. 7A2); yet, despite their reduced mobility in the first 10 min, they responded to nearly 90% of the sucrose-predictive cues, similar to vehicle-treated animals (Fig. 7B1; P = 0.125, Fig. 7B2). Subsequently, mobility and DS responding declined in tandem and remained reduced, compared with vehicle, throughout the remainder of the session (mobility in the last 1 h in SCH vs vehicle, P = 0.016, Fig. 7A2; DS response ratio in the last 1 h in SCH vs vehicle, P = 0.031, Fig. 7B2; DS response ratio in SCH during the last 1 h vs the first 10 min, P = 0.016, Fig. 7B2). Co-administration of CAF caused slight increases in both mobility (Fig. 7A1) and DS response ratio (Fig. 7B1) compared to SCH alone, although these did not approach significance (mobility: P = 0.063 in the first 10 min and P = 0.14 in the last 1 h, Fig. 7A2; DS response ratio: P = 0.31 in the first 10 min and P = 0.16 in the last 1 h, Fig. 7B2). When SCH was administered prior to placing the animals in the operant chamber for a 1 h waiting period, animals showed pronounced inhibition of movement in the absence of reward-predictive cues; almost all movement ceased within 10 min from the transfer (Fig. 7C1; P values for comparisons of mobility in SCH vs vehicle in the first 10 min, in SCH vs vehicle in the last 30 min of the waiting period, and of the last 30 min vs the first 10 min in SCH were all P = 0.016; Fig. 7C2). When cues were presented, the low levels of mobility (10-30%) and DS responding (20-40%) were comparable to the levels observed in the second hour after SCH administration in other experiments (compare Fig. 7D1 to Fig. 7A1; P values for comparisons of DS response ratio in SCH vs vehicle in both the first 10 min after session start and the last 1 h were both P = 0.016).

In summary, our data show that SCH uniformly lowers mobility levels across the session, while DS responding remains unaffected at the beginning of the session when arousal level is high. The data further suggest a simple threshold hypothesis: animals respond to almost all reward-predictive cues whenever their mobility, likely reflecting the level of arousal, is maintained above a threshold of approximately 30% (horizontal dashed line in Fig. 7A1, C1). In vehicle-treated rats, mobility initially declines, but it is maintained above this threshold throughout the session, allowing for continued efficient performance. Although SCH uniformly decreases mobility across the session, elevated arousal at the beginning of the session, caused by transfer into the operant box, prevents mobility from dropping below the threshold and, therefore, DS responding is initially unaffected. As the session progresses through time, the effects of the drug sum with the natural decline of arousal, bringing mobility below the threshold and disrupting cued approach.

Similarly to SCH, administration of HAL uniformly decreased mobility in all time bins (Fig. 8A1,A2; P values for all comparisons indicated by a single dagger are P = 0.016 and P = 0.031 for double dagger; N = 6). Despite this nearly complete inhibition of mobility, HAL-treated rats were nevertheless able to maintain moderate levels of cue responding (about 40%, Fig. 8B1,B2; comparison of HAL to vehicle in the first 10 min, P = 0.063; HAL to vehicle in the last 1 h, P = 0.031; P = 0.5 and P = 0.6875 for comparisons of the first 10 min to the last 1 h in vehicle and HAL, respectively). Co-administration of CAF completely abolished the effect of HAL on both mobility (Fig. 8A) and cued approach (Fig. 8B; comparison of HAL+CAF to HAL+vehicle in the first 10 min and last 1 h, P = 0.125 and P = 0.016, respectively).

**Figure 8.**
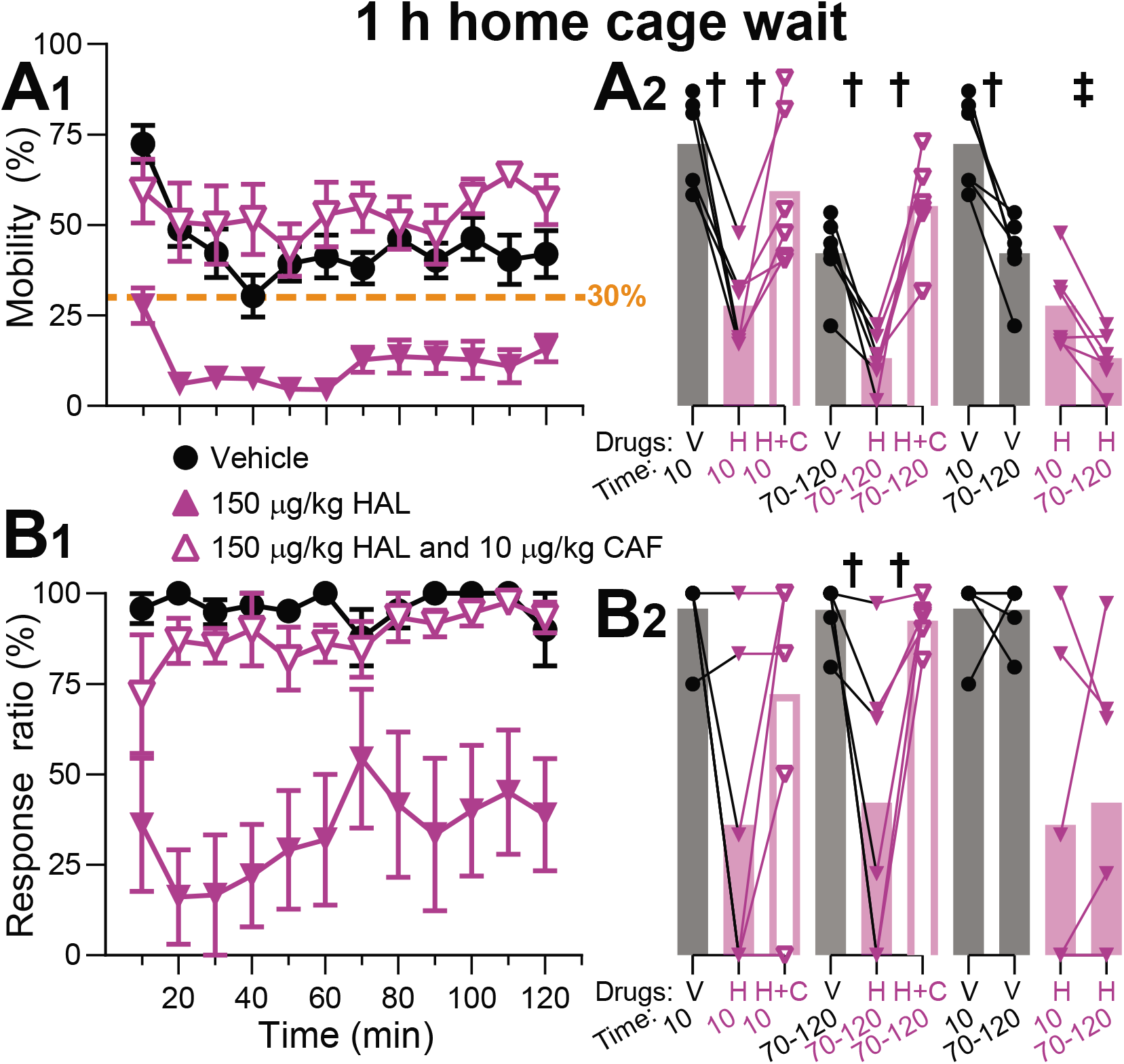
Impact of haloperidol on the mobility of rats performing cued sucrose seeking task. The graphs show data for the subset of subjects used for the experiment shown in Fig. 5B for which video tracking data was available. A1 shows mobility (percent of time spent in motion within the indicated time bin) whereas B1 shows DS response ratio in the same bins. A2 and B2 show individual subjects (symbols) and mean (bars) values as in previous figures. Dashed horizontal line in A1 and C1 indicate 30% mobility. Significance symbols are as in Fig. 7.

Dopamine antagonists have been hypothesized to inhibit movement by inducing catalepsy (Lappalainen et al., 1989; Morelli and Di Chiara, 1985; Van Hartesveldt and Meyer, 1993), disrupting an “activational” component of motivation (Salamone and Correa, 2012; Salamone et al., 2016), or even impairing motor learning (Beeler et al., 2012). However, since both D1 and D2 receptors have also been implicated in the maintenance of wakefulness (Isaac and Berridge, 2003; Ongini et al., 1985; Ongini et al., 1987; Qu et al., 2010), inhibited mobility after antagonism of these receptors can also reflect drowsiness. Importantly, doses of SCH comparable to those used in our study were found to enhance both REM and non-REM sleep in rats within a three-hour window after drug administration; SCH-treated animals slept over three times more than controls (Trampus and Ongini, 1990). We therefore hypothesized that drowsiness may be responsible for decreased mobility and responsivity to sucrose-predictive cues in our SCH-treated rats. This hypothesis implies that fluctuations of arousal that alter animals’ wakefulness should similarly modulate mobility and DS responding and, therefore, these behavioral variables should co-vary in tandem with fluctuating arousal. Consequently, if the level of arousal fluctuates throughout the session, DS responding should be highly correlated with mobility in SCH-treated rats. In line with this possibility, even when the explicit arousing stimulus was not present at the beginning of the session, cross-animal mean mobility was highly variable, possibly due to arousing stimuli that were not under experimental control, and changes in mean DS response ratio closely paralleled changes in mean mobility (Fig. 7C1, compare the mobility in SCH to the dashed line representing DS response ratio). To further investigate the relationship between mobility and cue responding, we performed a correlation analysis using cross-animal mean mobility and DS responding data from 10 min time bins (Fig. 9; see Table S1 for full statistical results). Confirming our predictions, a high degree of correlation was observed in animals administered SCH, regardless of whether the initial arousing stimulus was present (Fig. 9A, E) or not (Fig. 9B, E). No correlation was observed when animals were administered vehicle, likely because their level of arousal was maintained above the threshold that allows for maximal responding to reward-predictive cues (Fig. 9E). In contrast to SCH-treated animals, we did not observe a significant correlation between mobility and DS responsivity in HAL-treated animals, even though their mobility was maintained below the threshold level throughout the session (Fig. 9C, E). These SCH and HAL results were recapitulated in cohort 2 (Fig. S5); animals in this cohort also showed a significant correlation between mobility and DS response ratio when given SCH but not HAL (Fig 9D,E). Our results show that SCH, by bringing arousal below the threshold, induces a coupling between mobility and DS responding that is not present in the control condition. On the other hand, HAL impacts either DS responding, or mobility, or both of these behavioral measures in an arousal-independent manner.

**Fig. 9.**
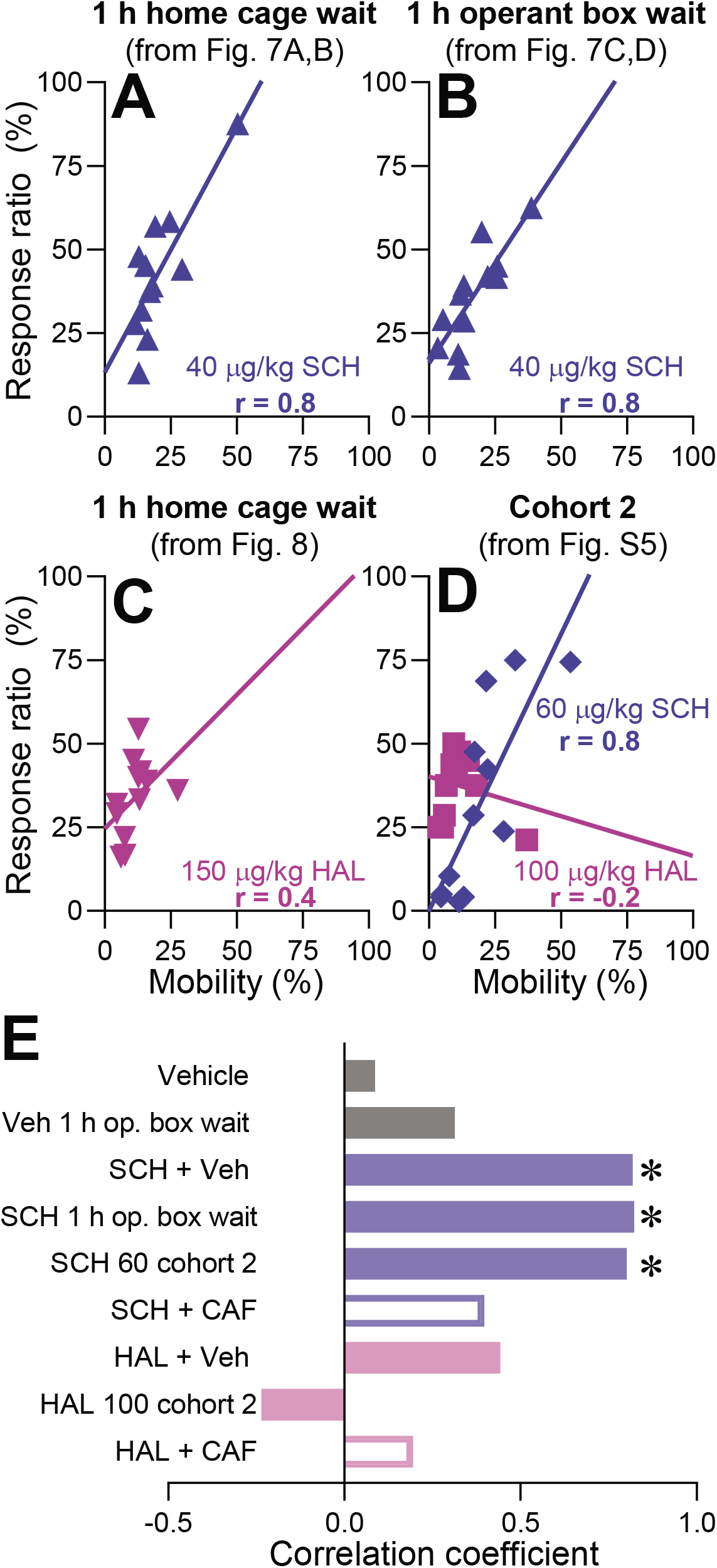
Mobility and DS response ratio are correlated after SCH23390 but not haloperidol injection. A-D, Correlation plots illustrating the dependence of the DS response probability on the probability of movement in each time bin in rats that received systemic SCH or HAL injections. Each data point represents the cross-subject mean mobility and corresponding mean DS response ratio from an individual time bin; the time course graphs from which the data are taken are indicated by the figure numbers in the plot title. The lines are linear regressions. R values represent Pearson correlation coefficients. E, Pearson correlation coefficients for the experiments presented in A-D as well as vehicle and caffeine experiments. Significance was assessed with t-tests for correlations; *, P < 0.05.

In summary, the correlation between mobility and cue responsivity that was observed only in rats administered SCH suggests that both these variables are strongly influenced by a common third variable, which we hypothesize to be arousal. In contrast, the absence of such a correlation after HAL administration suggests that HAL impairs mobility and cue responsivity via distinct mechanisms.

### Dopamine antagonists induce long immobile periods by decreasing the probability of movement initiation; the effect is reversed by CAF

Rodents’ exploratory behavior is characterized by brief periods of movement separated by periods of quiescence, which we will refer to as mobile and immobile periods. Analysis of lengths of these periods and how they are affected by experimental manipulations provides a more complete picture of the effects of the drugs on movement. Specifically, the decrease in mobility caused by both SCH and HAL (Figs. 7-9) raises the question of whether the drugs decreased the probability of movement initiation (which would reduce the number of movements and increase the intervals between them) or the length of movements once initiated, or both.

Both SCH and HAL increased the total time spent immobile (SCH: t_5_ = 3.7, P = 0.014; HAL: t_5_ = 7.9, P = 0.0005; Fig. 10A1), and in the case of HAL, these effects were prevented by CAF (t_5_ = 4.6, P = 0.0059, Fig. 10A1). CAF did not significantly attenuate the effects of SCH (t_5_ = 1.6, P = 0.17, Fig. 10A1) although some animals showed less immobile time in SCH+CAF than in SCH+vehicle (Fig. 10A1). Both dopamine antagonists reduced the total number of movements (mobile periods; SCH: t_5_ = 4.3, P = 0.008; HAL: t_5_ = 8.6, P = 0.0004; Fig. 10A2) which, together with an increase in the total time spent immobile, indicates that they impaired movement initiation. CAF abolished the effect of HAL on number of movements (t_5_ = 7.7, P = 0.0006, Fig. 10A2) and, although CAF did not significantly alter the effects of SCH (t_5_ = 1.6, P = 0.18), four of the six subjects showed more mobility periods in SCH+CAF than in SCH+vehicle (Fig. 10A2). Thus, SCH and HAL reduce the probability of movement initiation, whereas CAF counteracts these effects, particularly those of HAL.

**Figure 10.**
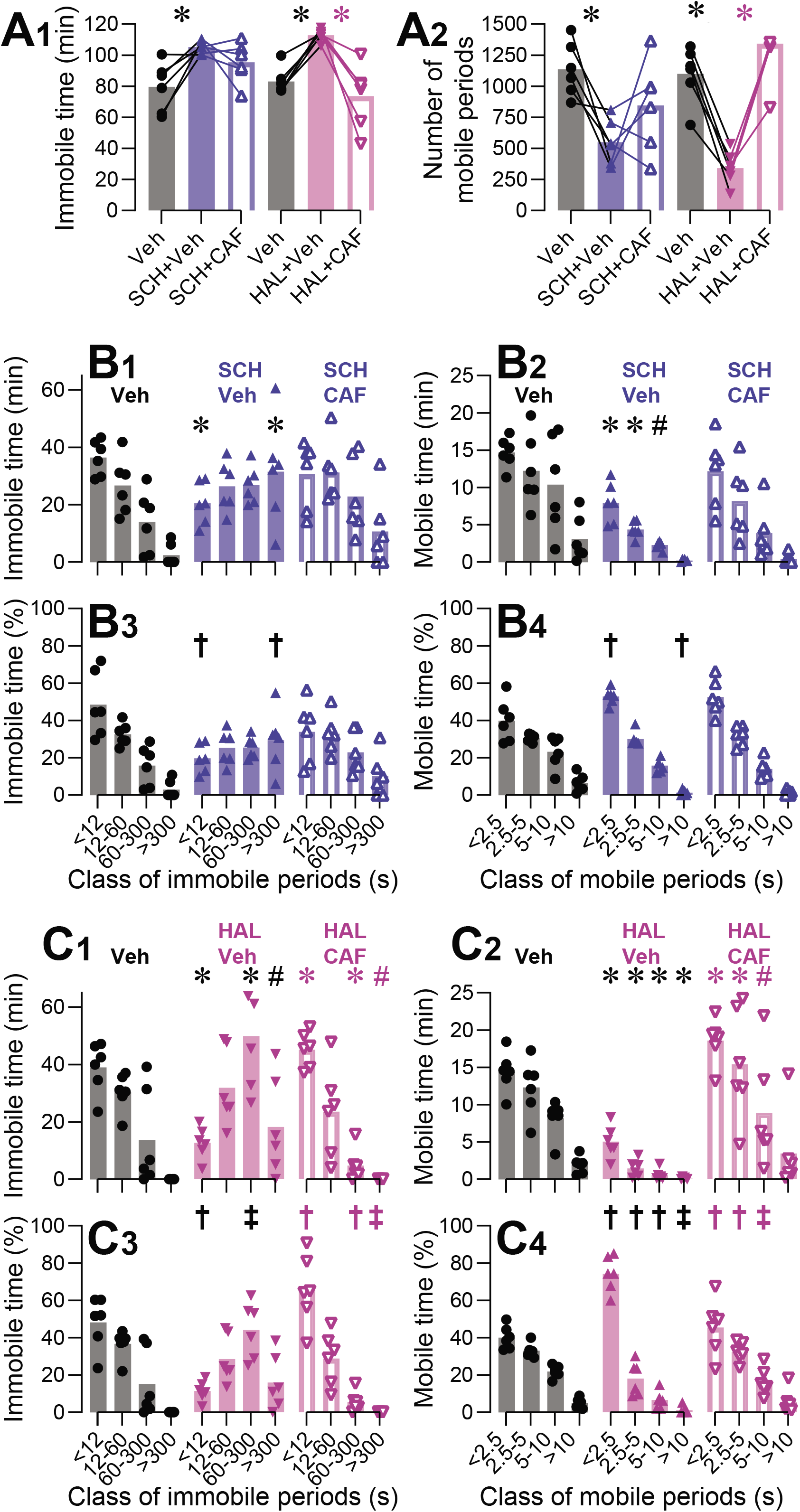
Impact of dopamine antagonists and caffeine on the distribution of mobility and immobility periods. A-C, Analysis of the distribution of immobility and mobility periods in 6 tracked rats from cohort 1 after administration of vehicle, SCH (40 µg/kg), SCH and CAF (40 µg/kg and 10 mg/kg), HAL (150 µg/kg), and HAL and CAF (150 µg/kg and 10 mg/kg). The data is from the experiments presented in Fig. 5 (the subset of subjects for which video tracking data was collected). All bars represent cross-subject means; symbols represent individual subjects. A, Total immobile time (A1) and the number of mobility periods (A2) registered throughout the entire operant session after administration of the indicated drugs. B,C, Total time (B1, B2, C1, C2), or percent of total time (B3, B4, C3, C4), spent in immobility (B1, B3, C1, C3) or mobility (B2, B4, C2, C4) periods belonging to the designated length classes indicated on the abscissa, after administration of the indicated drugs. A1,A2,B1,B2,C1,C2, Significance symbols indicate the results of paired t-tests vs vehicle for the SCH/Veh and HAL/Veh groups (black symbols) or vs the dopamine antagonist alone for the SCH/CAF and HAL/CAF groups (colored symbols). *, P < 0.05, remaining significant after adjustment for multiple comparisons; #, P < 0.05 but the result exceeded the critical P value adjusted for multiple comparisons; no symbol, P > 0.05. B3,B4,C3,C4, Significance symbols represent the results of permutation tests, which were not adjusted for multiple comparisons (see Methods, Statistical Analysis), vs vehicle for the SCH/Veh and HAL/Veh groups or vs the dopamine antagonist alone for the SCH/CAF and HAL/CAF groups. †, P = 0.016; ‡, P = 0.031; no symbol, P > 0.046.

While these results demonstrate that movement initiation is impaired by both dopamine antagonists, they do not exclude the possibility that the drugs may also impact maintenance of initiated movements. To further characterize these effects, we asked how the drugs influenced the distribution of lengths of immobile and mobile periods. Immobile periods were divided into four classes defined by a geometric sequence with the common ratio of 5: short (< 12 s), medium (12 – 60 s), long (1 – 5 min) and very long (> 5 min). Mobile periods were similarly classified, but because they were generally shorter than immobile periods, the first term of the sequence was set to 2.5 s, rather than 12 s. These choices resulted in similar shapes of the distributions of immobile and mobile periods in control animals (compare grey bars in Figs. 10B1 and 10B2, and in 10C1 and 10C2). Although in the control (vehicle) condition animals spent most of their time throughout the entire session immobile (mean ± SEM of 79.6 ± 6.4 min out of the 120 min session, grey bar in Fig. 10A1), this immobility was mostly accounted for by periods of short and medium length; no immobile periods longer than 5 min were observed in most vehicle sessions (Fig. 10B1,C1). In stark contrast, animals administered SCH spent most of their immobile time in very long immobile periods (SCH vs vehicle, >300 s: t_5_ = 4.3, P = 0.007, Fig. 10B1), while the time spent in short immobile periods was significantly decreased compared to vehicle (<12 s: t_5_ = 4.2, P = 0.009; 12-60 s: t_5_ = 0.06, P = 0.96; 60-300 s: t_5_ = 1.9, P = 0.12; Fig. 10B1). The decrease in mobile time induced by SCH was observed across all classes of movement lengths such that the shape of the distribution was largely unaffected compared to vehicle (<2.5 s: t_5_ = 4.0, P = 0.01.; 2.5-5 s: t_5_ = 3.9, P = 0.01; 5-10 s: t_5_ = 3.0, P = 0.03; >10 s: t_5_ =2.5, P = 0.06; Fig. 10B2). This difference between the effects of SCH on immobile and mobile periods is most clearly seen when the data are normalized as the percentage of the total immobile time (Fig. 8B3) or mobile time (Fig. 8B4) accounted for by periods from each class of length: a major change of the distribution of immobile periods is observed, while the distribution of mobile periods is only slightly shifted towards the shortest lengths. CAF appeared to partially normalize the shift in distribution of immobile periods, but this effect did not reach statistical significance for either non-normalized data (SCH+CAF vs SCH, <12 s: t_5_ = 1.5, P = 0.19; 12-60 s: t_5_ = 0.86, P = 0.43; 60-300 s: t_5_ = 0.56, P = 0.60; >300 s: t_5_ = 2.2, P = 0.084; Fig. 10B1) or normalized data (see figure legend for Fig. 10B3). CAF did not affect the non-normalized (<2.5 s: t_5_ = 1.6, P = 0.16; 2.5-5 s: t_5_ = 1.8, P = 0.13; 5-10 s: t_5_ = 1.0, P = 0.37; >10 s: t_5_ = 0.98, P = 0.37; Fig. 10B2) or normalized (Fig. 10B4) distribution of mobile periods. HAL similarly decreased the time spent in short immobile periods. However, most of the immobility in rats administered HAL is accounted for by the long 1 – 5 min, rather than very long (>5 min), immobile periods, which is apparent in both non-normalized data (HAL vs vehicle, <12 s: t_5_ = 6.7, P = 0.001; 12-60 s: t_5_ = 0.26, P = 0.81; 60-300 s: t_5_ = 3.7, P = 0.014; >300 s: t_5_ = 2.6, P = 0.046; Fig. 10C1) and normalized data (Fig. 10C3). HAL also decreased the time spent in movements of all length classes (<2.5 s: t_5_ = 9.2, P = 0.0003; 2.5-5 s: t_5_ = 6.3, P = 0.0014; 5-10 s: t_5_ = 7.5, P = 0.0006; >10 s: t_5_ = 3.8, P = 0.012; Fig. 10C2), but the percentage of mobile time spent in short movements was enhanced compared to vehicle (Fig. 10C4). All these effects in both non-normalized data (HAL+CAF vs HAL+vehicle, immobile periods: <12 s: t_5_ = 13.4, P = 0.00004; 12-60 s: t_5_ = 1.3, P = 0.25; 60-300 s: t_5_ = 6.0, P = 0.0019; >300 s: t_5_ = 2.6, P = 0.046; Fig. 10C1) (HAL+CAF vs HAL+vehicle, mobile periods: <2.5 s: t_5_ = 11.4, P = 0.00009; 2.5-5 s: t_5_ = 4.8, P = 0.0048; 5-10 s: t_5_ = 2.8, P = 0.038; >10 s: t_5_ = 1.5, P = 0.18; Fig. 10C2) and normalized data (Fig. 10C3,C4) were reversed by CAF.

In summary, the observed shift of the distribution of immobile periods towards longer lengths induced by either dopamine antagonist indicates that both drugs impaired the ability to initiate movement. On the other hand, the shift of the distribution of mobile periods towards the shortest lengths was much more pronounced in HAL-treated rats, indicating that HAL, but not SCH, impaired the ability to maintain movements once initiated.

### Differential impact of SCH and HAL on movement maintenance and signs of drowsiness

Both SCH and HAL induced very long immobile periods (> 5 min) that were almost never observed in rats administered vehicle (Figs. 10B1,C1, S6B1). Such long immobile periods are compatible with episodes of sleep, but they can also occur as a manifestation of motor impairment or catalepsy. To gain insight into the possible causes of this prolonged immobility, two observers blind to drug treatment (SCH or HAL) scored video recordings for the presence of postures that are characteristic of rats’ natural sleeping positions. This analysis utilized the cohort 2 animals for which video tracking was available; the effects of 60 µg/kg SCH and 100 µg/kg HAL on mobility and immobility periods in these subjects were similar to the effects of these drugs in cohort 1 (Fig. S6). The observers examined videos of these sessions to identify instances of two natural sleep postures, “curled-up” and “ball” (van Betteray et al., 1991) (Fig. 11A, Supplementary Video 2). While other postures, such as “stretch-out” and “sit”, can also indicate sleep in rats (Coenen et al., 1983; van Betteray et al., 1991), they are also compatible with prolonged wake immobility and difficult to dissociate from motor freezing or catalepsy, and therefore they were not included as a criterion. The observers were also told to pay attention to behaviors that occur before immobility periods: if immobility is preceded by behavior that resembles searching for the most comfortable position, it is more likely to represent sleep than immobility occurring suddenly in the middle of locomotion or grooming. We will refer to this phenomenon as “comfortable position-seeking behavior”.

**Figure 11.**
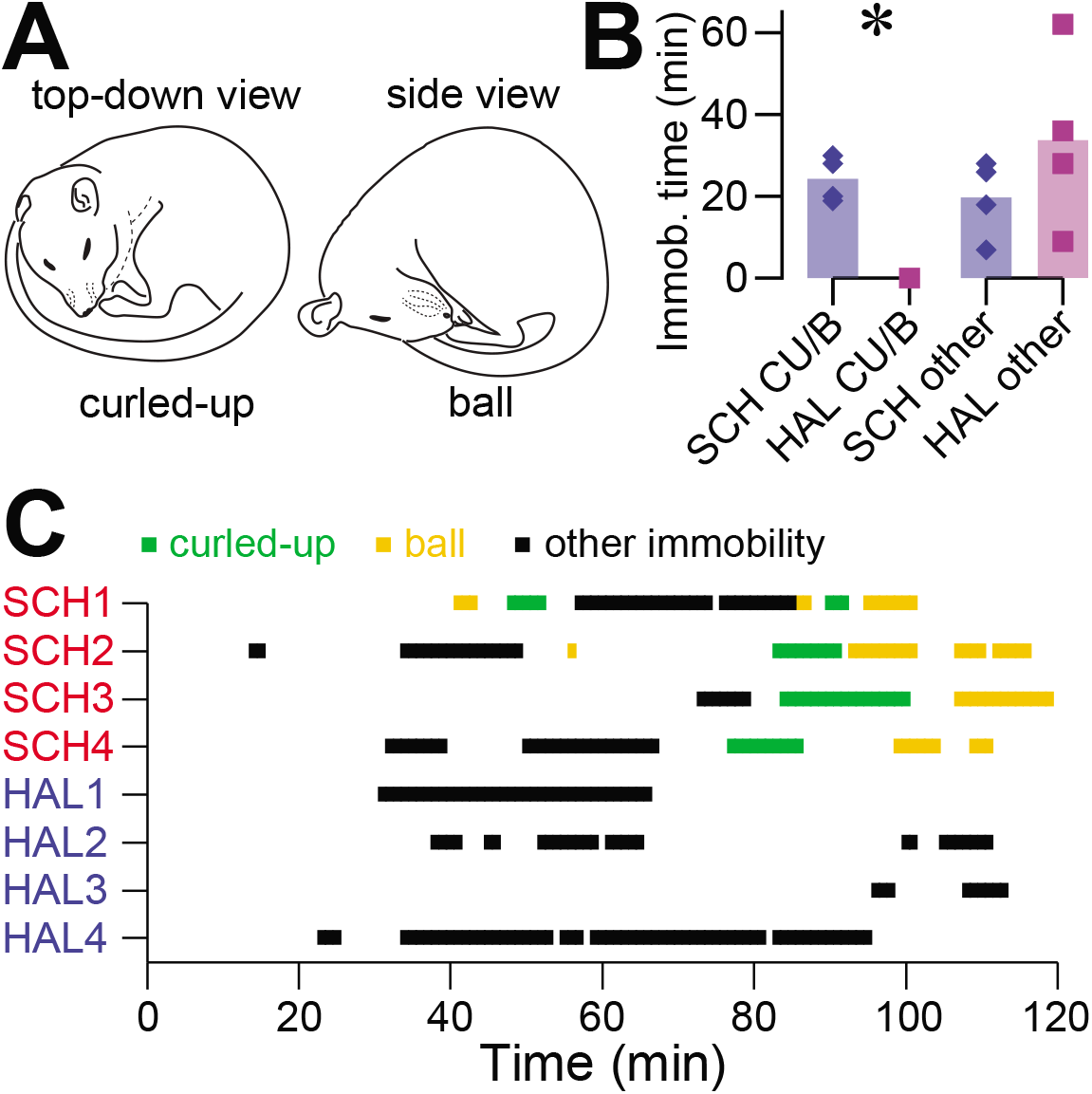
Sleep posture analysis. A, Curled-up and ball positions that are characteristic of sleep in rats. B, The total time spent immobile in sleep-characteristic curled up and ball (“CU/B”) and rigid (“Other”) positions in cohort 2 rats administered 60 µg/kg SCH and 100 µg/kg HAL. C, The distribution of long immobility periods of all three types across the operant session. *, P < 0.05, unpaired t-test.

Based on the occurrence of curled-up and ball positions, and of comfortable position-seeking behavior, both observers identified presumed episodes of sleep in four out of eight rats. All of these turned out to be the animals that received SCH. No obvious signs of sleep were observed in HAL-treated rats. The likelihood that the subgroup of four SCH-treated rats happened to be identified as satisfying the criteria by chance is equal to 1/70 = 0.014 for each observer, and the likelihood of both independent observers performing successful identification by chance is equal to (1/70)^2^ = 0.0002 (see Methods, Statistical Analysis). Therefore, the null hypothesis that there is no difference in the signs of sleepiness between SCH and HAL-treated rats can be rejected.

In a second analysis, an observer was asked to specifically identify the times during which presumed episodes of sleep occurred, identified based on the criteria described above, as well as to identify other immobile periods that do not satisfy the criteria and, therefore, are less likely to represent sleep. For this analysis, only immobile periods longer than 1 min were taken into account. Since very few (in most subjects, the number was zero) immobile periods longer that 1 min occurred in vehicle-treated rats (Fig. S6B1), the analysis was restricted to the animals administered dopamine antagonists. Consistent with the previous finding, only SCH-treated rats spent time in presumed sleep postures, while immobility that is less likely to represent sleep was observed in both groups (unpaired t-test comparison of SCH vs HAL, immobile time spent in sleep postures: t_6_ = 9.7, P = 0.0001; immobile time spent in other postures: t_6_ = 1.2, P = 0.29; Fig. 11B). Therefore, the video analysis suggests that SCH induces sleep much more potently than HAL, which is in agreement with previous research (Trampus and Ongini, 1990). This finding provides another argument in favor of the hypothesis that SCH impairs responding to reward-predictive cues by directly decreasing arousal. While curled-up and ball positions characteristic of sleep were observed mostly in the second hour in SCH-treated rats (Fig. 11C), it is possible that immobility occurring earlier in the session was caused by sleeping in less typical postures, or by the decreased arousal state that precedes sleep. On the other hand, because HAL-treated rats did not show any obvious signs of sleep, it is more likely that their immobility represented catalepsy. Importantly, the doses of SCH and HAL used equivalently affected cued approach behavior (Fig. 6) and total time spent immobile (Fig. S6A) and therefore the differences in the effects of these drugs on sleep are not due to differences in dose.

## Discussion

In this study, we examined the inhibition of the cued pursuit of sucrose reward caused by intra-NAc and systemic administration of the D1 antagonist SCH, systemic administration of the D2 antagonist HAL, and the effects of co-administration of the adenosine antagonist CAF. We conclude that the impaired pursuit of rewards in our animals cannot be ascribed to reward-related functions such as reinforcement, motivation, liking, or wanting. This conclusion holds despite the diversity of interpretations and definitions that some of these terms have acquired (Nicola, 2016), some of which will be discussed in the following subsections. Instead, responding to cues by our rats administered dopamine antagonists was hampered by drowsiness and catalepsy. To justify this claim, we argue, first, that these results could not have been due to impaired reinforcement or liking for the following reasons. In the case of HAL administration, animals exhibited decreased responding to cues from the very beginning of the session, in agreement with other studies (Fibiger et al., 1976; Phillips and Fibiger, 1979), rather than a gradual decline; there was no need for the animals to consume rewards under the influence of the drug for their responding to decrease. In the case of SCH administration, gradual extinction-like declines of responding were observed; however, these reflected declining arousal and not extinction because the effects of the drug were immediate when the high initial arousal state caused by transferring animals from their home cage to the operant chamber was eliminated. These arguments rule out impaired reinforcement or liking as a cause of decreased cue responding. Furthermore, our analysis of the lengths of receptacle entries is incompatible with explanations based on other reward-related deficits such as decreased motivation or wanting. Specifically, it is hard to imagine why *less* motivated animals, or animals that want the rewards less, would stay in the receptacle for extended periods of time. Since long periods of immobility were also observed outside the receptacle, this phenomenon is most parsimoniously explained by sleepiness and/or catalepsy. Moreover, our finding that HAL shortened movements is consistent with drug-induced Parkinsonism and cannot be explained by reward-specific effects of the drug. Although we cannot entirely rule out the possibility that dopamine antagonists affected reward-specific functions, we did not obtain any evidence to support such a claim. On the other hand, the effects of these drugs on arousal and motor function were evident and sufficient to explain all observed impacts of the drugs on behavior. It is, therefore, most parsimonious to conclude that decreased responsiveness to the reward-predictive cues was a secondary consequence of more general performance problems.

Our findings imply that the behavioral cause of impaired pursuit of rewards in a particular operant paradigm can only be identified with a comprehensive analysis and control experiments that are rarely performed. The analysis should ideally address the following questions: Is the time course of the impairment immediate or gradual? If gradual, is the delay due to pharmacodynamics, an extinction-like phenomenon, declining arousal, progressive satiety, physical fatigue, or another time-dependent variable? Do animals show deficits that are not directly linked to rewards but might nevertheless impact their pursuit? If so, can these deficits *entirely explain* the behavior? In particular, is the analysis comprehensive enough to detect signs of drowsiness and/or catalepsy?

While both SCH and HAL impaired cued pursuit of sucrose by inducing performance problems, the effect of external arousing stimuli on animals administered these two drugs were very distinct. In the following subsection, we will discuss the observed differences between the effects of SCH and HAL and hypothesize on the mechanisms that might be responsible for them.

### Differential behavioral effects of SCH and HAL

Our study identified four differences between the effects of SCH and HAL:

1. Animals administered SCH showed more overt signs of sleepiness, as evidenced by assuming sleep-characteristic positions after periods of comfortable position seeking. On the other hand, animals administered HAL froze in seemingly accidental positions without any obvious preparatory activities.
2. While both drugs lengthened periods of immobility, only HAL shortened movements.
3. Deficits in responding to sucrose-predictive cues caused by SCH, but not HAL, were readily alleviated by arousal induced by either transfer to the operant box (manifested as initial resistance to the effects of SCH followed by gradually increasing impairment in responding) or handling.
4. Mobility and responsiveness to the cues were correlated throughout the session in rats administered SCH, but not HAL.

Difference 1 is compatible with previous studies that found SCH more effective in inducing both total sleep and REM sleep than HAL (Ongini et al., 1993; Trampus and Ongini, 1990). Because even small doses of SCH (0.003-0.03 mg/kg SC) were found to promote sleep in these studies, drowsiness should always be considered as a potential contributor to the behavioral effects of SCH, even when no overt signs of sleepiness are observed.

Difference 2 indicates the presence of motor impairment in HAL-treated animals. While both drugs caused periods of freezing in the receptacle that likely indicate a certain degree of catalepsy, the shortening of movements observed exclusively in HAL-administered animals suggests a more severe and multifactorial motor impairment. The shortening of movements that we observed may have similar origin to the shortening of feeding bouts observed by others in HAL-administered rats (Salamone, 1988).

Difference 3 may seem to follow from 1) and 2): SCH-induced drowsiness and all its associated secondary consequences, such as decreased mobility and cue responsivity, should be alleviated by manipulations that increase arousal, whereas the more complicated motor deficits induced by HAL may be insensitive to these manipulations. However, the mobility of HAL-administered rats was elevated at the beginning of operant sessions for both cohorts of rats (Figs. 8A1, S5A), suggesting some degree of alleviation of their motor problems by the initial arousal. This apparent motor improvement was not accompanied by increased cue responsivity, which is particularly evident for cohort 2 whose cue responding was lowest in the first 10 min bin while their mobility was highest (Fig. S5). One possible explanation of this finding is that arousing conditions alleviated HAL-induced catalepsy, but other aspects of motor function remained compromised and prevented the animals from collecting rewards. Another possibility is that sucrose seeking in HAL-treated rats was hampered not only by their motor problems but also other HAL-induced deficits that cannot be alleviated by arousing stimuli, such as anxiety or akathisia, a known side effect of HAL that is manifested by the subjective feeling and behavioral expression of restlessness and inability to sit still (Van Putten et al., 1984). As akathisia can develop within two hours after the first oral dose of HAL in some patients (Van Putten et al., 1984), it is plausible that it can emerge faster after IP injection and contribute to the phenotype observed in our rats who were placed in the operant chamber 1 h after injection. Although studying akathisia in animals is extremely challenging because neuroleptic-induced dyskinesia masks behavioral manifestations of restlessness and the subjective feeling of restlessness cannot be directly assessed (Sachdev and Brune, 2000), the widespread occurrence of akathisia as a side effect of HAL in human patients (Van Putten et al., 1984) and certain rodent studies (Sachdev and Saharov, 1997) suggest that it may contribute to the experience of our HAL-treated rats and steer them away from sucrose consumption, even when their dyskinesia transiently improves. Finally, we cannot exclude the possibility that HAL, in addition to causing motor and possibly other performance problems, also decreased the perceived value of sucrose rewards, motivation to pursue them, or the ability to respond to predictive cues. Reward-specific deficits could be masked by co-occurring severe performance problems but could nevertheless explain why transient alleviation of dyskinesia does not result in increased sucrose consumption in rats administered HAL. The dissociation between dyskinesia and impaired cue responsiveness, regardless of the specific reason, gives rise to the lack of correlation between these behavioral variables in rats administered HAL (difference 4 in the list above).

Although arousing stimuli failed to improve responding to sucrose-predictive cues in HAL-administered animals performing our task, the fact that their dyskinesia was alleviated suggests that the effects of neuroleptic drugs on performance of other operant tasks might be more sensitive than our task to manipulations that alter arousal. In particular, we argued in the Introduction that within-session declines of performance observed in neuroleptic-administered animals when the reward rate is high, such as in the FR1 task, are more likely to reflect decreasing arousal than extinction. These declines usually occur within the initial few minutes of the FR1 task. Our cued VI50s task that involved randomly interspersed reward-predictive and neutral stimuli was not suitable to detect such fast declines because the expected time interval between reward-predictive cues was close to 2 min and sometimes, when a few neutral stimuli occurred in sequence, no reward-predictive cue occurred for many minutes. Therefore, we cannot exclude the possibility that arousing stimuli could transiently improve operant performance in animals with antagonized D2 receptors, but our results imply that such improvement should be short-lived. This conclusion should be contrasted with the slow and pronounced within-session declines of performance spanning approximately 1 h that we observed in animals administered SCH. These slow declines are consistent with the hypothesis that antagonizing D1 receptors disrupts performance in our task by making the animals sleepy and the transfer to the operant chamber improved their performance by simply waking them up.

On the other hand, a very brief improvement of motor function in HAL-treated animals may reflect paradoxical kinesia, which has been observed in rats administered HAL (Clark et al., 2009; Melo-Thomas and Thomas, 2015; Tonelli et al., 2018a, b) and described as a brief improvement of their motor function by an arousing stimulus followed by recurrence of motor impairment soon after the stimulus is discontinued (Tonelli et al., 2018a). Importantly, we do not mean to claim that SCH did not induce any kind of motor impairment in our rats. The periods of freezing in the receptacle that we observed after SCH administration are more likely to represent catalepsy than sleep, in agreement with other reports of SCH-induced catalepsy (Meller et al., 1985; Morelli and Di Chiara, 1985). However, the degree of catalepsy induced by the dose of SCH used in this study was small enough to allow animals to respond efficiently to cues when their level of arousal was high, which might be at least in part due to the alleviation of SCH-induced catalepsy by arousal. While the session progressed, the natural drop of arousal augmented by SCH led to the increase in the length of immobility periods, which could have represented a combination of catalepsy and microsleeps (Friedman et al., 1979; Levitt, 1967; Tirunahari et al., 2003; Vyazovskiy et al., 2011), and ultimately to the episodes of genuine sleep preceded by assuming sleep-characteristic positions.

### Potential neural mechanism explaining the different behavioral effects of SCH and HAL

In this study, we increased the level of arousal by transferring the rats from their home cage to the operant chamber or by opening the chamber door in the middle of the session and touching them. Both procedures involved handling, which by itself provides a powerful arousing stimulus with a broad impact on brain activity. Handling rats triggers epinephrine and norepinephrine release into the bloodstream (Ondicova et al., 2019), increases hippocampal and cortical acetylcholine (Inglis and Fibiger, 1995), hippocampal norepinephrine and serotonin (Kalen et al., 1989), cortical histamine (Westerink et al., 2002) and stimulates the release of norepinephrine in the locus ceruleus and both norepinephrine and dopamine in the medial prefrontal cortex (Kawahara et al., 1999). Finally, handling rats was found to increase dopamine levels in multiple dopamine terminal regions, including the basolateral amygdala, the prefrontal cortex, and the NAc (Inglis and Moghaddam, 1999). These increases are at least in part due to the activation of glutamatergic inputs onto VTA dopamine neurons (Enrico et al., 1998). We hypothesize that increased dopamine plays a major role in the restoration of efficient operant performance in our SCH-administered rats after arousing stimulation. In this subsection, we first argue that this hypothesis explains the difference between the effects of natural arousing stimuli on rats administered SCH and HAL, and then we propose that the hypothesis also explains the indiscriminate ability of CAF to counteract the impairments induced by both SCH and HAL.

While the deficits elicited by dopamine antagonists likely involved multiple brain systems, all these deficits originally resulted from inhibited dopamine transmission. Therefore, increased release of dopamine can be expected to reduce the impairment, assuming that dopamine receptors are only partially blocked and/or there is some degree of functional redundancy between different classes of dopamine receptors such that increased activity of the receptors that are not targeted by the drug can be compensatory. Because our dose of SCH was low compared to the dose-response range typically used in catalepsy studies, a significant percentage of D1 receptors almost certainly remained functional. Therefore, the arousal-induced increase in striatal dopamine was detected by D2 receptors and by the D1 receptors that were not blocked by SCH. As both D1 and D2 receptor-expressing striatal neurons regulate the level of wakefulness (Luo et al., 2018; Oishi et al., 2017b), both of these effects likely contributed to the alleviation of SCH-induced drowsiness. Enhanced activation of D2 receptors might have been particularly important for the alleviation of SCH-induced motor problems as it has been shown that D2 agonists ameliorate catalepsy in SCH-administered rats (Meller et al., 1985; Morelli and Di Chiara, 1985). Prolonged improved performance after arousing stimulation can be maintained by increased ambient dopamine levels that are detected by high affinity D2 receptors (Goto et al., 2007; Grace, 1991; Richfield et al., 1989; Schultz, 1998), but also by persistent changes in the excitability of D1-expressing MSNs triggered by the initial robust phasic activity of dopamine neurons (Lahiri and Bevan, 2020). The relative role of D2 receptors and unblocked D1 receptors in the restoration of distinct aspects of performance could be tested by examining the effects of D1 and D2 agonists on cued sucrose seeking and mobility in rats administered SCH.

Assuming we are correct that arousing stimuli cause performance improvements in SCH-treated rats via increased dopamine release, why was a similar improvement not observed in HAL-treated animals? Many differences between the two families of dopamine receptors could have contributed to this observation. First, D1 and D2 receptors are differentially involved in the regulation of prefrontal and limbic afferents to the NAc (Goto and Grace, 2005). Second, based on the fact that D1 and D2 receptor families tend to occupy states that differ in their affinity for dopamine (Richfield et al., 1989), it has been hypothesized that high-affinity D2 receptors are well suited to detect changes in the ambient dopamine levels, whereas low-affinity D1 receptors are mostly activated by higher dopamine levels produced by phasic activity of dopamine neurons (Goto et al., 2007; Grace, 1991; Schultz, 1998), although this hypothesis has been questioned (Hunger et al., 2020). Third, striatal cholinergic interneurons respond to salient stimuli (Schulz and Reynolds, 2013), at least in part due to the action of dopamine on D2 receptors expressed by cholinergic interneurons (Straub et al., 2014), and these D2 receptors were recently implicated in HAL-induced catalepsy (Kharkwal et al., 2016). Fourth, D2 receptors are expressed on the cell bodies and terminals of dopamine neurons serving an inhibitory auto-receptor function (Ford, 2014). While all these differences might have contributed to our results, we argue that the inhibitory auto-receptor function of D2 receptors may be sufficient to explain why arousal-induced increases in dopamine did not restore performance in rats administered HAL. By blocking this auto-receptor function, HAL increases the activity of dopamine neurons and striatal dopamine levels (Blaha and Lane, 1984; Bunney and Grace, 1978; Bunney et al., 1980; Bunney et al., 1973; Imperato and Di Chiara, 1985; Lidsky and Banerjee, 1993). Therefore, administration of a D2 antagonist such as HAL activates two opposing processes: decreased availability of functional D2 receptors is compensated for by increased baseline level of dopamine. At low doses of HAL, this increased dopamine activity overcompensates for decreased receptor availability and, therefore, the effects of low doses of HAL (40 µg/kg) are stimulating (Dias et al., 2012). For higher doses, the proportion of functional D2 receptors is too small to compensate for those that are antagonized, despite increased dopamine levels, resulting in akinesia, catalepsy, and other motor deficits. The doses of HAL used in our study (100, 150 and 200 µg/kg) were chosen so that the effects of HAL on cue responding and mobility were comparable to those of SCH when averaged over the entire session. For these high doses, the block of D2 striatal receptors was widespread enough to inhibit locomotion and task performance despite enhanced dopaminergic activity.

With this hypothesis, it is straightforward to explain why the mechanism of reactivation of performance by arousing stimuli that we proposed for SCH-administered rats could not be effective in HAL-administered animals: because the activity of dopamine neurons and dopamine levels are already high in the presence of HAL, any neural mechanism that depends on increasing them further has limited impact. In fact, because the degree of HAL-induced increase in dopamine neuron activity is high for neurons with low baseline firing and small for those with high baseline firing (Bunney et al., 1973), it has been suggested that HAL can bring the baseline firing rate of dopamine neurons close to the limit that these neurons can consistently maintain (Bunney and Grace, 1978). Moreover, even if dopamine neuron firing could be further increased, it would have little impact on dopamine levels because they are already elevated due to the increase in both baseline dopamine neuron activity and terminal dopamine release caused by the antagonism of somatodentritic and terminal auto-receptors, respectively. Finally, a very limited increase in ambient dopamine that might still be triggered by arousing stimulation under these conditions would have little impact on striatal processing because a significant percentage of D2 receptors is inactivated. While D1 receptors could in principle register this small dopamine increase, it is unlikely to have much impact on striatal neural activity because D1-expressing striatal medium spiny neurons are potently inhibited by D2-expressing neurons in the presence of a D2 antagonist (Dobbs et al., 2016) and a small relative increase in D1 receptor activity is unlikely to release D1-expressing neurons from this lateral inhibition. Thus, in the presence of HAL but not SCH, there exist several strong and likely insurmountable obstacles to the ability of arousing stimuli to reactivate behavioral performance.

All of these obstacles are circumvented by using CAF in place of natural arousing stimulation. Because adenosine and dopamine have long been hypothesized to have opposing impact on the physiology of striatal neurons (Ferre et al., 1997), antagonists of adenosine and dopamine receptors can be expected to have opposing impacts as well. According to this idea, CAF is likely to compensate for the disruptive effects of HAL on striatal medium spiny neurons via a dopamine-independent process: by antagonizing adenosine 2A receptors on D2 receptor-expressing striatal neurons, CAF is likely to partially normalize their function. If natural arousing stimuli could decrease the levels of extracellular adenosine, or antagonize adenosine receptors, they would likely have alleviated the effects of HAL on operant performance. However, although arousing stimuli activate diverse neurophysiological processes, there is no known arousal-driven neural mechanism that would decrease the activity of adenosine receptors: the clearance of adenosine from extracellular space is driven mostly by homeostatic mechanisms (Fredholm et al., 2005); no endogenous antagonists of adenosine receptors have been identified; and salient events do not alter the adenosine level in the NAc (Nagel and Hauber, 2002). Moreover, while natural arousing stimuli likely activate dopamine-independent neurophysiological processes, our results indicate that none of them is effective in restoring the performance of HAL-administered animals in our task. It remains to be explained why the effects of HAL on mobility were briefly alleviated at the beginning of the session, even though the emergence and detection of the arousal-related dopamine increase was compromised according to our hypothesis. We hypothesized above that this effect might have been due to paradoxical kinesia, which does not require increased dopamine (Keefe et al., 1989) and may depend on inferior collicular activity (Melo-Thomas and Thomas, 2015; Tonelli et al., 2018b; Tostes et al., 2013), although the circuitry that triggers paradoxical kinesia in our rats was likely broader due to the polymodal nature of our arousing sensory stimulation.

We argued above that the differences in the effects of natural arousal on performance of rats administered SCH and HAL are compatible with a simple hypothesis motivated by the observation that D2, but not D1, receptors serve as inhibitory auto-receptors. While our auto-receptor hypothesis certainly does not provide a complete picture, it nevertheless provides a coherent explanation for our observed differences in the effects of SCH and HAL. Moreover, this hypothesis can be further experimentally challenged. One possible experiment could aim to restore performance in HAL-administered animals by optogenetic activation of dopamine neurons at the time of arousing stimulation. However, if the level of activity of dopamine neurons is already increased as a consequence of blocked autoinhibition, the efficacy of optogenetic activation might be compromised. A more direct approach to test our hypothesis would utilize optogenetic inhibition of dopamine neurons as a substitute for impaired autoinhibition. A lower dose of HAL should then be sufficient to impair performance because optogenetic inhibition would prevent the increase in dopamine release that otherwise compensates for partial antagonism of D2 receptors. Moreover, a pause in optogenetic inhibition at the time of arousing stimulation (e.g. at the beginning of the session or during mid-session handling) should trigger an increase in dopamine that was otherwise missing and allow cue responding to be restored by the stimulation.

### The effect of CAF and its implications for the necessity of phasic dopamine signaling for cued approach

CAF is believed to exert its stimulant actions by antagonizing adenosine A1 and A2A receptors (Fredholm, 1995). Based on the expression patterns of these receptors and G-proteins to which they are coupled, it has been hypothesized that adenosine and dopamine have opposing impacts on striatal neurons (Ferre et al., 1997). In agreement with this hypothesis, CAF and other antagonists of adenosine receptors can alleviate symptoms of Parkinson’s disease (Prediger, 2010), as well as reverse catalepsy (Acuna-Lizama et al., 2013; Hauber et al., 1998; Hauber et al., 2001; Kanda et al., 1998), hypophagia (Kim and Palmiter, 2003), or deficits in operant performance (Farrar et al., 2007; Mott et al., 2009; Robinson et al., 2005; Salamone et al., 2009; Worden et al., 2009) in animals with disrupted dopaminergic function.

On that account, our finding that CAF can reverse the effects of SCH and HAL on both mobility and operant performance agrees with previous studies. However, operant performance in our task explicitly depended on animals’ ability to respond to the reward predictive cues. The fact that CAF restored cue responding in animals administered dopamine antagonists argues against the importance of temporally precise phasic dopaminergic signaling for this form of behavior. Pharmacological antagonism of adenosine receptors could conceivably reverse tonic deficits induced by pharmacological blockade of dopamine receptors on striatal neurons, such as changes in excitability, responsivity to excitatory and inhibitory inputs, or the probability of vesicular GABA release. However, pharmacological blockade of adenosine receptors is unlikely to restore temporally precise dopaminergic signaling. Regardless of the compensatory actions of CAF downstream, blocked dopamine receptors cannot detect changes in dopamine level and the ability of striatal neurons to respond to these changes remains compromised. Thus, the CAF rescue effect argues against a requirement for phasic cue-evoked dopamine release for the approach response to the cue in our task.

This conclusion may seem at odds with multiple studies reporting robust release of striatal dopamine in response to reward-predictive cues (Day et al., 2007; Roitman et al., 2004) or ramping dopamine signals preceding reward consumption (Collins et al., 2016; Howe et al., 2013; Kim et al., 2020; Phillips et al., 2003; Roitman et al., 2004). However, these studies do not examine the necessity of these dopamine signals for behavior. While stimulation of VTA dopamine neurons was found to decrease response latency (Hamid et al., 2016), trigger movements (Barter et al., 2015; da Silva et al., 2018; Howe and Dombeck, 2016), and promote lever pressing during post-reinforcement pauses in rats self-administering cocaine (Phillips et al., 2003), all these findings show that dopamine increases may facilitate or invigorate operant behavior under certain conditions but do not demonstrate the necessity of endogenous phasic dopamine signals for behavior. An argument in favor of the necessity of dopamine signals follows from a study in which optogenetic inhibition of VTA dopamine neurons at different time epochs of the FR8 task was shown to decrease the probability of initiation and maintenance of bouts of nose-poking (Fischbach-Weiss et al., 2018). However, the fact that optogenetic inhibitions were relatively long (15 s) and not precisely timed to disrupt expected phasic dopamine signals makes it difficult to determine whether performance was impaired due to the elimination of phasic dopaminergic signaling or rather due to transient decreases in dopaminergic tone.

In sum, no conclusive evidence has yet been given that phasic dopamine signals are necessary for ongoing behavior, although such signals may be required during reward delivery to maintain future responding (Lee et al., 2020). It has been noted by others that the ability to reverse the deficits resulting from dopamine disruptions with systemic dopamine agonists or dopamine precursors undermines the interpretation of these deficits as resulting from disrupted phasic dopamine signaling (Niv et al., 2007; Schultz, 1998). Accordingly, we previously observed that administrations of dopamine agonists into the NAc improve performance of our cued sucrose-seeking task, even though these administrations increase the ambient level of dopamine receptor activity but likely blunt temporally-specific signaling due to partial saturation of the receptors (du Hoffmann and Nicola, 2016). Our current finding that cue responding of dopamine-compromised rats is so efficiently restored by CAF further argues in favor of the hypothesis that cue-evoked dopamine signals are not necessary for the performance of a learned cued reward-seeking task. However, an alternative interpretation of our result is that temporally-precise dopaminergic signaling was only partially attenuated and, once tonic effects of the antagonists were compensated for by CAF, this attenuated signaling was sufficient to drive responding to the cues.

### Relationships among arousal, motivation, motor dysfunction, resource allocation and dopamine

The concept of arousal has been a central one in psychology for nearly a century and has been used to describe the degree to which an organism is “activated”, or the intensity of behaviors regardless of their direction, or the degree of internal activation, which may not be manifested behaviorally if inhibitory processes are at play (Duffy, 1957). More recently, arousal has been regarded as one of the main components of feelings, emotions, and affect. According to an influential theory by Russell and Barrett, at the heart of every emotion lies the core affect, an elementary consciously accessible emotional experience that may but does not have to be directly related to any specific stimulus, object, or action (Russell and Barrett, 1999). Core affect can be thought of as a raw feeling that can be consciously assessed at each instant and represented by a point in a two-dimensional space, one dimension representing the degree of pleasure/displeasure (also referred to as hedonic valence) and the other representing the level of arousal. The arousal dimension represents a continuum ranging from sleep, through drowsiness, increasing stages of alertness, up to frenetic excitement (Russell, 2003).

The common feature of the definitions of arousal employed by different authors is its lack of directionality. This distinguishes arousal from attention and motivation. Attention is necessarily linked to a stimulus that is being attended to and it is usually defined as the amount of brain processing power allocated to the attended stimulus (Coull, 1998). The amount of processing power depends on arousal, which determines the overall rate of utilization of physiological resources, as well as on the relative allocation of the processing power to the stimulus (attention selectivity). Similarly, motivation is usually considered to be composed of an arousal component describing the intensity of behaviors or feelings, and a goal-directed component specifying the direction of executed or planned behaviors (Duffy, 1957; Simpson and Balsam, 2016). A more formal definition of motivation has been proposed by Salamone as “the set of processes through which organisms regulate the probability, proximity and availability of stimuli” (Salamone and Correa, 2012). While the caveat of this definition is that it conflates motivation with all aspects of task performance, it also implicitly assumes that motivation includes a directional component oriented towards a goal (altering the probability, proximity, or availability of a stimulus), in agreement with other definitions. Simpson and Balsam proposed an illustrative analogy to describe the link between arousal and motivation. They imagined motivation as a vector whose length represents the intensity of the behavioral pursuit and direction points towards a specific goal (Simpson and Balsam, 2016). In this analogy, arousal corresponds to the vector length. The similarity between the nondirectional (“activational”) components of motivation and affect underscores the importance of the concept of arousal for describing a broad range of likely related but distinct phenomena and hints towards its usefulness in describing the effects of dopaminergic manipulations.

In our rats administered dopamine antagonists, we did not observe a specific impairment of goal-directed behavior but rather a general behavioral inhibition, as evidenced by markedly decreased mobility, both in the presence and in the absence of reward predictive cues. After either SCH or HAL administration, we observed multiple instances of animals staying very close to the receptacle, apparently falling asleep, but clearly trying to respond to the cues and collect their rewards, suggesting that their focus on the goal was not disrupted by the drugs. Therefore, if motivation is composed of arousal and a directional component of behavior as proposed by psychologists (Duffy, 1957), our findings would indicate that dopamine antagonists affected motivation by decreasing arousal. Using the analogy introduced above, the length of the motivation vector decreased while its direction remained largely intact.

Importantly, however, the psychological definition of motivation is not entirely applicable to the case of laboratory animals, or to clinical patients who may suffer from severe ailments that disrupt their very ability to perform actions, rather than their level of arousal or motivation. This possibility must always be considered when dopamine is disrupted due to the ample evidence that dopamine is critical for normal motor function. Our observation that rats treated with dopamine antagonists occasionally froze in the receptacle suggests motor problems, while episodes of extended immobility in sleep-characteristic positions indicate decreased arousal. We have explained above why we believe that our animals treated with dopamine antagonists had motor problems, which were more severe and complex after administration of HAL than SCH. Similar observations and conclusions were made by others but phrased in a different way. Salamone (1988) observed that HAL-administered rats exhibited decreased mobility and motor problems, yet they retained interest in food and performed, or attempted to perform, the activities aimed at obtaining and consuming food. He concluded that dopamine is necessary for the activational, rather than directional, aspect of motivation. While motor deficits were clearly acknowledged, the author argued that the distinction between impaired motor function and motivation is difficult to make. We prefer to refer to the “activational component of motivation” as arousal because, according to most definitions, motivation necessarily has to include a goal-directed component (Duffy, 1957; Simpson and Balsam, 2016). We also believe that the distinction between the animal not being able to perform the task due to motor problems, or not being motivated to perform, is of fundamental importance, even if the answer is difficult to determine in laboratory animals. Of course, if motivation is defined as “the set of processes through which organisms regulate the probability, proximity and availability of stimuli” then any disruption of motor function could be interpreted as a motivational deficit. Yet, if the animal is paralyzed and cannot collect rewards because it is unable to walk, it would be erroneous to conclude that the animal was not motivated.

The identification of multiple physiological correlates of arousal (EEG, skin conductance, muscle tension, and pulse rate) in the first half of the twentieth century contributed to the importance of this concept for bridging psychology and physiology (Duffy, 1957) and inspired definitions of arousal that are based on the physiological reactivity of the subject (Coull, 1998). More recent studies have typically focused on one of the three main varieties of arousal: wakeful, autonomic, or affective (Satpute et al., 2019). Investigations of the neural basis of sleep and wakefulness mostly used EEG desynchronization and decreases in EEG power as measures of wakeful arousal, and attributed a wake-promoting function to the activity of cholinergic cells in the basal forebrain (Han et al., 2014; Irmak and de Lecea, 2014; Xu et al., 2015) (but see Anaclet et al., 2015; Zant et al., 2016), noradrenergic (Berridge et al., 2012), serotonergic (Li et al., 2021; Monti, 2011), and dopaminergic (Cho et al., 2017; Eban-Rothschild et al., 2016) neurons located in the pons and the midbrain, as well as histaminergic (Fujita et al., 2017; Thakkar, 2011) and orexinergic (Sasaki et al., 2011; Tsunematsu et al., 2013; Tyree et al., 2018) hypothalamic neurons. The wake-promoting effects of the activity of these cell groups are antagonized by accumulation of extracellular adenosine (homeostatic sleep drive) and modulated by the oscillations in the activity of neurons in the suprachiasmatic hypothalamic nucleus (circadian regulation) (Brown et al., 2012; Saper et al., 2010). Investigations of autonomic arousal focus on parameters that depend on the activity of sympathetic and parasympathetic systems, such as skin conductance, heart rate, or pupil diameter (Wang et al., 2018). Finally, studies of affective arousal employ self-reports of felt arousal or infer the level of arousal from the degree of salience of presented stimuli (Carr et al., 2020; Deckert et al., 2020; Imbir et al., 2021; Sato et al., 2020).

The diversity of employed methodologies may suggest that these varieties of arousal represent distinct neurophysiological phenomena. However, multiple lines of research indicate that they assess different aspects of the same phenomenon with broad cognitive, behavioral, physiological, and neurological manifestations (Satpute et al., 2019). Accordingly, studies utilizing multiple measures of arousal find that the subjective feeling of arousal and its associated autonomic responses are highly correlated with measures of electrocortical arousal (Lim et al., 1996; Nagai et al., 2004; Waldstein et al., 2000). The brain states corresponding to different arousal levels are set by the activity of cholinergic and monoaminergic neural populations (Lee and Dan, 2012). Most importantly for our study, elevating arousal in rats by handling them was found to increase the release of acetylcholine, norepinephrine, serotonin, histamine, and dopamine, in multiple brain regions and in the periphery (Enrico et al., 1998; Inglis and Fibiger, 1995; Inglis and Moghaddam, 1999; Kalen et al., 1989; Kawahara et al., 1999; Ondicova et al., 2019; Westerink et al., 2002). While all these neuromodulatory systems participate in the natural arousal response, that does not mean that they play the same role. For example, although both central dopamine and norepinephrine are critical for the maintenance of wakeful arousal (Berridge et al., 2012; Carter et al., 2010; Cho et al., 2017; Eban-Rothschild et al., 2016), the firing of LC norepinephrine neurons, but not SNc dopamine neurons, was found to correlate with changes in pupil diameter (Varazzani et al., 2015), consistent with other reports of the involvement of LC norepinephrine neurons in the regulation of autonomic arousal (Larsen and Waters, 2018; Wang et al., 2014). Because there is ample evidence for the control of wakeful arousal by dopamine while its involvement in other arousal varieties has not been directly proven, we referred to wakefulness/drowsiness when discussing the connections between dopamine and arousal throughout the paper. However, manipulations that produce natural behavioral manifestations of drowsiness and sleep, such as those observed in SCH-administered rats, can be expected to affect all aspects of arousal. The natural character of these manifestations and the ability to reverse all of them with a natural arousing stimulus allowed us to conclude that both systemic and intra-NAc SCH exerts its effects on reward seeking by decreasing arousal. The situation is more difficult to interpret in animals that received systemic HAL. Although HAL was found by others to promote sleep, the effects were less potent than those of SCH and restricted to non-REM sleep (Ongini et al., 1993; Trampus and Ongini, 1990). Moreover, HAL and other antipsychotic drugs are known to elevate blood levels of epinephrine and norepinephrine, suggesting that they might *increase* certain aspects of autonomic arousal (Boyda et al., 2020). Finally, as discussed above, HAL increases brain levels of dopamine, likely by antagonizing D2 auto-receptors on dopamine neurons (Blaha and Lane, 1984; Bunney and Grace, 1978; Bunney et al., 1980; Bunney et al., 1973; Imperato and Di Chiara, 1985; Lidsky and Banerjee, 1993), and small doses of HAL are stimulating (Dias et al., 2012). Together with our inability to compensate for the effects of HAL with natural arousing stimuli, these facts suggest that HAL may produce an artificial state that cannot be simply recapitulated as either decreased or increased arousal.

At higher levels of arousal, animals are more likely to exert effort, perform work, remain alert and attentive, and quickly react to environmental signals. While these activities may be critical for survival, performing them exhausts precious resources such as energy and time, and may disrupt physiological homeostatic balance. Therefore, to maximize the chances of survival, the level of arousal must be precisely tuned to the environmental and internal physiological requirements. Functionally, the physiological factors that set the arousal level can thus be thought of as controlling the rate of expenditure of resources. Therefore, a recently proposed hypothesis implicating dopamine in the control of allocation of limited resources (Berke, 2018) is compatible with the role of dopamine as one of the neuromodulators setting the level of arousal. However, describing the function of arousal in terms of resource allocation does not entirely capture its scope. For example, a decrease in wakeful arousal indeed counteracts the expenditure of energetic resources, but it does so by introducing a drive to sleep (Eban-Rothschild et al., 2017), which may be very potent at low arousal levels. Although sleep makes animals vulnerable to predation and is not conducive to foraging and reproduction, its pervasiveness across phylogeny points towards a fundamental function (Anafi et al., 2019). While this function is still debated, multiple lines of evidence indicate that it cannot be reduced simply to saving resources. Behaviorally, sleep can be defined as a state of quiescence during which animals remain immobile in species-dependent sleep-characteristic postures, with significantly reduced responsiveness to mild and moderate stimuli but increased state transition-like responses to strong stimuli (waking) (Campbell and Tobler, 1984). Another defining property of sleep is its homeostatic regulation: sleep deprivation increases the depth and length of sleep that follows when deprivation is discontinued. Using these criteria, sleep was observed across the animal kingdom, including invertebrate organisms such as *Drosophila* and *C. elegans*, and even those lacking cephalized nervous system, such as the jellyfish *Cassiopea* (Anafi et al., 2019; Nath et al., 2017). Not only sleep itself but also its control by dopamine appears to be preserved across phylogeny, as evidenced by the involvement of dopamine in wakeful arousal in *Drosophila* and *C. elegans* (Andretic et al., 2005; Birman, 2005; Foltenyi et al., 2007; Kume et al., 2005; Lebestky et al., 2009; Seugnet et al., 2009; Singh et al., 2014; Van Swinderen and Andretic, 2011; Wu et al., 2008). That sleep is critical for survival, and its function goes beyond saving resources, is underscored by the fact that rats provided with food and water and allowed to rest, but not sleep, develop gross abnormalities in multiple organs within a few days of sleep deprivation, including accumulation of fluid in the lungs, stomach ulcers, and internal hemorrhages. The animals die if sleep deprivation is continued (Rechtschaffen et al., 1983). Sleep is also critical for memory consolidation (Rasch and Born, 2013) and the regulation of the immune system (Bryant et al., 2004). As these functions of sleep clearly go beyond preserving resources, neurophysiological variables that impact the decision whether to sleep or stay awake serve a crucial function that cannot be entirely reduced to resource expenditure. Due to the strong link between dopamine and sleep observed across phylogeny, we think that describing dopamine as one of the arousal variables, rather than reducing its role to resource allocation, provides a more complete picture. Nevertheless, in awake animals, arousal certainly promotes expenditure of resources, whereas motivation and attention allocate them among specific behaviors and mental activities. Importantly, while postulating the involvement of dopamine in resource allocation, Berke concluded that “dopamine provides activational signals—increasing the probability that some decision is made—rather than directional signals specifying how resources should be spent” (Berke, 2018). This description is consistent with the hypothesis implicating dopamine in arousal.

Even in invertebrate organisms, unravelling the mechanism by which dopamine modulates arousal is complicated by the presence of multiple dopaminergic pathways and types of receptors. In agreement with our findings from SCH-administered rats, disruptions of function of D1 receptor orthologs in both *Drosophila* and *C. elegans* were found to enhance sleep (Lebestky et al., 2009; Liu et al., 2012; Singh et al., 2014; Ueno et al., 2012). In flies, strong evidence suggests that wakeful arousal is controlled by D1-like receptors in the dorsal fan-shaped body (Liu et al., 2012; Ueno et al., 2012), although some studies also implicate D1-like receptors in the mushroom body (Driscoll et al., 2020; Sitaraman et al., 2015). In rodents, the first group of dopamine neurons involved in the regulation of sleep was identified in the ventral periaqueductal grey, also referred to as the dorsal raphe nucleus (vPAG/DRN); these cells were active exclusively when rats were awake, and their ablation markedly increased the total time spent asleep (Lu et al., 2006). A recent study found that they are activated in response to arousing stimuli and during waking, and optogenetic and chemogenetic experiments causally linked their activity to arousal (Cho et al., 2017). Dopamine levels in the dorsal striatum were also found to fluctuate across the sleep-wake cycle and increase in response to arousing stimuli (Dong et al., 2019), but the causal involvement of dorsal striatal dopamine in the regulation of wakefulness has not yet been established. Finally, we emphasized in the Introduction the recent evidence for the involvement of VTA dopamine neurons and the NAc in wakeful arousal. Importantly, when the effects of chemogenetic activation of dopamine neurons in the VTA and SNc were directly compared, only the activity of VTA dopamine neurons was effective in promoting wakefulness (Oishi et al., 2017a). Besides the NAc, VTA dopamine neurons innervate other structures. However, when four VTA dopamine neuron terminal regions (NAc, mPFC, CeA, DLS) were directly compared, only stimulation of NAc terminals maintained wakefulness (Eban-Rothschild et al., 2016). Together with recent findings that optogenetic manipulation of NAc medium spiny neurons can robustly and bi-directionally modulate wakeful arousal (Luo et al., 2018; Oishi et al., 2017b), as well as multiple other studies implicating NAc in the control of arousal (Barik and de Beaurepaire, 2005; Lazarus et al., 2011; Qiu et al., 2012; Valencia Garcia and Fort, 2018; Zhang et al., 2013; Zhou et al., 2019), this research suggests that the NAc might be the major region where dopamine exerts its wake-promoting actions. Our finding that intra-NAc administration of SCH results in a within-session decline of performance caused by decreased arousal is consistent with this view. According to the resource allocation hypothesis of dopamine function discussed above, it was proposed that the resource controlled by NAc dopamine is time (Berke, 2018). The hypothesis was motivated by a broad literature showing that inhibition of NAc dopamine function strongly disrupts performance when obtaining rewards requires lengthy operant activities or wait times and it is mostly unaffected when the reward rate is high (Nicola, 2007). As discussed in the Introduction, these results are also expected if NAc dopamine promotes arousal.

We have emphasized the role of dopamine in controlling the level of arousal. However, the symptoms displayed by Parkinson’s patients and multiple experimental results from vertebrate and invertebrate animals support the view that dopamine is also critical for the execution of motor programs (Cermak et al., 2020; Redgrave et al., 2010; Riemensperger et al., 2013; Sharples et al., 2014). While proper motor function certainly requires dopaminergic projections from the SNc to the dorsal striatum and from the hypothalamic A11 dopamine cell group to the spinal cord (Koblinger et al., 2014; Sharples et al., 2014), certain studies indicate that infusions of dopamine receptor antagonists into either the NAc or the dorsal striatum produce comparable degrees of catalepsy (Fletcher and Starr, 1988; Hauber and Munkle, 1997; Ossowska et al., 1990). Our observation that animals occasionally froze in the receptacle, or in awkward positions, indicates that motor problems also contributed to the behavior of our animals. In a typical catalepsy assay, animals’ paws are placed in an uncomfortable position and the time to change this position to a more comfortable one is measured. Interpreting increases in these latencies as impaired motivation is inconsistent with the common understanding of this term, as argued above. While increased arousal can alleviate catalepsy (paradoxical kinesia), and artificial activation of A2A adenosine receptors (which are normally implicated in mediating sleep pressure) can elicit catalepsy (Hauber and Munkle, 1997), catalepsy does not occur when arousal decreases naturally and certainly cannot be reduced to decreased arousal. Therefore, we believe that our findings can be fully accounted for by dual roles for dopamine in both arousal and motor function. We will argue in the next subsection that these two functions of dopamine are sufficient to explain the overwhelming majority of published findings on the effects of dopaminergic disruptions on behavior. In particular, there is little direct evidence for the involvement of dopamine in reward-related functions if one considers the fact that both reward deliveries and expectations are arousing, and collecting rewards involves motor activity. We will refer to this view as the “**arousal-motor hypothesis of dopamine function**”.

### Arousal-motor hypothesis of dopamine function and motivation to exert effort

The hypothesis that implicates mesolimbic dopamine in the motivation to exert effort (Salamone and Correa, 2012) was initially inspired by the critiques of the dopamine hypotheses of reinforcement (Salamone et al., 1997) and learning (Yin et al., 2008), as well as the observation that deleterious effects of NAc dopamine depletions on performance sharply increase with the number of operant responses required (Aberman and Salamone, 1999). Because NAc dopamine disruptions do not impede feeding (Baldo et al., 2002), have little impact on performance of some low-effort operant tasks, and produce much higher sensitivity to the operant requirement than devaluation of food reward by pre-feeding (Aberman and Salamone, 1999), it has been hypothesized that NAc dopamine is important for the exertion of effort to earn rewards, rather than for the primary motivation to obtain them. The effect is most obvious when animals can choose between exerting more effort to earn a preferred reward or less effort to obtain a lesser reward; NAc dopamine depletions induce a bias towards the low-effort/low-reward option (see Salamone et al., 2018 for a review). However, this bias could be due at least in part to motor impairment and/or decreased arousal. Indeed, while extrapyramidal motor symptoms in Parkinson’s patients have been ascribed to dopamine depletion in the dorsal striatum (Di Chiara et al., 1992), some studies suggest that catalepsy and muscle rigidity induced by the systemically administered selective D2 blocker raclopride are predominantly due to its action in the ventral striatum (Alcock et al., 2001; Hemsley and Crocker, 2002). Accordingly, intra-NAc infusions of dopamine antagonists induce catalepsy (Hartgraves and Kelly, 1984; Ossowska et al., 1990) and, at high doses, muscle rigidity (Hemsley and Crocker, 2001). It is therefore conceivable that this and similar problems can steer animals away from the option that requires a higher degree of motor ability. While it has been recognized that NAc dopamine is important for “functions that represent an area of overlap between motor and motivational processes” (Salamone et al., 1999), the relative contributions of motor performance deficits and decreased motivation to exert effort have not been unraveled. Moreover, the potential contribution of drowsiness has been largely unexplored. Notably, an extensive body of research shows that sleep-deprived human subjects perceive their tasks as more difficult and this can “lead to decisions to work on easier tasks” (Engle-Friedman, 2014). Another independent mechanism that can promote increased consumption of less-preferred rewards by drowsy animals is that of compromised attention selectivity; in multitasking subjects, arousal improves performance of the primary task at the expense of less important options, whereas drowsiness has the opposite effect (Hockey, 1970). If dopamine-compromised animals are drowsy, both decreased willingness to exert effort and compromised attention selectivity could contribute to the apparent shift of preference towards the low-effort/low-reward option. It is therefore possible that the effects of NAc dopamine manipulations on effort-based decision making are entirely due to compromised motor function and decreased arousal, and the apparent decrease in motivation is an epiphenomenon.

Some support for the contribution of drowsiness to the behavioral results that inspired the motivation-effort hypothesis of mesolimbic dopamine come from the studies on the mechanism of action of the natural somnogen adenosine (Lazarus et al., 2019). Enhanced drive to sleep after periods of prolonged wakefulness has been linked to the accumulation of adenosine in the basal forebrain (Porkka-Heiskanen et al., 2000). Consistently, intraventricular infusions of an adenosine A2A receptor agonist CGS21680 showed that infusion into the ventricle closest to the NAc had the strongest sleep-promoting effects of this compound (Satoh et al., 1999). Interestingly, intra-NAc administrations of CGS21680 into rats performing operant schedules for food reward caused very similar effects to those of dopamine depletions, including preferential inhibition of performance of high ratio schedules (Mingote et al., 2008a) and a bias towards a less-effortful option in concurrent schedules (Font et al., 2008). While these findings were interpreted as evidence for the involvement of NAc A2A receptors in the control of motivation and effort, the contribution of drowsiness, likely induced by intra-NAc administration of this somnogen (Satoh et al., 1999), was not convincingly ruled out. In fact, systemic administrations of CGS21680 were found to inhibit operant performance mainly due to drowsiness (Mingote et al., 2008b) and, because the sleep-promoting effects of this compound were attributed to its action in the NAc (Satoh et al., 1999), it would be surprising if intra-NAc administrations inhibited motivation to exert effort via a mechanism that does not involve drowsiness. The main argument against the involvement of drowsiness, used by the proponents of the motivation/effort hypothesis, is that while intra-NAc administrations decrease lever pressing for preferred food pellets, they also increase consumption of chow that is freely available on the floor – an effect that is distinct from those of systemic administrations (Font et al., 2008). However, this outcome is very much compatible with the induction of sleepiness in severely food-restricted rats; animals that are both hungry and sleepy are likely to choose to appease their hunger with minimal effort. That systemic CGS21680 inhibits lever pressing without enhancement of chow intake could be due to an additional impairment of the ability to handle food, or appetite-suppressant effects (Micioni Di Bonaventura et al., 2012), caused by the action of the drug in other brain regions. Furthermore, the fact that the experimenters did not observe “any overt signs of sedation in rats that received intra-accumbens injections of CGS 21680” might be due to the compensatory influence of heightened arousal caused by the transfer of the animals to the operant chamber. With this regard, we emphasize that we did observe overt signs of sedation after intra-NAc administration of SCH (Supplemental Video 2), but these signs were not as evident during the initial 30 min. Because effort-based choice experiments usually involve 30 min operant sessions, it is possible that the effect of initial arousal could prevent the development of overt signs of sedation.

### Arousal-motor hypothesis of dopamine function and wanting/incentive salience

An influential hypothesis implicates dopamine in wanting rewards and attributing “incentive salience” to reward-predictive cues (Berridge, 2007). This hypothesis is based on the premise that disruptions of dopamine impair pursuit of rewards due to interference with one of the three possible reward-related functions: liking, learning, or wanting. The arguments against the involvement of dopamine in liking and learning are used to support its role in wanting. However, if the main function of dopamine is to maintain arousal and motor capability necessary for performance, the experiments testing its involvement in liking and learning would give negative results as well. In fact, to our knowledge no experiment in which dopamine function was reduced has implicated dopamine in wanting in a way that is not also compatible with a reduction in arousal and motor function. For example, in the general form of Pavlovian-instrumental transfer (PIT), presentation of a conditioned stimulus (CS) increases operant responding even in cases where the reward associated with the CS is different from that associated with the operandum. Interference with NAc dopamine reduces the general PIT effect (but not the “specific” PIT effect in which CSs specifically potentiate responding on an operandum associated with the same outcome that is also associated with the CS, and not an operandum that earns a different outcome) (Corbit and Balleine, 2016; Nicola, 2016). These results have been interpreted as evidence for a role for mesolimbic dopamine in representing the value of external stimuli and internal states (Dayan and Balleine, 2002; Yin et al., 2008), which is similar (if not identical) to the idea that mesolimbic dopamine heightens incentive salience. However, the PIT results are equally compatible with roles for dopamine in arousal and motor function – i.e., presentation of a CS increases arousal, resulting in increased probability of any behavior (aside from sleep preparation), including seeking rewards that are not explicitly associated with the CS. The arousal-motor hypothesis therefore predicts that events that activate reward-seeking behavior need not themselves carry motivational significance, whereas the incentive salience hypothesis posits that only events that increase motivation should have such an effect. In our experiments, brief mid-session handling had no particular motivational value, yet it strongly activated performance in SCH-treated animals. Notably, this result occurred in the presence of a D1 receptor antagonist, yet blockade of D1 receptors has been shown to reduce the general PIT effect (Lex and Hauber, 2008), superficially suggesting that mid-session handling and the CS presented in PIT tests activate performance via different neural mechanisms. However, if the CSs presented in the PIT test are not strongly arousing (e.g., they are similar in their ability to promote arousal to the DSs we present here), D1 receptor blockade would be expected to reduce their ability to promote behavior during the long PIT test session just like SCH reduced the ability of DSs (but not handling, a more strongly arousing stimulus) to promote behavior. To further test between incentive salience and arousal-motor interpretations, it would be necessary to assess whether non-motivational arousal-promoting manipulations can, just like CSs, activate operant performance in extinction (i.e., in a situation similar to the PIT test session) and also in the DS task under conditions in which animals’ performance is reduced by a natural low state of arousal (e.g., after sleep deprivation).

Another example of the incentive salience interpretation drawn from the behavioral results of dopaminergic disruptions comes from the study of genetically engineered dopamine-deficient mice in a T-maze reward seeking task (Robinson et al., 2005). These mice were hypoactive, they did not eat or drink on their own, and died within 4 weeks after birth unless given L-dopa injections (Zhou and Palmiter, 1995). When taken off L-dopa, dopamine-deficient mice were unable to explore the T-maze due to severe hypoactivity. The authors found that CAF ameliorated the hypoactive phenotype and allowed the mice to explore the maze, consume rewards, and learn the task. Still, CAF-treated dopamine-deficient mice had longer latencies to reach the T-maze intersection and to consume the reward after the appropriate arm was entered. The longer latencies might have been due to residual performance problems that were not eliminated by CAF, or due to new performance problems induced by a very high dose of CAF (25 mg/kg, compared to 10 mg/kg used in our study). However, the authors chose to interpret them as an indication of impaired wanting. As an argument in favor of this interpretation, they noticed that when CAF-treated mice were switched to L-dopa, the latency to consume rewards remained initially elongated. They argued that if mice wanted rewards but were unable to consume them quickly due to performance problems, then the delay should disappear when mice were given L-dopa and performance problems were eliminated. However, if the delay were due to decreased wanting caused by the lack of dopamine, then administration of L-dopa should also have instantly eliminated it. Therefore, using this argument to favor a wanting deficit over performance problems is unfounded.

Despite these issues, the idea that CAF can alleviate performance deficits in dopamine-compromised animals without affecting reward-specific problems is intriguing and motivated us to examine the effects of CAF on our SCH-and HAL-treated rats. After CAF treatment, our animals responded to almost all reward-predictive cues and consumed the rewards, showing no indication of impaired wanting. It has been previously reported that CAF, as well as specific antagonists of A2A (but not A1) adenosine receptors, abolishes the inhibitory effects of D2 antagonists on the pursuit of rewards in diverse operant tasks (Farrar et al., 2007; Mott et al., 2009; Salamone et al., 2009), and partially alleviates the effects of D1 antagonists (Worden et al., 2009). Our findings show that the interactions between the activity of dopamine and adenosine receptors similarly control responding to the reward-predictive cues, and that these cues retain incentive salience and motivate approach in dopamine-deficient animals as long as their performance problems are compensated for. Thus, both our results and previous reports of the behavioral effects of dopamine disruption can be explained by reductions in arousal and motor ability rather than a role for dopamine in attribution of incentive salience.

### Arousal-motor hypothesis and ramping dopamine signals

An interpretation of dopamine signals as temporal difference reward prediction error (TD RPE) signals (Schultz, 1998; Schultz et al., 1997; Schultz et al., 2017) has recently been challenged by the observation of ramping dopamine signals preceding reward receipt in animals performing certain operant tasks (Hamid et al., 2016; Howe et al., 2013; Mohebi et al., 2019). These increases in dopamine were observed in well-trained animals even during periods when no unexpected rewards or stimuli were delivered and thus it was argued that they cannot represent a prediction error (Hamid et al., 2016; Howe et al., 2013; Mohebi et al., 2019; Niv, 2013). It was proposed that these dopamine signals may represent future reward value discounted by time (Hamid et al., 2016). However, this interpretation was subsequently questioned by a study in which mice moved along a virtual alley to obtain a reward at a designated location (Kim et al., 2020). The authors observed ramping dopamine signals resembling those reported by others, but when they unexpectedly advanced spatial cues so that the animals were teleported in the virtual alley towards the reward delivery location, they observed sudden increases in dopamine that were greater in magnitude than the discounted value interpretation would predict. When the progression in the alley was paused or reversed, dopamine signals that had started ramping up gradually declined to the baseline level. Based on these findings, the authors argued that ramping dopamine signals represent the TD RPE. When mice were teleported to an equivalent position in an alternative virtual alley that utilized different but functionally equivalent visual cues, no significant increase in dopamine was observed.

While the discussions concerning whether ramping dopamine signals are better described in terms of discounted value or TD RPE are ongoing, the results recapitulated above suggest that neither of these simple interpretations alone can entirely explain the ramping phenomenon. In fact, the results observed so far in experiments that have revealed and manipulated dopamine ramp signals are precisely what one would expect if dopamine represents the level of arousal. For example, an approaching salient event, whether it is a forthcoming reward or an impending punishment, should gradually increase arousal; sudden advancement of the temporal progression could conceivably increase arousal above the baseline ramping trend; and signaling to the animal that the salient stimulus is delayed could decrease the level of arousal back to baseline. Finally, while the absence of a dopamine increase after teleportation to an equivalent position in another virtual alley was used as an argument that dopamine does not represent “sensory arousal” (Kim et al., 2020), this experiment cannot be used to rule out the arousal interpretation because salient events and their predictors are presumably much more arousing than meaningless sensory alterations.

### Arousal-motor hypothesis and dopamine signaling theories

Recordings from dopamine neurons and measurements of dopamine release in the NAc and striatum in response to specific events have formed the basis for a variety of hypotheses regarding the nature of dopamine signaling. The most prominent of these is that dopamine conveys TD RPE signals, which have been proposed to contribute to reinforcement learning (Schultz et al., 1997; Schultz et al., 2017). Although a specific role for dopamine in reinforcement learning seems to be nominally inconsistent with the arousal-motor hypothesis, the two hypotheses should be considered in light of extensive evidence for other forms of information encoding by dopamine neurons. Different subsets of these neurons are responsive to sensory salience (e.g., how loud or “alerting” an auditory stimulus is), motivational salience (the degree to which a stimulus predicts positive and negative value), aversive events and their predictors, and RPEs (Bromberg-Martin et al., 2010). Notably, although the specific behavioral (approach, avoidance, orienting) and cognitive (learning, paying attention) responses appropriate to each of these signals differ, they have in common that all of the stimuli that elicit these various dopamine responses are arousing. Moreover, these stimuli almost always trigger motor responses whose characteristics depend on the meaning of the stimulus, suggesting that dopamine signals may encode motor responses. This proposal could explain much of the observed diversity in dopamine signaling. For example, events ranging in aversiveness from failure to receive a predicted reward to receiving an electric shock have been shown to cause decreases in the dopamine signal (Badrinarayan et al., 2012; Brischoux et al., 2009; de Jong et al., 2019; Hart et al., 2014; Lerner et al., 2015; Menegas et al., 2018; Mirenowicz and Schultz, 1996; Yuan et al., 2019). Although declines in dopamine in response to strongly aversive stimuli are nominally inconsistent with a role for dopamine in arousal, recent studies have consistently identified specific subregions in which aversive events and/or their predictors elevate the dopamine signal (Badrinarayan et al., 2012; Brischoux et al., 2009; de Jong et al., 2019; Lerner et al., 2015; Menegas et al., 2018; Yuan et al., 2019), and these elevations could drive increased arousal. Moreover, the specific nature of the local dopamine response may serve to facilitate specific behaviors in response to the stimulus, such as freezing vs escape.

Further evidence supports the possibility that dopamine neurons encode aspects of motor behavior. Some dopamine neurons increase activity before the onset of spontaneous movements, and their optogenetic activation can trigger movements (Barter et al., 2015; da Silva et al., 2018; Howe and Dombeck, 2016). Rather than indiscriminately invigorating movement, some dopamine neurons were reported to encode specific components of the velocity vector (Barter et al., 2015), as well as the initiation, termination, and duration of lever presses (Fan et al., 2012). While most studies implicating dopamine neurons in the control of movement were performed in the SNc, it has been recently reported that activity of VTA dopamine neurons in head-restrained mice collecting rewards, a setup that has been extensively used to study RPE encoding, correlates with motor actions such as initiation of licking (Coddington and Dudman, 2018) or force impulse (Hughes et al., 2020). Coddington and Dudman (2018) suggested that all aspects of RPE encoding can be recapitulated by a model that takes into account exclusively responses of dopamine neurons to reward-predictive cues (which become increasingly arousing as the animal learns their meaning) and the encoding of reward-related motor actions (brief excitations preceding initiation of licking followed by inhibitions after the onset of licking); the dependence of timing of these events on the stage of learning has to be taken into account. Recently, utilizing a head fixation apparatus incorporating force sensors in mice performing a fixed interval task, Hughes et al. (2020) found that VTA dopamine neurons can be segregated into those that are excited by forward movements and inhibited by backward movements, and those that are excited by backward movements and inhibited by forward movements. While most of these neurons were excited around the time of reward delivery, the authors found that mice generated characteristic movement patterns when the rewards were delivered and consumed, and that the reward-related activity can be entirely explained by these movements. More precisely, the number of spikes generated within each movement was linearly correlated with the integral of force over the time period of movement occurrence - the force impulse (the time window for the spike count was advanced in time to account for the fact that neural activity preceded movement onset). These correlations looked the same when movements during the ITIs or outside the reward context were analyzed. The force impulse is proportional to the change of velocity in unrestrained animals. Encoding of velocity by dopamine neurons is consistent with our previous finding in rats performing a cued sucrose seeking task that cue-evoked excitations in the NAc, which are inhibited by ipsilateral but not contralateral dopamine antagonist infusions and thus directly depend on dopamine (du Hoffmann and Nicola, 2014), strongly correlate with the speed of the ensuing approach (McGinty et al., 2013).

These findings show that the RPE encoding by dopamine neurons can be at least partially explained by the encoding of movement. Moreover, selective correlation of the activity of different dopamine neuronal populations with distinct movements at the sub-second time scale cannot be thought of as secondary to their role in motivation or arousal, and, therefore, the control of movement must be a part of any comprehensive theory of dopamine function. Conversely, the involvement of dopamine in the control of arousal cannot be entirely explained by its role in motor function. While the inhibition of movement could conceivably contribute to drowsiness, the experiments in which optogenetic stimulation of dopamine neurons or activation of D1 receptors during sleep (Eban-Rothschild et al., 2016) or anesthesia (Taylor et al., 2013; Taylor et al., 2016) induced signs of electrocortical arousal followed by transition to wakefulness cannot be explained by movement-enhancing effect of the stimulation because animals did not move under these conditions. Moreover, inhibition of dopamine function does not result in the cessation of all movements followed by sleep, but usually increases “motivation to sleep” (Eban-Rothschild et al., 2017) as evidenced by preparatory activities whose specific nature may depend on the experimental setup. For example, our SCH-administered rats sought comfortable positions, and mice provided with bedding engaged in nest building when their VTA dopamine neurons were inhibited (Eban-Rothschild et al., 2016), suggesting that increased drowsiness occurred before the ability to move was severely disrupted. Finally, taking the role of dopamine in arousal into account is indispensable to explain the within-session declines of performance we observed in our SCH-administered rats, and it may also be necessary to correctly interpret within-session declines observed after dopaminergic disruptions by others. Indeed, the assumption that performance problems induced by dopamine antagonists should exclusively reflect motor impairments or physical fatigue led to the hypothesis that within-session declines cannot be explained by performance problems, leaving impaired reinforcement as the only viable possibility (Fouriezos and Wise, 1976; Franklin and McCoy, 1979; Wise, 2004). However, reduced arousal must also be considered as a potential cause of such declines; we discuss the reasons for this in detail in the Introduction.

Disruptions in performance due to decreased arousal or motor deficits could also explain observations that interference with dopamine transmission impairs learning. For example, systemic dopamine receptor blockade prevents the emergence of sign tracking but not goal tracking (Flagel et al., 2011). However, the antagonist also impaired performance during acquisition (consistent with an arousal and/or motor deficit), and if learning to sign track simply required a greater number of successful trials than learning to goal track, this could have caused the apparently specific deficit in sign tracking acquisition. Similarly, a recent study demonstrated that optogenetically inhibiting dopamine neurons specifically during reward delivery caused a run-down in licking responses to subsequent cues, suggesting a role for dopamine released during reward consumption in maintaining this behavior (Lee et al., 2020). However, reducing dopamine neuronal activity just when it is highest could have caused a longer-term reduction in downstream dopamine levels, resulting in reduced arousal and therefore reduced responding. Indeed, even when two cues were presented in interleaved trials, and dopamine neurons were inhibited during rewards delivered after only one of them, the greatest effect appeared to be a run-down in responding to both cues, as predicted by the arousal hypothesis (although the run-down was slightly greater for the cue that was paired with dopamine neuron inhibition) (Lee et al., 2020). Although it remains, in theory, possible that transient dopamine activations by rewards or other events participate in learning via a RPE or other mechanism, it is important to consider whether the observations that support a role in learning could be explained by a role for dopamine in arousal or motor performance.

### Summary and conclusions

Because the involvement of dopamine in both motor control and arousal is sufficient to explain our results without the need to assume that dopamine neurons engage in reward-related signaling, our findings are most parsimoniously explained by the arousal-motor hypothesis of dopamine function, as opposed to other general hypotheses that aim to explain the effects of nonspecific dopaminergic disruptions on behavior by decreased reinforcement, wanting, motivation, or willingness to exert effort. Of course, none of these simple and general hypotheses, including the arousal motor hypothesis, can account for the activity of each individual dopamine neuron, especially if neural activity is analyzed at high temporal resolution. However, as most of the stimuli that excite dopamine neurons are arousing and the activity of most of these neurons is implicated in motor control, a decrease in arousal and impaired motor function may be the primary manifestation of an indiscriminate disruption of dopaminergic function that targets all these functionally distinct populations. For this reason, the arousal-motor hypothesis has important implications for the interpretation of studies in which dopamine function is disrupted. For example, we and others have observed that dopamine antagonists reduce animals’ responding to reward-predictive cues in various behavioral paradigms (Di Ciano et al., 2001; Lex and Hauber, 2008; Nicola, 2010; Ogren and Fuxe, 1988; Saunders and Robinson, 2012; Wakabayashi et al., 2004; Wenzel et al., 2018; Yun et al., 2004). These effects may have been amplified by reduced arousal, especially if operant sessions and ITIs are long. In theory, brief and temporally specific manipulations of dopamine using optogenetics should produce results less susceptible to this confound; however, it is possible that even brief reductions in dopamine function could transiently produce effects that are normally associated with drowsiness, such as decreased attention, which could decrease the likelihood of a behavioral response to cues.

## Abbreviations

CAF: caffeine
CS: conditioned stimulus
DS: discriminative stimulus
HAL: haloperidol
ICSS: intracranial self-stimulation
IP: intraperitoneal
ITI: intertrial interval
NAc: nucleus accumbens
NS: neutral stimulus
PIT: Pavlovian-instrumental transfer
SC: subcutaneous
SCH: SCH23390
TD RPE: temporal difference reward prediction error
VTA: ventral tegmental area

## Acknowledgments

This work was supported by NIH grants DA019473, DA038412, and DA044761. We thank Leah Linfield for technical assistance.

