## Supplemental Figures, Table and Legends for "The arousal-motor hypothesis of dopamine function: evidence that dopamine facilitates reward seeking in part by maintaining arousal"

**Supplemental Videos**

**Supplemental Video 1.** Mid-session handling procedure in a rat given an intra-NAc injection of SCH23390. The animal is in a quiescent state prior to handling but alert and mobile during and after handling. Auditory cues were normally presented; for video analysis, they were accompanied by a light.

**Supplemental Video 2.** The curled-up position. This position is a typical sleep posture, yet the animal is able to respond to a DS presentation (which, unlike in Supplementary Video 1, is not shown by a light).

| Experiment | r | t | P |
|---|---|---|---|
| VEH | 0.087 | 0.275 | 0.789 |
| VEH 1 h op. box wait | 0.313 | 1.042 | 0.322 |
| SCH 1 h op. box wait | 0.822 | 4.571 | **0.001** |
| SCH + VEH | 0.818 | 4.502 | **0.001** |
| SCH 60 cohort 2 | 0.802 | 4.245 | **0.002** |
| SCH + CAF | 0.397 | 1.368 | 0.201 |
| HAL + VEH | 0.443 | 1.563 | 0.149 |
| HAL 100 cohort 2 | -0.236 | -0.767 | 0.461 |
| HAL + CAF | 0.195 | 0.630 | 0.543 |

**Table S1. Statistical results of Pearson's correlations shown in Fig. 9E.** For each experiment, the r value for the correlation between mobility and DS response ratio is shown, as well as the corresponding t and P values. Bold values indicate significant correlations ($P < 0.05$).

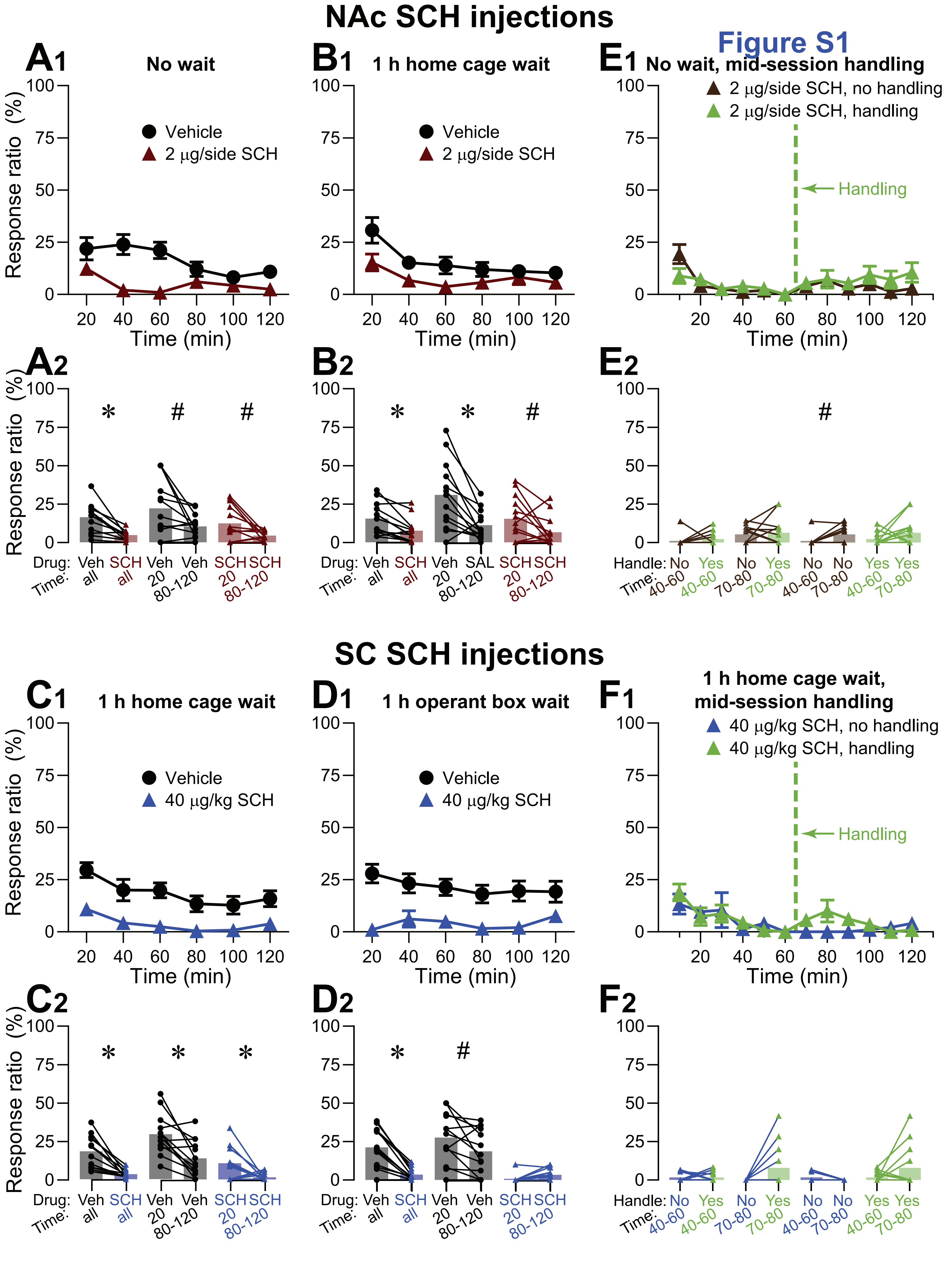
NAc SCH injections
Figure S1
A1 No wait
B1 1 h home cage wait
E1 No wait, mid-session handling
Vehicle
2 μg/side SCH
2 μg/side SCH, no handling
2 μg/side SCH, handling
Handling
Response ratio (%)
Time (min)
A2
B2
E2
Drug: Veh SCH Veh Veh SCH SCH
Drug: Veh SCH Veh SAL SCH SCH
Time: all all 20 80-120 20 80-120
Handle: No Yes No Yes No No Yes Yes
Time: 40-60 40-60 70-80 70-80 40-60 70-80 40-60 70-80
SC SCH injections
C1 1 h home cage wait
D1 1 h operant box wait
F1 1 h home cage wait, mid-session handling
40 μg/kg SCH
40 μg/kg SCH, no handling
40 μg/kg SCH, handling
C2
D2
F2

**Figure S1. NS response ratios after intra-NAc and systemic SCH23390 injections.** Following a similar format to Fig. 2 (with similar graphing conventions), the graphs show the NS response ratio for the corresponding experiments in Fig. 2. Although the NS response ratio is considerably lower than the DS response ratio, the effects of SCH microinjected in the NAc (A, B, E) or systemically (C, D, F) were similar to the effects on the DS response ratio (Fig. 2), although mid-session handling did not increase the NS response ratio (E, F). Symbols above the connecting lines show the results of paired comparisons (two-tailed permutation test) between the points connected by the lines: ∗, $P < 0.05$, adjusted for multiple comparisons; #, $P < 0.05$, but the result did not reach significance after adjusting the critical P value for multiple comparisons; no symbol, $P > 0.05$.

### Figure S2

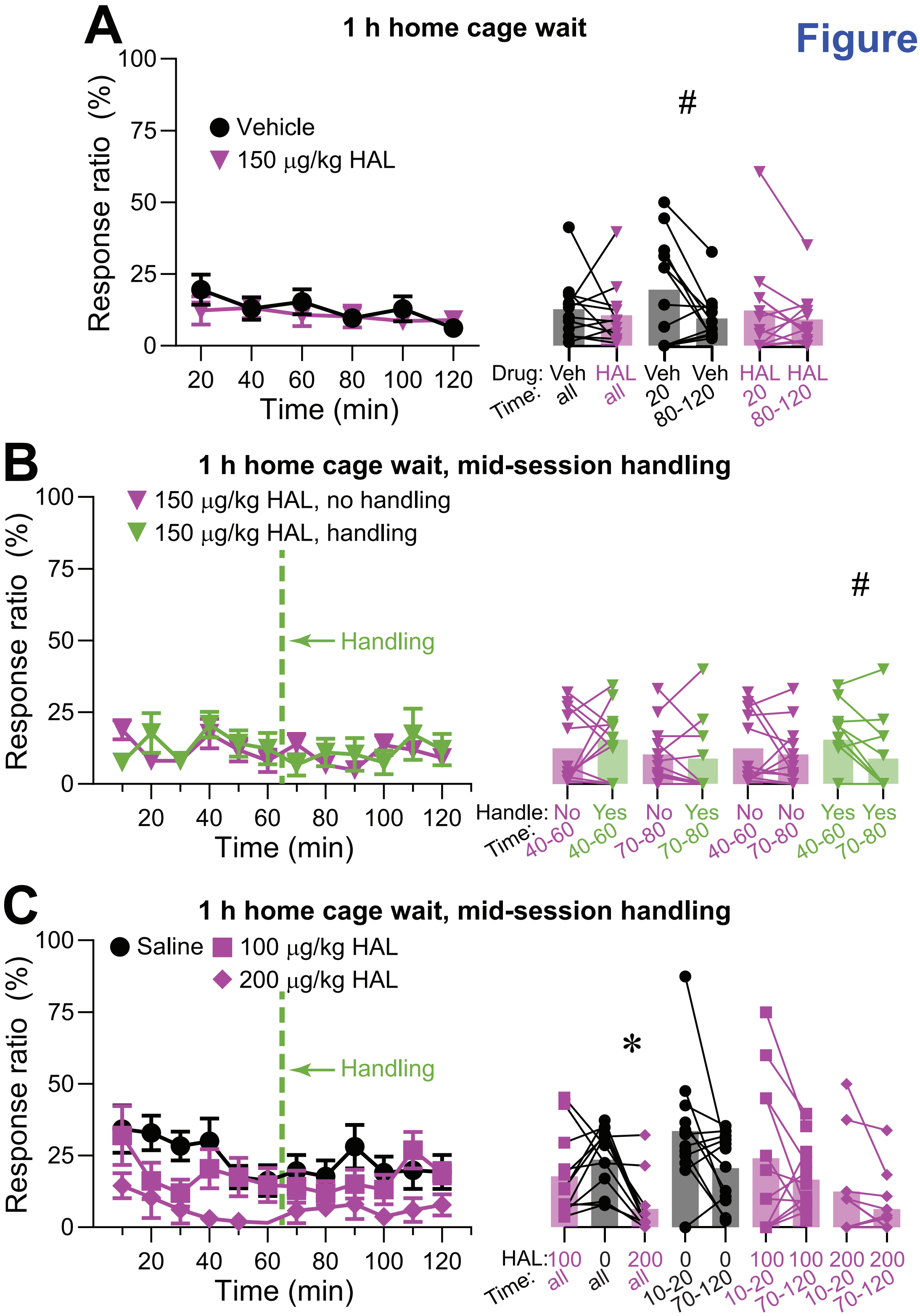
A
1 h home cage wait
Vehicle
150 µg/kg HAL
Response ratio (%)
Time (min)
#
Drug: Veh HAL Veh Veh HAL HAL
Time: all all 20 80-120 20 80-120
B
1 h home cage wait, mid-session handling
150 µg/kg HAL, no handling
150 µg/kg HAL, handling
Handling
Response ratio (%)
Time (min)
#
Handle: No Yes No Yes No No Yes Yes
Time: 40-60 40-60 70-80 70-80 40-60 70-80 40-60 70-80
C
1 h home cage wait, mid-session handling
Saline
100 µg/kg HAL
200 µg/kg HAL
Handling
Response ratio (%)
Time (min)
*
HAL: 100 0 200 0 0 100 100 200 200
Time: all all all 10-20 70-120 10-20 70-120 10-20 70-120

**Figure S2. NS response ratios after systemic SCH23390 injections.** Following a similar format to Fig. 4 (with similar graphing conventions), the graphs show the NS response ratio for the corresponding experiments in Fig. 4. Symbols above the connecting lines show the results of paired comparisons (two-tailed permutation test) between the points connected by the lines: ∗, $P < 0.05$, adjusted for multiple comparisons; #, $P < 0.05$, but the result did not reach significance after adjusting the critical P value for multiple comparisons; no symbol, $P > 0.05$.

Figure S3

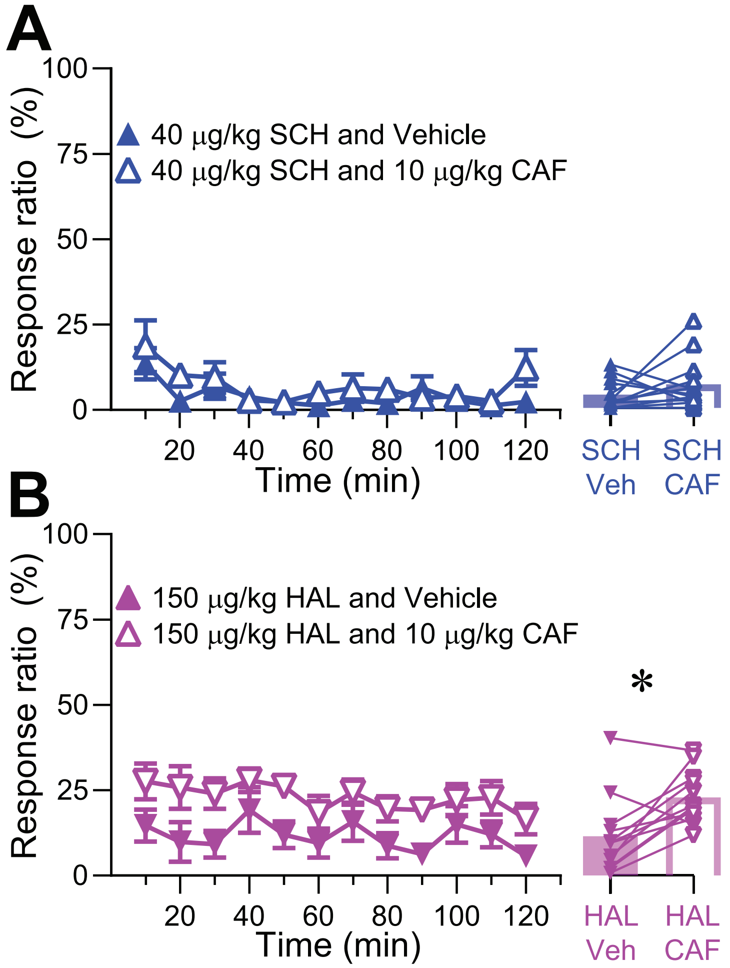
A
Response ratio (%)
100
75
50
25
0
40 µg/kg SCH and Vehicle
40 µg/kg SCH and 10 µg/kg CAF
20
40
60
80
100
120
Time (min)
SCH Veh
SCH CAF
B
Response ratio (%)
150 µg/kg HAL and Vehicle
150 µg/kg HAL and 10 µg/kg CAF
*
Time (min)
HAL Veh
HAL CAF

**Figure S3. NS response ratios during caffeine experiments.** Following a similar format to Fig. 5 (with similar graphing conventions), the graphs show the NS response ratio for the corresponding experiments in Fig. 5. Symbols above the connecting lines show the results of paired comparisons (two-tailed permutation test) between the points connected by the lines: *, $P < 0.05$, two-tailed permutation test; no symbol, $P > 0.05$.

### Figure S4

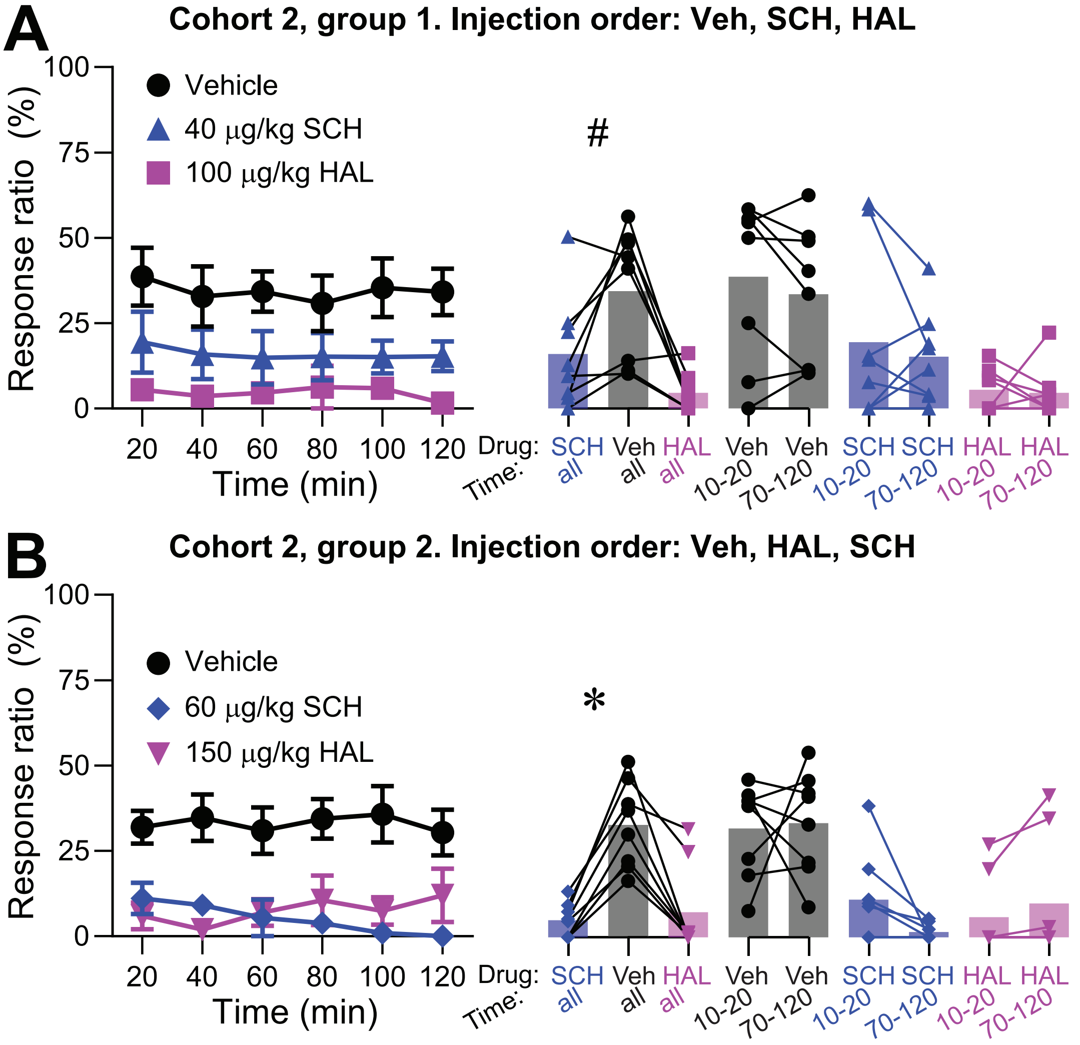
A
Cohort 2, group 1. Injection order: Veh, SCH, HAL
Response ratio (%)
Vehicle
40 μg/kg SCH
100 μg/kg HAL
#
Time (min)
Drug:
Time:
SCH Veh HAL Veh Veh SCH SCH HAL HAL
all all all 10-20 70-120 10-20 70-120 10-20 70-120
B
Cohort 2, group 2. Injection order: Veh, HAL, SCH
Response ratio (%)
Vehicle
60 μg/kg SCH
150 μg/kg HAL
*
Time (min)
Drug:
Time:
SCH Veh HAL Veh Veh SCH SCH HAL HAL
all all all 10-20 70-120 10-20 70-120 10-20 70-120

**Figure S4. NS response ratios for cohort 2.** Following a similar format to Fig. 6 (with similar graphing conventions), the graphs show the NS response ratio for the corresponding experiments in Fig. 6. #, $P < 0.05$, but the result did not reach significance after adjusting the critical P value for multiple comparisons; no symbol, $P > 0.05$; two-tailed permutation tests.

Figure S5

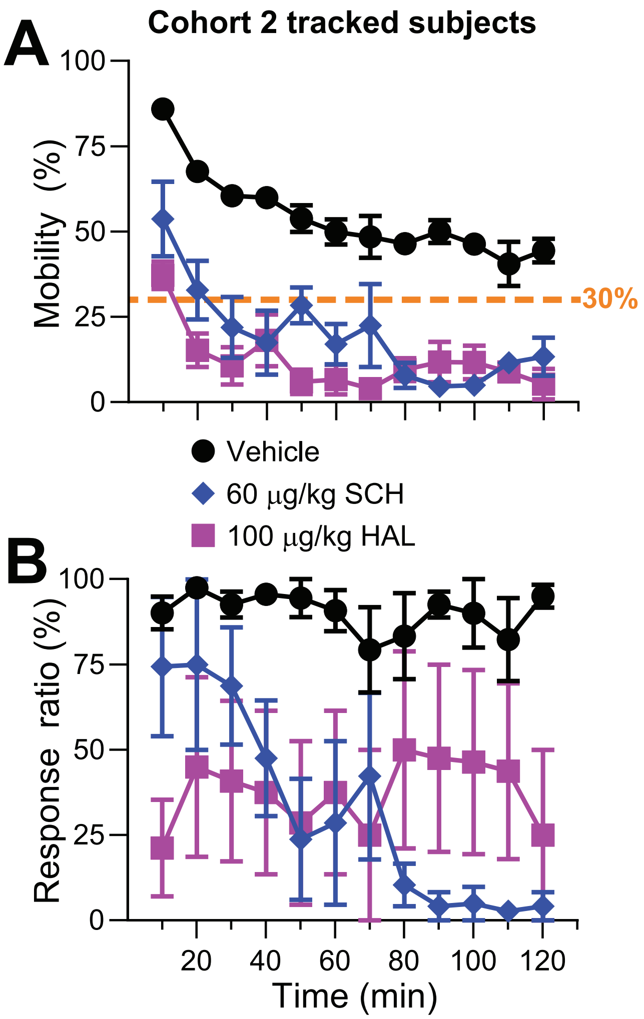
A
Cohort 2 tracked subjects
Mobility (%)
100
75
50
25
0
30%
Vehicle
60 µg/kg SCH
100 µg/kg HAL
B
Response ratio (%)
100
75
50
25
0
20
40
60
80
100
120
Time (min)

**Figure S5. Impact of SCH23390 and haloperidol on the mobility of rats in cohort 2.** The graphs show data for the subset of subjects used for the experiment shown in Fig. 6 for which video tracking data was available. A shows mobility (percent of time spent in motion within the indicated time bin) whereas B shows DS response ratio in the same bins. Dashed horizontal line in A indicates 30% mobility. Statistical tests were not performed because the N of 4 subjects per group was too low to yield interpretable permutation test results (see Methods, Statistical Analysis).

### Cohort 2

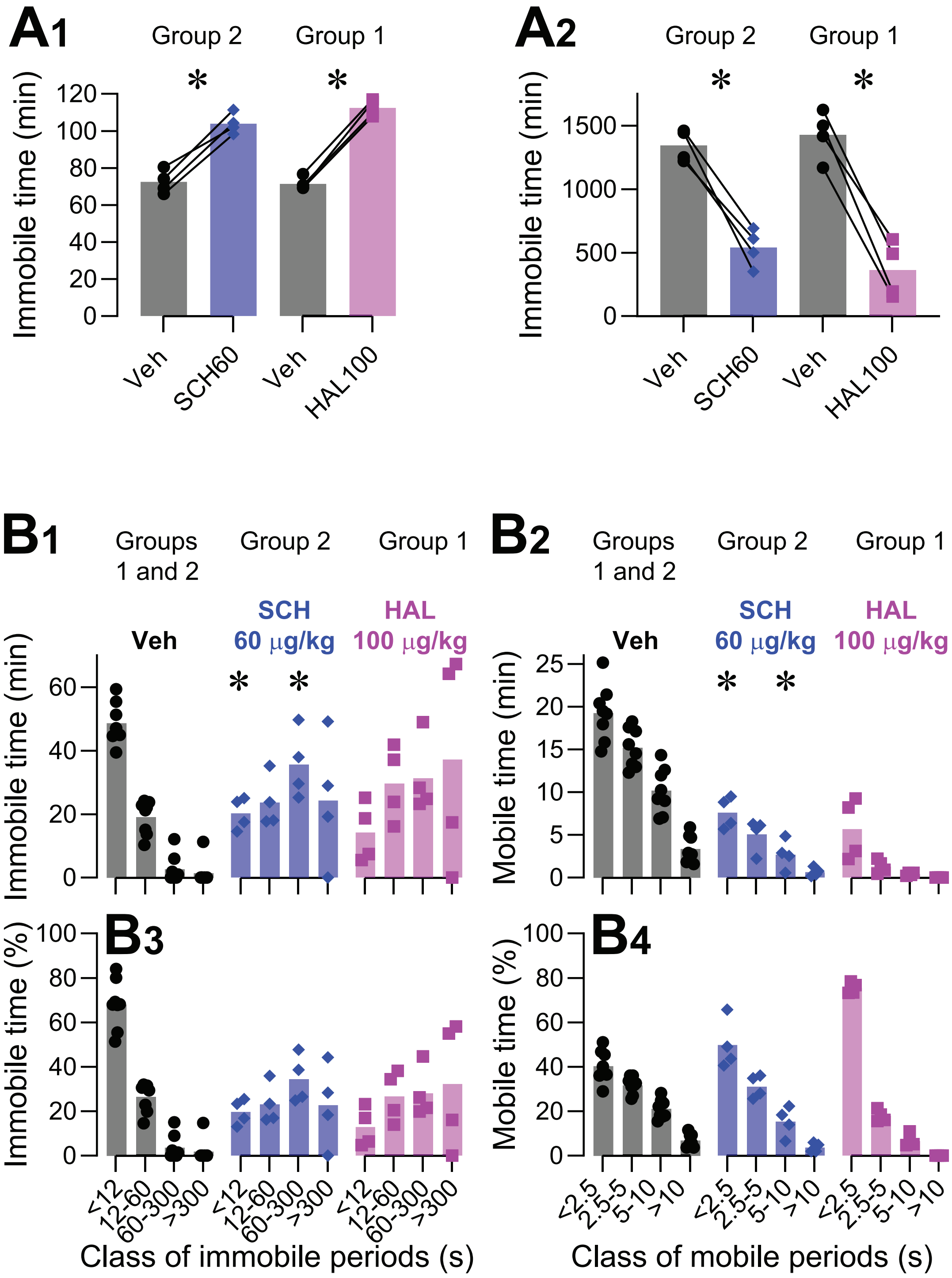
A1
Group 2
Group 1
*
*
Immobile time (min)
Veh
SCH60
Veh
HAL100
A2
Group 2
Group 1
*
*
Immobile time (min)
Veh
SCH60
Veh
HAL100
B1
Groups 1 and 2
Group 2
Group 1
Veh
SCH 60 μg/kg
HAL 100 μg/kg
Immobile time (min)
B2
Groups 1 and 2
Group 2
Group 1
Veh
SCH 60 μg/kg
HAL 100 μg/kg
Mobile time (min)
B3
Immobile time (%)
B4
Mobile time (%)
<12
12-60
60-300
>300
Class of immobile periods (s)
<2.5
2.5-5
5-10
>10
Class of mobile periods (s)

**Figure S6. Impact of dopamine antagonists on the distribution of mobility and immobility periods for cohort 2.** The data are taken from the cohort 2 experiment shown in Fig. 6, limited to the subset of rats (N=8) for which video tracking was performed. These consist of 4 rats in group 1 that received vehicle and 60 µg/kg SCH, and 4 rats in group 2 that received vehicle and 100 µg/kg HAL. A, Total immobile time (A1) and the number of mobility periods (A2) registered throughout the entire operant session after administration of the indicated drugs. B,C, Total time (B1, B2), or percent of total time (B3, B4), spent in immobility (B1, B3) or mobility (B2, B4) periods belonging to the designated length classes indicated on the abscissa, after administration of the indicated drugs. Vehicle data includes values from both groups 1 and 2 (N = 8), whereas drug data includes values only from a single group (N = 4). A1,A2,B1,B2, Significance symbols indicate the results of paired t-tests vs vehicle. ∗, $P < 0.05$, remaining significant after adjustment for multiple comparisons; no symbol, $P > 0.05$. B3,B4, Statistical tests were not performed due to the low N.
